# Phylogeny, evolution, and classification of the ant genus *Lasius*, the tribe Lasiini, and the subfamily Formicinae (Hymenoptera: Formicidae)

**DOI:** 10.1101/2021.07.14.452383

**Authors:** B. E. Boudinot, M. L. Borowiec, M. M. Prebus

**Affiliations:** Department of Entomology & Nematology, University of California, Davis CA; Friedrich-Schiller-Universität Jena, Institut für Spezielle Zoologie, Jena, Germany; Department of Plant Pathology, Entomology and Nematology, University of Idaho, Moscow ID; Institute for Bioinformatics and Evolutionary Studies, University of Idaho, Moscow ID; School of Life Sciences, Arizona State University, Tempe AZ

**Keywords:** Integrated taxonomy, morphology, biogeography, convergent evolution, character polarity, total-evidence

## Abstract

Within the Formicidae, the higher classification of nearly all subfamilies has been recently revised due to the findings of molecular phylogenetics. Here, we integrate morphology and molecular data to holistically address the evolution and classification of the ant genus *Lasius*, its tribe Lasiini, and their subfamily Formicinae. We accomplish this through a critical re-examination of morphology of extant and fossil taxa, molecular phylogenetic analyses, total-evidence dating under fossilized birth-death process, phylogeography, and ancestral state estimation. We use these results to provide revised taxonomic definitions for the Lasiini and select genera, and we provide a key to the genera of the Lasiini with emphasis on the *Lasius* genus group. We find that the crown Lasiini originated around the end of the Cretaceous on the Eurasian continent and is divisible into four morphologically distinct clades: *Cladomyrma*, the *Lasius* genus group, the *Prenolepis* genus group, and a previously undetected lineage we name *XXX* **gen. n.** The crown of the *Lasius* genus group is considerably younger than that of the *Prenolepis* genus group, indicating that extinction has played a major role in the evolution of the former clade. *Lasius* itself is divided into two well-supported monophyletic groups which are approximately equally speciose. We present evidence that temporary social parasitism and fungiculture arose in *Lasius* two times independently. Additionally, we recover the paraphyly of three *Lasius* subgenera and propose replacing all subgenera with an informal species group classification: *Lasius* = *Acanthomyops* **syn. rev.**, = *Austrolasius* **syn. n.**, = *Cautolasius* **syn. n.**, = *Chthonolasius* **syn. n.**, = *Dendrolasius* **syn. n.** Total-evidence analysis reveals that the Baltic-region amber fossil species †*Lasius pumilus* and †*Pseudolasius boreus* are misplaced to genus; we therefore designate †*XXX* **gen. n.** for the former and †*XXX* **gen. n.** for the latter. Further, we transfer †*XXX* and †*Glaphyromyrmex* out of the tribe, considering the former to be *incertae sedis* in the subfamily, and the latter a member of the Formicini (**tribal transfer**). Two final taxonomic actions are deemed necessary: synonymy of *Lasius escamole* Reza, 1925 with *Liometopum apiculatum* Mayr, 1870 **syn. n.** (**subfamilial transfer**), and transfer of *Paratrechina kohli* to *Anoplolepis* (**tribal transfer**, forming *A. kohli* (Forel, 1916) **n. comb.**).

**Summary of taxonomic actions:**

1. Subgenera of *Lasius* synonymized: *Lasius* = Acanthomyops **syn. rev**. = Austrolasius **syn. n**. = Cautolasius **syn. n.** = Chthonolasius **syn. n.** = Dendrolasius **syn. n.**
2. *Lasius myrmidon* transferred to *XXX* **gen. n.** (Lasiini, *XXX* genus group).
3. †*Lasius pumilus* transferred to †*XXX* **gen. n.** (Lasiini, *XXX* genus group).
4. †*Pseudolasius boreus* transferred to †*XXX* **gen. n.** (*incertae sedis* in Formicinae) (**tribal transfer**).
5. †*Glaphyromyrmex* transferred to the Formicini from the Lasiini (**tribal transfer**).
6. *Lasius escamole* Reza, 1925 synonymized with *Liometopum apiculatum* Mayr, 1870, syn. n. (subfamilial transfer).
7. *Paratrechina kohli* (Forel, 1916) transferred to *Anoplolepis* (Plagiolepidini) (**genus and tribal transfer**).

## Introduction

Ant taxonomy is a rapidly transforming field, incorporating studies across data types and systematic methodologies. Built from the synthesis of all species-level and higher taxonomic works on the Formicidae, the morphology-based systematic hypotheses of Brown (*e.g.*, 1954; see also Keller 2011) and Bolton (*e.g.*, 1990a,b,c, 1994, 1995, 2003) were first tested using parsimony methods (*e.g.*, Ward 1990, 1994; Agosti 1991; Baroni Urbani *et al*. 1992; Shattuck 1992, 1995; Lattke 1994; Grimaldi *et al*. 1997; Agosti *et al*. 1999, Keller 2000) followed by likelihood-based and Bayesian phylogenetics (*e.g*., Felsenstein 1983, 1988, 2001, 2003). Whereas a minority of ant phylogenetic studies combined morphological and multi-locus data (Ward & Brady 2003; Astruc *et al*. 2004; Janda *et al*. 2004; Ward & Downie 2005; Maruyama *et al*. 2008) and/or included fossils as terminals (Barden *et al*. 2017a, Prebus 2017), most phylogenetic studies have only utilized data from Sanger-sequenced loci (*e.g.*, Moreau *et al*. 2006; Brady *et al*. 2006; Ward *et al*. 2010, 2015; Ward & Fisher 2016; Borowiec *et al*. 2019). More recently, high-throughput sequencing platforms enabled genomic approaches to ant systematics, such as targeted enrichment of ultraconserved elements (UCEs) (*e.g.*, Faircloth *et al*. 2015; Blaimer *et al*. 2015, 2016, 2018; Jesovnik *et al*. 2017; Branstetter *et al*. 2017a,b,c; Prebus 2017, 2021; Sosa-Calvo *et al*. 2018; Borowiec 2019; Branstetter & Longino 2019), RADseq (Fischer *et al*. 2015, Darwell *et al*. 2019, Liu *et al*. 2020), and transcriptomics (Jesovnik *et al*. 2016, Romiguier *et al*. 2018). Here, we employ an array of phylogenetic methods, including traditional comparative morphology, total-evidence divergence dating (Ronquist *et al*. 2012a, 2016), phylogeographic analysis (Bouckaert *et al*. 2014; Matzke 2014), ancestral state estimation (Paradis *et al*. 2004, Paradis & Schliep 2018), and species tree estimation (Mirarab & Warnow 2015) to holistically address the phylogeny, evolution, and classification of the genus *Lasius* F., 1904, its tribe Lasiini Ashmead, 1905, and the subfamily Formicinae Lepeletier de Saint-Fargeau, 1934 (see Table 1 for a summary of our questions and methods).

**Table 1.** Overview of study design, explaining which datasets and analyses were used to answer our primary questions.

| Question | Data | Program | Analysis |
| --- | --- | --- | --- |
| **Phylogenetic Analyses** |  |  |  |
| **1. Is *Lasius* monophyletic?** | 89 taxa, 959 UCE locus alignments<br>135 taxa, 11 Sanger loci<br>55 taxa, 11 Sanger loci | UCEs: ASTRAL-II<br>Sanger: MrBayes 3.2.6, IQ-TREE | UCE: summary coalescence<br>135 taxa, 11 Sanger loci: Bayesian tree inference, & maximum likelihood bootstrap<br>55 taxa, 11 Sanger loci: stepping stone, concordance factor |
| **2. Are the subgenera of *Lasius* monophyletic?** | 55 taxa, 11 Sanger loci | MrBayes 3.2.6, IQ-TREE | Bayesian: Bayesian tree inference, stepping stone<br>ML: maximum likelihood bootstrap, concordance factor |
| **3. What are the relationships of fossil Formicinae?** | 154 taxa: 11 Sanger loci for 135 extant taxa, and 110 binary morphological characters for 136 extant and 18 extinct taxa | MrBayes 3.2.6 | Tree search: (clock vs. non-clock; morphology-only vs. combined data) |
| **Evolutionary History** |  |  |  |
| **4. Under what paleoclimatological conditions and where biogeographically did the Lasiini and its internal clades originate?** | 135 extant taxon, 11 Sanger loci chronogram trimmed from 154 taxon tree generated in Question 4 | BioGeoBEARS | BioGeoBEARS: several models (with and without +J) |
| **5. What was the evolutionary sequence of transformation for taxonomically important characters and life history traits?** | 135 taxon chronogram from Question 5, 5 binary morphological characters and 1 ternary morphological character; 21 taxon chronogram trimmed from Question 5, 2 binary life history traits | R (*ape*) | *ace* with ER (equal rates) vs. SYM (symmetrical rates) vs. ARD (all rates different) models |
| **Taxonomy** |  |  |  |
| **6. What are the morphological states that define the Lasiini and focal clades therein?** | Qualitative comparisons (various)<br>Morphometrics (104 specimens, 56 spp.) | Qualitative: N/a<br>Quantitative: R (ggplot) | Qualitative: Key + Tabulation<br>Quantitative: Plotting |

*Lasius* is a northern temperate ant genus that is important in terms of its relative ecological impact and as a model for studying social insect biology. For example, *Lasius* was a frequent choice in both early and recent work on pheromone communication (Bergström & Löfqvist 1970; Holman et al. 2013), dominance hierarchy, competition, and succession in ant communities (Pontin 1961; Traniello & Levings 1986; Markó & Czechowski 2004; Parr & Gibb 2010), and the evolution of social parasitism (Hasegawa 1998; Janda et al. 2004), among other topics (Hölldobler & Wilson 1990; Quque & Bles 2020). *Lasius* ants are particularly known for well-documented symbioses with honeydew producing insects and for temporary social parasitism. The latter phenomenon is a mode of colony foundation for many species of *Lasius*, and involves parasite queens invading nests of other *Lasius*, killing the host queen, and using subordinated workers to raise the first batch of their own workers. This is followed by attrition of the host and eventually a colony consisting solely of parasite conspecifics. Social parasitism is known in species currently classified in subgenera *Austrolasius, Acanthomyops, Chthonolasius,* and *Dendrolasius* (Hölldobler & Wilson 1990; Buschinger 2009; Raczkowski & Luque 2011). Certain species of the subgenus *Chthonolasius* are known to use ascomycete fungi to bind masticated wood and soil to reinforce nest walls. This association is considered an example of fungiculture because the ants provide the fungi with honeydew and prevent overgrowing by competing fungal species (Schlick-Steiner et al. 2008).

Historically, *Lasius* was split by Fabricius (1804) from Linnaeus’s (1758) all-encompassing genus *Formica* based on features of the mouthparts; Latreille (1809) soon after erected the family Formicidae for the ant genera known at the time. The first tribal classification of the family is attributed to the hymenopterist Amédée Michel Louis Lepeletier de Saint-Fargeau (1835), who included *les Formicites*, a precursor to the modern Formicinae. Lepeletier de Saint-Fargeau’s classification of ants incorporated information on wing venation (Lepeletier de Saint-Fargeau 1836). Later investigators relied on natural history, external, as well as internal morphology, especially of the proventriculus, which is a valve located between the crop and midgut of ants and other insects (Eisner 1957). A relatively stable supra-generic classification of the subfamily emerged around the turn of the prior century (Forel 1878, 1886, 1912; Emery 1895, 1925), which was used into the beginning of the 1990s. Agosti (1991) provided a critical reassessment of this system and used new exoskeletal characters to diagnose four informal genus groups in the Formicinae. Expanding on Agosti’s skepticism and system, Bolton (2003) divided the Formicinae into a total of 11 morphologically defined tribes, which were placed in the “lasiine” and “formicine” tribe groups, with three and six tribes, respectively, plus two unplaced tribes.

Ant classification has increasingly become more reliant on phylogenetic trees inferred from molecular data, with subsequent morphological revisionary work. The Formicinae is no exception: Blaimer et al. (2015) provided a comprehensive molecular phylogenetic study of the subfamily which was used for a morphological reassessment of its tribal classification, emphasizing the hyperdiverse Camponotini (Ward et al. 2016; Ward & Boudinot 2021). Like Bolton (2003), Ward et al. (2016) also recognized 11 tribes, for which the generic composition of most remained largely unchanged. The notable exceptions included tribes Lasiini, Melophorini, and Plagiolepidini, whose scope was substantially revised but for which no updated morphological circumscription was provided. Within the Lasiini, the modern taxonomy of *Lasius* was initiated by Wilson’s revision of the genus (Wilson 1955), for which six subgenera have been recognized (Ward 2005). Monophyly of the subgenera has not been questioned except for the nominotypical subgenus (Janda et al. 2004; Maruyama et al. 2008; see Fig. 1A, B, C), but the monophyly of *Lasius* was cast into doubt by Blaimer et al. (2015), whose analysis of Sanger data suggested paraphyly of the genus with respect to *Myrmecocystus*. A more recent study using UCEs and a limited taxonomic sampling of the genus recovered *Lasius* as monophyletic (van Elst et al. 2021).

**Fig. 1.**
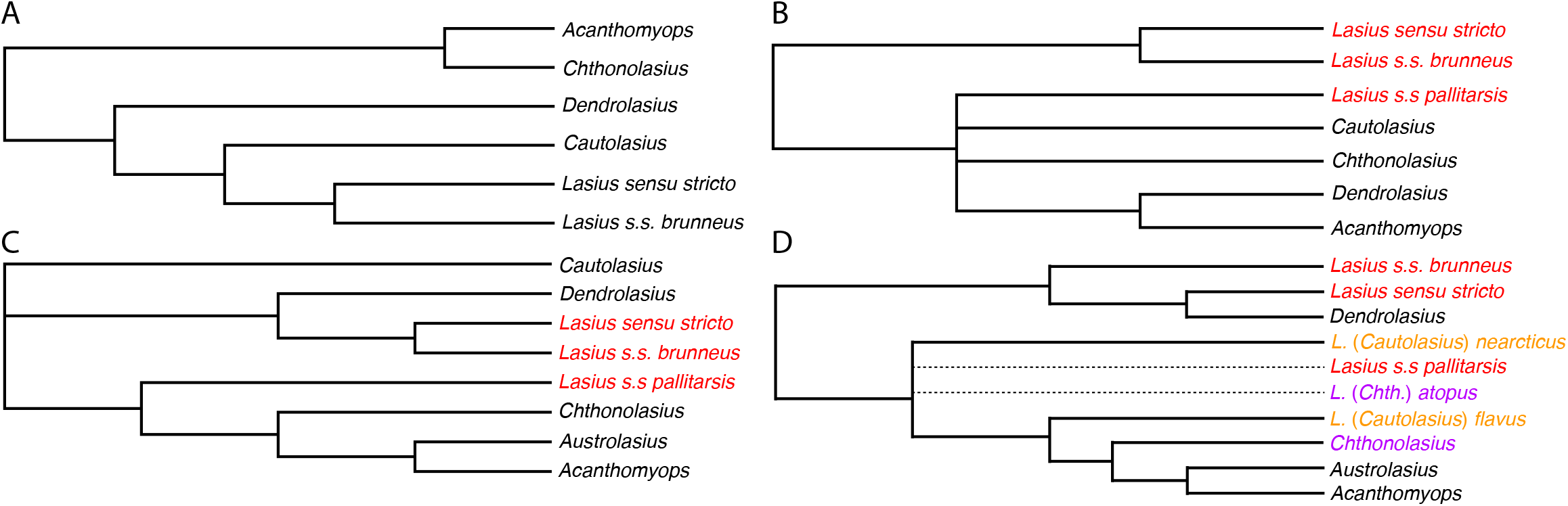
Comparison of phylogenetic hypotheses for *Lasius*; red indicates paraphyly of *Lasius sensu stricto*, orange of *Lasius* (*Cautolasius*), and violet of *Lasius* (*Chthonolasius*). (A) Intuition-based topology from Wilson (1955). (B) and (C) combined morphological and mitochondrial topologies of Janda *et al*. (2004) and Maruyama *et al*. (2008). (D) Consensus topology from the current study; see Table 4 for Bayes Factor tests of our constraint analyses.

In this contribution we focus on the genus *Lasius* and the tribe Lasiini, integrating molecular and morphological methods to address several questions about their classification and evolution (Table 1). In addition to the monophyly of *Lasius* and its subgenera (Table 1, *Questions 1 & 2*), we are also interested in the fossil history of the genus and the Lasiini (e.g., Mayr 1868; Wheeler 1915; Wilson 1955; LaPolla et al. 2013; Barden 2017a), from both taxonomic (Table 1, *Question 3*) and evolutionary perspectives (Table 1, *Questions 4 & 5*). We propose a revised morphological definition of *Lasius*, the Lasiini, and a number of newly recognized nodes on the formicine phylogeny, resulting from explicit integration of morphology of both extant and fossil lineages into our phylogenetic analyses. We also provide a systematic, illustrated key to the extant genera of the tribe (Table 1, *Question 6*). Our overarching hope is to encourage the reconciliation of morphological and molecular systematics.

## Materials and methods

### Study design

Our study addresses six questions outlined in Table 1. A summary of methods is provided below with additional details available as supplementary text. In brief, our objective is to holistically evaluate the phylogeny and evolution of extant and extinct *Lasius* and Lasiini in the context of the Formicinae. We employed an array of tools to three datasets comprising a broad sample of the Formicinae and selected non-formicine outgroups: (1) a set of previously published 959 individual UCE locus alignments for 89 extant terminals (Blaimer *et al*. 2015, *Question 1*); (2) a set of 11 Sanger loci for 135 extant terminals (*Questions 1, 3–5*); (3) a set of 11 Sanger loci for 55 extant terminals in the Lasiini (*Question 2*); and (4) a set of 110 binary morphological characters for 154 terminals (136 extant and 18 extinct) (*Question 3*). Additionally, we tested the polarity of six morphological characters traditionally used in the classification of the Formicinae (*Question 5*). In order to revise the generic classification of the Lasiini and to provide a new key, we comparatively evaluated several qualitative traits, and took six linear measurements for 109 specimens of 56 species (*Question 6*). The three new genera we recognize are registered with ZooBank (LSIDs in respective taxonomic accounts below). All Bayesian inference (BI) and maximum likelihood (ML) analyses were conducted on the CIPRES Science Gateway V. 3.3 (Miller *et al*. 2010, accessed at http://www.phylo.org/portal2).

### Sanger sequence data

New sequence data were generated from a selection of 20 *Lasius* and two *Myrmecocystus* species. Our sampling of *Lasius* was designed to encompass the morphological, geographical, and life history variation within the genus (Table 2), spanning all currently recognized subgenera (Wilson 1955; Bolton 2003; Ward 2005), and was partially guided by previous molecular phylogenies (Janda *et al*. 2004, Maruyama *et al*. 2008). Notably, we included a species closely related to or conspecific with the aberrant and previously unsampled species, *Lasius atopus*, from California, U.S.A. as well as another unsampled species, *Lasius myrmidon*, described from Greece (Mei 1998). Although we included only three species of *Myrmecocystus*, they were chosen to span the root node of the genus (O’Meara 2008, van Elst et al. 2021). We also included data from all genera recognized in the *Prenolepis* genus group (LaPolla *et al*. 2010, LaPolla *et al*. 2012), the majority of formicine genera, and a sample of formicoid outgroups. Voucher specimens from DNA extractions made for this study are deposited in the University of California Bohart Museum of Entomology collection (UCDC) and collection data associated with these specimens can be found in Table 2 and on AntWeb at http://antweb.org/.

**Table 2.** Voucher specimen collection data for sequences generated *de novo*. Specimen codes link to more detailed collection data on AntWeb (http://www.antweb.org); all specimens were deposited in the Bohart Museum of Entomology (UCDC) except for *L. sonobei*, which was a destructive extraction (collection code: PSW14663, duplicates available).

| Species | Specimen code | Locality (year) | Collector |
| --- | --- | --- | --- |
| *Lasius americanus* nr | CASENT0732004 | CA, U.S.A. (2014) | Borowiec, M.L. |
| *Lasius atopus* nr | CASENT0234858 | CA, U.S.A. (2014) | Boudinot, B.E. *et al.* |
| *Lasius brevicornis* | CASENT0234864 | CA, U.S.A. (2012) | Borowiec, M.L. |
| *Lasius brunneus* | CASENT0234862 | Dolny Slask, Poland (2010) | Borowiec, M.L. |
| *Lasius carniolicus* | CASENT0731085 | Dolny Slask, Poland (2012) | Salata, S. |
| *Lasius emarginatus* | CASENT0234865 | Dolny Slask, Poland (2010) | Borowiec, M.L. |
| *Lasius flavus* | CASENT0234860 | Dolny Slask, Poland (2010) | Borowiec, M.L. |
| *Lasius fuliginosus* | CASENT0234861 | Dolny Slask, Poland (2010) | Borowiec, M.L. |
| *Lasius latipes* | CASENT0234866 | CA, U.S.A. (2007) | Ward, P.S. |
| *Lasius myops* | CASENT0732040 | N Calabria, Italy (2014) | Borowiec, L. |
| *Lasius nearcticus* | CASENT0732003 | MA, U.S.A. (2008) | Alpert, G.D. & Borowiec, M.L. |
| *Lasius pallitarsis* | CASENT0234859 | CA, U.S.A. (2014) | Borowiec, M.L. |
| *Lasius platythorax* | CASENT0732002 | N Dalmatia, Croatia (2000) | Borowiec, M.L. & Poprawska, M. |
| *Lasius psammophilus* | CASENT0732000 | Dolny Slask, Poland (2009) | Borowiec, M.L. |
| *Lasius sabularum* | CASENT0234863 | Dolny Slask, Poland (2010) | Borowiec, M.L. |
| *Lasius sonobei* | CASENT0730575 | Hokkaido, Japan (2002) | Ward, P.S. |
| *Lasius spathepus* | CASENT0732006 | Hokkaido, Japan (2002) | Ward, P.S. |
| *Lasius subumbratus* nr | CASENT0731084 | CA, U.S.A. (2014) | Ward, P.S. |
| *Lasius turcicus* | CASENT0732001 | Rhodes, Greece (2008) | Borowiec, M.L. |
| *XXXmyrmidon* | CASENT0811544 | Korinthia, Greece (2013) | Borowiec, L. |
| *Myrmecocystus mexicanus* | CASENT0234867 | CA, U.S.A. (2014) | Borowiec, M.L. |
| *Myrmecocystus tenuinodis* | CASENT0234868 | CA, U.S.A. (2004) | Ward, P.S. |

We used the DNeasy Blood and Tissue Kit (Qiagen Inc., Valencia, CA, USA) to conduct non-destructive DNA extractions by piercing the cuticle of each specimen, then soaking the specimens overnight in a solution of proteinase K and the lysis buffer provided with the kit. For the remainder of the extraction, we followed the manufacturer’s protocol except for eluting the extract with sterilized water rather than the supplied buffer. With these 22 extractions, we amplified and sequenced fragments of eight nuclear protein coding genes: abdominal-A (*abdA*), arginine kinase (*ArgK*), rudimentary (*CAD*), elongation factor 1 α copy F2 (*EF1aF2*), long-wave rhodopsin (*LW Rh*), DNA topoisomerase 1 (*Top1*), ultrabithorax (*Ubx*), and wingless (*wg*). Amplifications of desired gene fragments were performed with primers listed in Table S1 using standard PCR methods described in Ward & Downie (2005). Sequencing reactions were carried out on an ABI 3730 Capillary Electrophoresis Genetic Analyzer with ABI Big-Dye Terminator v3.1 Cycle Sequencing chemistry (Applied Biosystems Inc., Foster City, CA).

Sequence base calling was performed in Sequencher v5.0 (Gene Codes Corporation, Ann Arbor, MI). Because it has been suggested that arbitrary choice of alleles can bias phylogenetic results in some data sets (Weisrock *et al*. 2012), we retained ambiguous base calls (R or Y) for any potentially heterozygous sites. To expand our taxon sampling, we downloaded sequences from GenBank. For 18 *Prenolepis* genus-group taxa from LaPolla *et al*. (2010, 2012) we obtained five nuclear loci (*ArgK*, *CAD*, *EF1aF1, EF1aF2*, and *wg*); we obtained nine nuclear loci (*abdA*, *ArgK*, *CAD*, *EF1aF1*, *EF1aF2*, *LW Rh*, *Top1*, *Ubx*, and *wg*) and two ribosomal loci (*18S* and *28S*) for an additional 116 formicine and 19 non-formicine taxa from datasets published by Brady *et al*. (2006), Branstetter (2012), Brady *et al*. (2014), Blaimer *et al*. (2015), Chomicki *et al*. (2015), and Ward *et al*. (2015). The final data set comprised a total of eleven nuclear loci for 135 taxa spanning the Formicinae and including non-formicine outgroups, with 21 *Lasius* species, 3 *Myrmecocystus* species, and 30 *Prenolepis* genus group species. GenBank identifiers for all sequences used in this study are listed in Table S2.

### Morphological data

Worker specimens of extant and extinct species were examined from the American Museum of Natural History (AMMH) and collections housed at the University of California, Davis: the Bohart Museum of Entomology (UCDC), and the personal collections of Philip S. Ward (PSWC) and the authors (BEBC, MLBC, MMPC). Additionally, morphological characters were evaluated from type specimens and other material imaged on AntWeb (http://antweb.org). We constructed two separate datasets for our study: (1) continuous morphometric data to differentiate “*L. myrmidon*” and “†*L. pumilus*” relative to other *Lasius* (measurements are defined in Appendix S1, and data is reported in Lasiini_eye_metrics_XXX_vs_XXX.csv, available on Dryad, https://doi.org/10.25338/B8B645), and (2) discretized binary morphological data to test our assumptions of the taxonomic placement of fossils informative for the Lasiini (characters are defined in Appendix S2, and states scored for each taxon are in Lasiini_w_outgroups_w_fossils_154t_morphology.fasta, available on Dryad, https://doi.org/10.25338/B8B645). For the discretized dataset, worker-based observations were made of 110 morphological characters coded in binary fashion for phylogenetic analysis (character list reported in Appendix S2). These data were scored for 150 terminals in total, of which 18 were fossils and 132 were extant Formicinae. Some of the characters were derived from prior studies, while others were interpreted *de novo* from comparative morphological study across the Formicidae. Each character is treated as a state which is either observed to be *true* (1) or *false* (0) for a given specimen. Therefore, some complex (multistate) characters were separated into multiple characters, with observed absences and inapplicable states scored as *false*, similar to the “Composite Coding” method described by Strong & Lipscomb (1999). An effort was made to score characters in a homology-neutral way, *i.e.*, that each *true* statement is based on the morphological definition, without assuming *a priori* that the state must be homologous when observed across the phylogeny. See the “Species group classification of *Lasius*” section below for a record of examined *Lasius* species.

### Phylogenetic analyses

#### Alignment and partitioning of Sanger sequence data

We generated two molecular datasets from our initial set of sequences: ‘Lasiini_w_outgroups_135t’ contains the full taxon set, with 135 taxa spanning the Formicinae, including formicoid clade outgroups (Brady *et al*. 2006); ‘Lasiini_55t’ is a subset of ‘Lasius_w_outgroups_135t’, containing only species from the genera of the Lasiini. For both datasets, individual loci were aligned with the L-INS-I algorithm in MAFFT v7.419 (Katoh *et al*. 2002; Katoh & Standley 2013), and exonic regions were inspected in Aliview (Larsson 2014) to ensure that the alignments were in frame. The individual alignments were trimmed with GBLOCKS v0.91b (Castresana 2000), adjusting the default settings to trim less stringently: b1 and b2 were set to half of the total taxon set in each locus (plus one), b3 was set to 12, b4 to 7, and b5 was set to half. After trimming, the ends of each sequence alignment were further trimmed such that the alignment began with the first codon position and ended with the third codon position. If the locus contained introns, the alignment was then split into exonic and intronic regions using AMAS (Borowiec 2016b). We concatenated the resulting data into the ‘Lasius_w_outgroups_135t’ and ‘Lasiini_55t’ matrices, as well as individual gene alignments for each matrix.

As the assumption that evolution of all sequence data is homogeneous is often violated in empirical data (Buckley *et al*. 2001), we partitioned our matrices into subsets of similarly evolving sites for our concatenated and single gene analyses. To find the optimal partitioning scheme and best substitution models we used PartitionFinder v2 (Lanfear *et al*. 2016) on our concatenated molecular datasets as well as the individual gene alignments. As input, we used blocks of data which represent codon positions within exonic regions of protein coding loci; introns and ribosomal loci were treated as individual data blocks. We included JC, K80, HKY, SYM, F81, and GTR, all with or without +Γ or +I, but we excluded models invoking both gamma and proportion of invariable sites (*i.e.*, +Γ+I) as it has been shown that interaction of these parameters may cause anomalies in Bayesian inference (Sullivan & Swofford 2001, Yang 2006). To find the optimal scheme, we chose the Bayesian information criterion (BIC) and the ‘greedy’ algorithm.

#### Phylogenetic analysis of molecular-only data

We used MrBayes 3.2.6 (Ronquist *et al*. 2012b) for BI on concatenated and single gene alignments (Table 1, *Questions 1–3*) using the model and partition schemes estimated from the PartitionFinder v2 analyses. State frequencies, substitution rates, shape of gamma distribution of substitution rates at sites, and proportion of invariable sites were unlinked and allowed to vary among partitions, while branch lengths remained linked. We ran each analysis for 25 million generations with two runs, four chains each, and sampled parameters every 2500 generations, leaving burnin at the default 25%. We used the following statistics to diagnose MCMC: PSRF values approaching 1.0, proposal acceptance rates between 20–70%, and standard deviation of split frequencies with values of approximately 0.01 or less. Additionally, we used the program Tracer v1.6.0 (Rambaut *et al*. 2014) to evaluate trace plots and minimum effective sample size values (should be > 200) for each parameter in the different runs.

For phylogenetic inference under ML (Table 1, *Question 2*), we used IQTREE v2 (Minh et al. 2020). We used the model and partition schemes estimated from PartitionFinder v2, allowing each partition to have its own rate of evolution (-p), and 1000 ultrafast bootstrap replicates. Trees for both Bayesian and ML analyses were manipulated with FigTree v1.4.2 (Rambaut & Drummond 2014). Additionally, we performed a gene and site concordance factor analyses for each concatenated dataset using the consensus tree and single gene trees from the MrBayes analyses above as input.

Based on the results from the above ML and BI analyses, we performed stepping-stone analyses of several contrasting topological hypotheses (Table 1, *Questions 1 & 2*). These follow Bergsten *et al*. (2013), who recommended comparisons of analyses under equally constrained topologies (Xie *et al*. 2011) to estimate marginal log likelihoods in MrBayes. The hypotheses we tested include: **(1)** the placement of *Lasius myrmidon* as either with *Lasius* and *Myrmecocystus* or with the *Prenolepis* genus group; **(2)** monophyly of *Lasius* with respect to *Myrmecocystus*; **(3)** monophyly of *Lasius s. str.*; **(4)** monophyly of *Cautolasius*; and **(5)** monophyly of *Chthonolasius* (Fig. S1). For these analyses, we ran MrBayes for 25.5 million generations with two runs, four chains, fifty steps, sampling every 100 generations, and with the alpha shape parameter set to 0.4. We took the mean marginal log likelihood of the two stepping-stone runs for each analysis, subtracted the likelihood of the alternative hypothesis from the null, then doubled this number. Results of these Bayes factor tests are interpreted after Kass & Raftery (1995), with these specific 2ln_e_(BF_01_) ranges: 0–2, “not worth explaining”; 2–6, “positive support”; 6–10, “strong support”; and > 10, “very strong support”.

Additionally, because *Lasius* was recovered as paraphyletic with respect to *Myrmecocystus* in the Sanger data analyses of Blaimer *et al*. (2015), we employed summary coalescent species tree analysis in ASTRAL-II (Mirarab & Warnow 2015) with individual UCE locus trees from Blaimer *et al*. (2015) supplied by the author as input. We did not use the statistical binning pipeline (Mirarab et al. 2014; Bayzid et al. 2015) due to recent criticism (Adams & Castoe 2019), but instead used individual gene trees as input to ASTRAL-II version 4.10.11, which we ran under default settings.

#### Phylogenetic analysis of combined data and divergence dating

We analyzed a morphology-only data matrix and two combined-data matrices with MrBayes to evaluate placement of fossil taxa and to estimate informative priors on relaxed clock models (Table 1*, Question 3*). For morphology-only analysis, we built the ‘Lasiini_w_outgroups_w_fossils_morphology_154t’ dataset, with 110 morphological characters for 154 taxa. For our first combined-data matrix, we created a 136-taxon dataset containing our partitioned 135-taxon 11-gene matrix and morphological data from extant taxa, adding *Paratrechina kohli,* which was represented only by morphological data (‘Lasiini_w_outgroups_136t’). For our second combined-data matrix, we added morphological data from 18 fossil taxa to the previous matrix (‘Lasiini_w_outgroups_w_fossils_154t’). All morphological data were included in a single partition to which we applied the Mk model (Lewis 2001), accounting for among character rate variation using +Γ. As rate asymmetry has been shown to have important consequences for morphological analysis (Wright *et al*. 2016; Klopfstein & Spasojevic 2019), particularly in the otherwise robust total-evidence dating framework (Klopfstein *et al*. 2019), the assumption of equal forward and reverse transformations was relaxed by setting the ‘symdirihyperpr’ to a uniform distribution (‘fixed(1.0)’) in our input files, thus allowing MrBayes to estimate asymmetric stationary frequencies (*π*) across states. To account for ascertainment bias (Lewis 2001), we set coding for the morphological partition to “variable”.

With the three datasets above, we performed alternate analyses combining different sets of parameters (Table 3), examining the effect on the placement of fossil taxa when analyzing morphological data in isolation in comparison to the results of combined-data analyses in the absence of a molecular clock. To estimate informative priors for the two commonly used relaxed clock models in MrBayes (IGR and TK02), we followed the methods used in Ronquist et al (2012a) with our Lasiini_w_outgroups_136t dataset. The two clock models with informative priors were then compared against each other using stepping-stone analysis with the Lasiini_w_outgroups_w_fossils_154t dataset. Because the results of the stepping-stone analyses were ambiguous, we analyzed the Lasiini_w_outgroups_w_fossils_154t dataset with both clock models for comparison. For the analyses of the Lasiini_w_outgroups_w_fossils_154t dataset, we used the fossilized birth-death branch length (FBD) model (Heath *et al*. 2014) and a uniform clock for comparison. All analyses included a root node topology constraint (*i.e.*, monophyly of core formicoids enforced), but were otherwise unconstrained. Because our taxon sample does not meet the expectations of the “diversified sampling” approach (Zhang *et al*. 2016), we implemented a random sample stratification for FBD analyses. For fossil terminals, uniform distributions were used for stratigraphic ages (New Jersey: 94.3–89.3 mya; Canadian: 84.9–70.6 mya; Baltic: 37.2–33.9 mya; Dominican: 20.4–13.7 mya). The root node prior age was set to the estimated age of the oldest known fossil in the Formicidae, an offset exponential distribution with a minimum of 90 and mean of 99 mya. We ran each analysis for 100 million generations with two runs of four chains each, with sampling every 10,000 generations, and the default burnin of 25%. MCMC diagnoses were conducted as for the molecular-only analyses described above.

**Table 3.** Datasets and analyses used for divergence dating.

| dataset name | purpose of analyses | data type | taxa | clock type |
| --- | --- | --- | --- | --- |
| Lasiini w outgroups w fossils morphology 154t | fossil placement comparison | morphology | extant & fossil | no clock |
| Lasiini_w_outgroups_136t | relaxed clock prior estimation | combined | extant taxa | no clock vs. strict clock |
| Lasiini_w_outgroups_w_fossils_154t | fossil placement comparison,<br>divergence dating | combined | extant & fossil | relaxed clock: IGR vs. TK02;<br>uniform clock: IGR vs. TK02 |

#### Historical biogeography

After divergence dating, the post-burnin consensus chronogram resulting from the MrBayes fossilized birth-death analysis was trimmed to 55 extant taxa of the Lasiini with the ‘drop.tip’ function in the R package ‘phytools’ (Revell 2012) and used as input for the likelihood-based biogeographic program BioGeoBEARS (Matzke 2013). We used a coarse biogeographical classification following previous studies (*e.g.*, Ward *et al*. 2015): Neotropical, Nearctic, Palearctic, Afrotropical, Indomalayan, and Australasian regions, with Wallace’s line dividing the last two areas. The maximum ancestral species range size was set to two areas, and the matrices of allowed areas and dispersal constraints were set for three periods: before 60 mya, 60–30 mya, and after 30 mya, reflecting tectonic drift referenced from Scotese (2010). We conducted our phylogeographic analyses using the DEC, DIVALIKE, and BAYAREALIKE models, all with or without +J, but due to criticism of +J (Ree & Sanmartín 2018) we excluded results that included this parameter from consideration. The script and input files used in this analysis can be found on Dryad (https://doi.org/10.25338/B8B645).

#### Ancestral state estimation

We employed the *ace* function of the R package *ape* (Paradis & Schliep 2018) to estimate ancestral states of six traditional morphological characters and two life history traits (Table 1, *Questions 5 & 6*). These six characters, described in the next paragraph, were scored for the MrBayes analysis above after trimming fossils from the 154-taxon tree with the ‘drop.tip’ function in the R package ‘phytools’, resulting in a 135-taxon phylogeny representing the Formicinae and outgroups, and a 21-taxon phylogeny representing *Lasius*. We contrasted two evolutionary models for five characters and two traits with binary states: equal (ER) and unequal rates (ARD) of forward and reverse transition. For the abdominal segment III character, which had ternary states, we tested symmetrical rates (SYM) in addition to the ER and ARD models. To determine if the more-complex SYM and ARD models had significantly better fit, we conducted pairwise ANOVA for each character.

We qualitatively reinterpreted the phenotypic diagnostic traits used by Bolton (2003) for classification at the tribe and ‘tribe group’ level, as well as the form of the proventriculus, a classic character in formicine classification (*e.g.*, Emery 1925). The six morphological characters that we reconstructed the ancestral states for are: **(1)** eye placement, with eyes set at or anterior to head midlength as measured from the anterolateral clypeal corners [state 1] or posterior to midlength [state 0]; **(2)** metacoxal separation, with coxae either close-set (“formicoform”) [state 0] or wideset (“lasiiform”) [state 1]; **(3)** abdominal sternum III with transverse sulcus at base of helcium [state 1] or without [state 0]; **(4)** propodeal spiracle circular to elliptical [state 0] or slit-shaped [state 1]; **(5)** abdominal segment III tergosternal suture extending laterally then broadly curving posteriorly or simply linear (“formicoform”/broadly shouldered) [state 0], suture curving forward then narrowly arching posteriorly (“lasiiform”/narrowly-shouldered, low) [state 1], or suture extended dorsally, with free tergite and sternite commencing well away from helcium (“plagiolepidiform”/narrowly-shouldered, high) [state 2]; and **(6)** proventriculus asepalous [state 0] or sepalous [state 1]. Character 6, proventriculus form, was scored as generalizations based on Forel (1878), Emery (1888, 1895, 1925), Eisner (1957), and Prins (1983), and therefore may be inaccurate; future anatomical studies should evaluate the plausibility of our generalized scoring. Note that metacoxal separation, as scored, is also a proxy for a long petiolar foramen and U-shaped petiolar sternum cross-section used by Bolton (2003) to define his “lasiine tribe group”. The two life history traits for which we reconstructed ancestral states are: **(1)** temporary social parasitism, absence [state 0] or presence [state 1]; **(2)** fungiculture, absence [state 0] or presence [state 1].

## Results and discussion

### Alignment and partitioning of the Sanger sequence data

Overall, the 11-gene 135-taxon molecular matrix is 8.3 kbp long and contains 21% missing data, 2902 variable sites (35%) of which 2449 are parsimony informative (30%). The 11-gene 55-taxon molecular matrix is 8.8 kbp long and contains 39% missing data, 1600 variable sites (18%) of which 1010 are parsimony informative (12%) (statistics calculated with AMAS, Borowiec 2016b). The PartitionFinder analysis resulted in a best-scoring scheme with 17 partitions for our 135-taxon matrix, and a 12-partition scheme for our 55-taxon matrix (Table S3).

### Phylogenetic relationships

Results in this section pertain to *Questions 1–3* of Table 1.

#### Question 1: Is *Lasius* monophyletic?

Answer: The extant fauna of *Lasius* is monophyletic with the exclusion of *XXX myrmidon* gen nov., comb. nov.

Across all analyses there is overwhelming support for the placement of *Lasius myrmidon* as sister to the *Prenolepis* genus-group (Table 4). For this reason, we transfer this species to a new genus, forming *XXX myrmidon* gen. nov., comb. nov., which we diagnose in the *Taxonomy* section below. With the exclusion of *M. myrmidon*, we find somewhat low topological support for *Lasius* monophyly using standard measures (0.96 posterior probability [PP], 87 maximum likelihood bootstrap [BS] (Fig.2); 36.6 gene concordance factor [gCF], 42.3 site concordance factor [sCF] (Fig. S2). However, genome-scale coalescence analysis recovers *Lasius* monophyly with respect to *Myrmecocystus* (Fig. 3), and a Bayes Factor constraint test using the Sanger loci results in “very strong support” for monophyly (Table 4). For these reasons, we accept the hypothesis that *Lasius* and *Myrmecocystus* are reciprocally monophyletic.

**Table 4.** Bayes Factor results, testing alternative placements of *XXX* and the monophyly of the subgenera using the Lasiini_55t matrix. The mean value for the best-supported hypothesis for each comparison is bolded.

| SS Run | Marg. Log Likelihood $H_0$ | Marg. Log Likelihood $H_1$ | $2\ln_e(B_{10})$ |
| --- | --- | --- | --- |
|  | XXX + *Lasius* g. g. | XXX + *Prenolepis* g. g. |  |
| 1 | -29156.21 | -29092.2 |  |
| 2 | -29153.15 | -29094.4 |  |
| mean | -29153.79 | **-29092.79** | -122 |
|  | *Lasius* monophyletic | *Lasius* paraphyletic |  |
| 1 | -29068.73 | -29072.46 |  |
| 2 | -29068.86 | -29072.04 |  |
| mean | **-29068.79** | -29072.23 | 6.88 |
|  | *Lasius* s. str. monophyletic | *Lasius* s. str. paraphyletic |  |
| 1 | -29092.4 | -29057.37 |  |
| 2 | -29094.02 | -29058.11 |  |
| mean | -29092.91 | **-29057.67** | -70.48 |
|  | *Cautolasius* monophyletic | *Cautolasius* paraphyletic |  |
| 1 | -29063.91 | -29054.38 |  |
| 2 | -29064.12 | -29054.21 |  |
| mean | -29064.01 | **-29054.29** | -19.44 |
|  | *Chthonolasius* monophyletic | *Chthonolasius* polyphyletic |  |
| 1 | -29100.32 | -29047.78 |  |
| 2 | -29099.17 | -29046.75 |  |
| mean | -29099.59 | **-29047.13** | -104.92 |

**Fig. 2.**
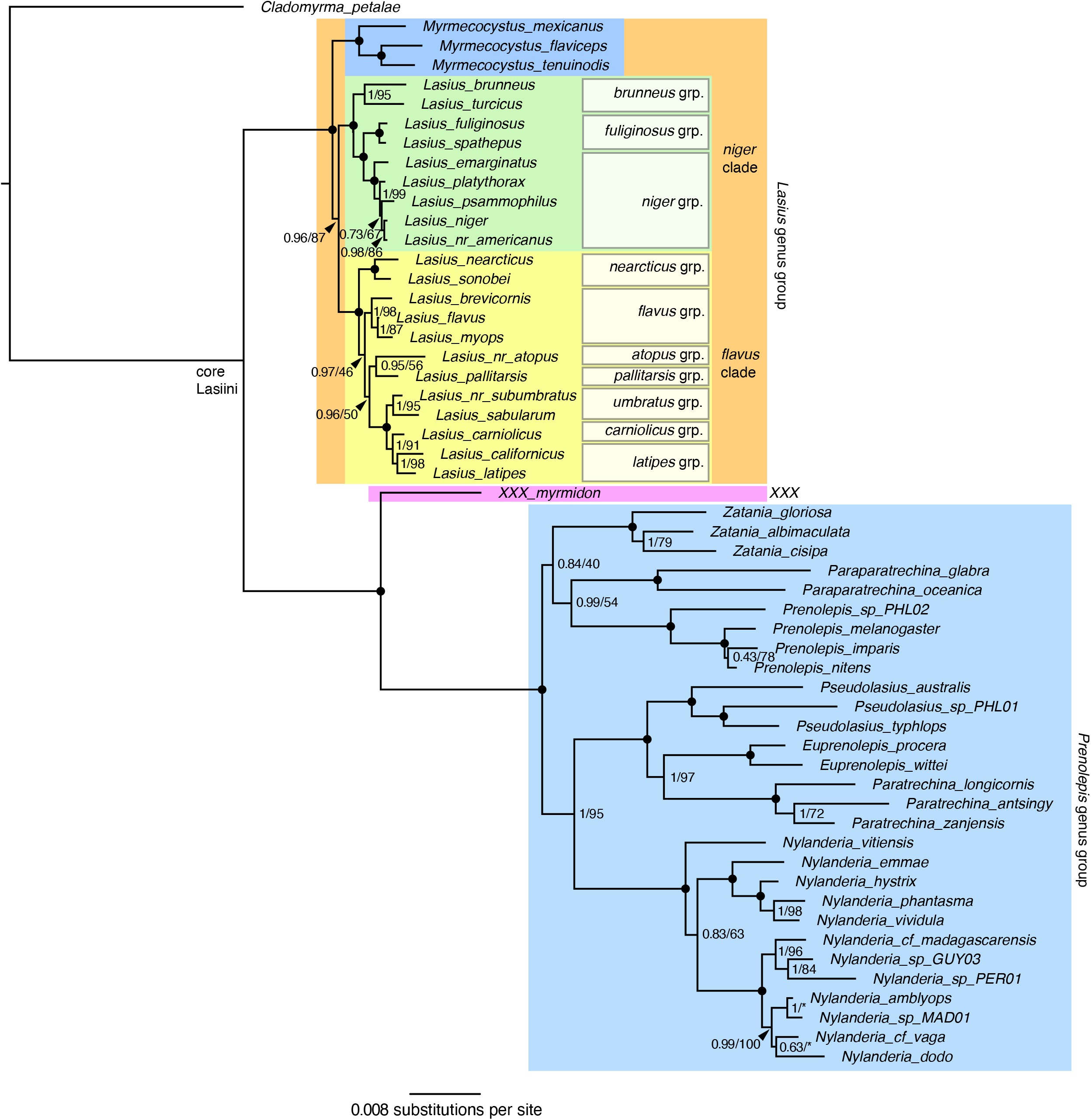
Molecular phylogeny of the Lasiini inferred from Bayesian analysis of the Lasiini_55t data matrix. Node support values are reported as Bayesian posterior probabilities (BI) vs. maximum likelihood bootstrap (ML). Filled circles at nodes indicate full support from both analyses. An asterisk (*) indicates that the topology of the ML analysis diverged from the BI analysis at the given node.

**Fig. 3.**
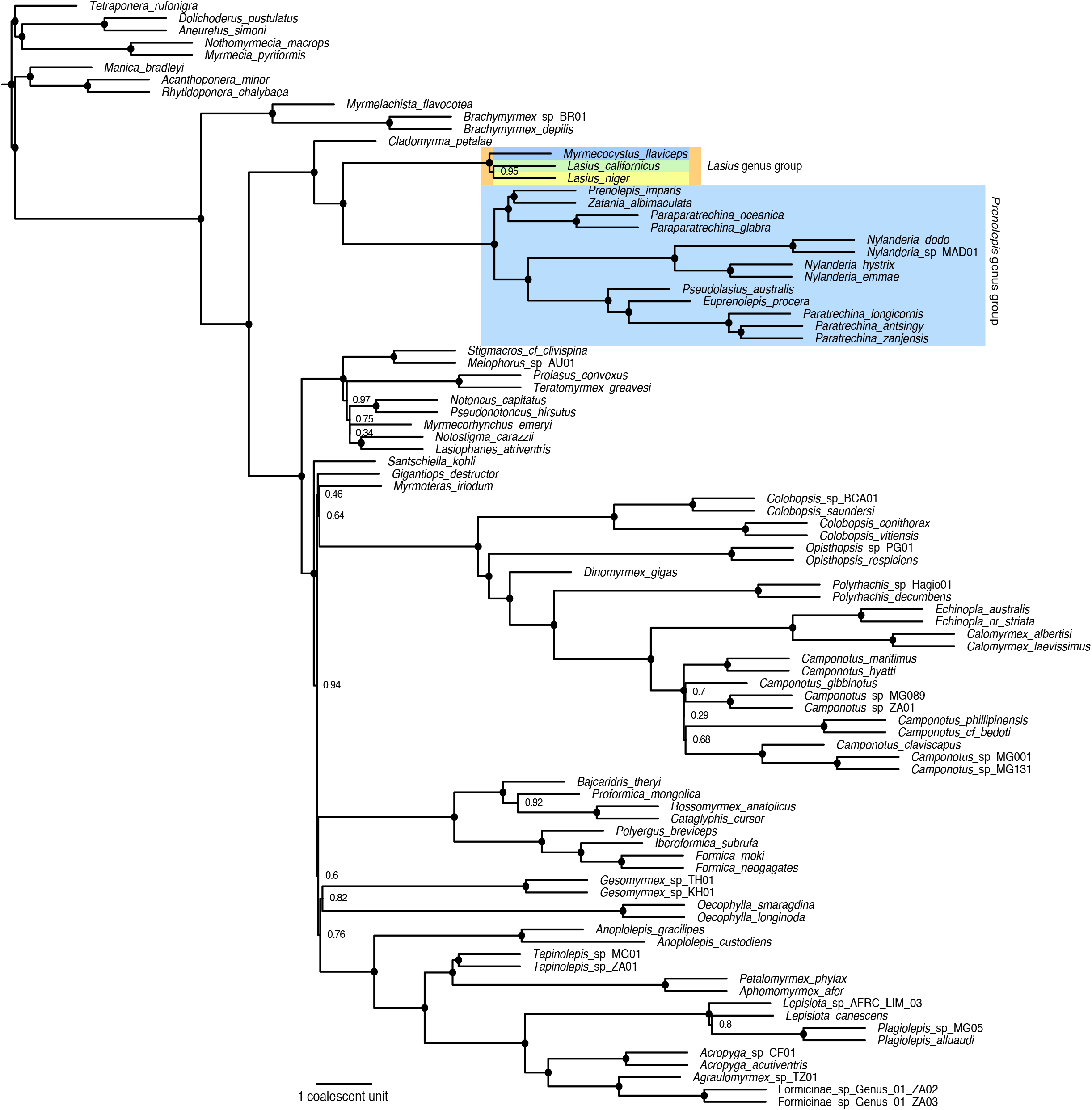
Summary coalescence species-tree phylogeny from ASTRAL-II analysis of 959 UCEs from Blaimer *et al*. (2015). Node support values are given as local posterior probabilities (LPP). Filled circles at nodes indicate full support.

#### Question 2: Are the subgenera of *Lasius* monophyletic?

Answer: Three of the six subgenera of *Lasius*—*Lasius sensu stricto*, *Cautolasius*, and *Chthonolasius*—are not monophyletic (Table 4, Figs. 1D, 2, 4).

**Fig. 4.**
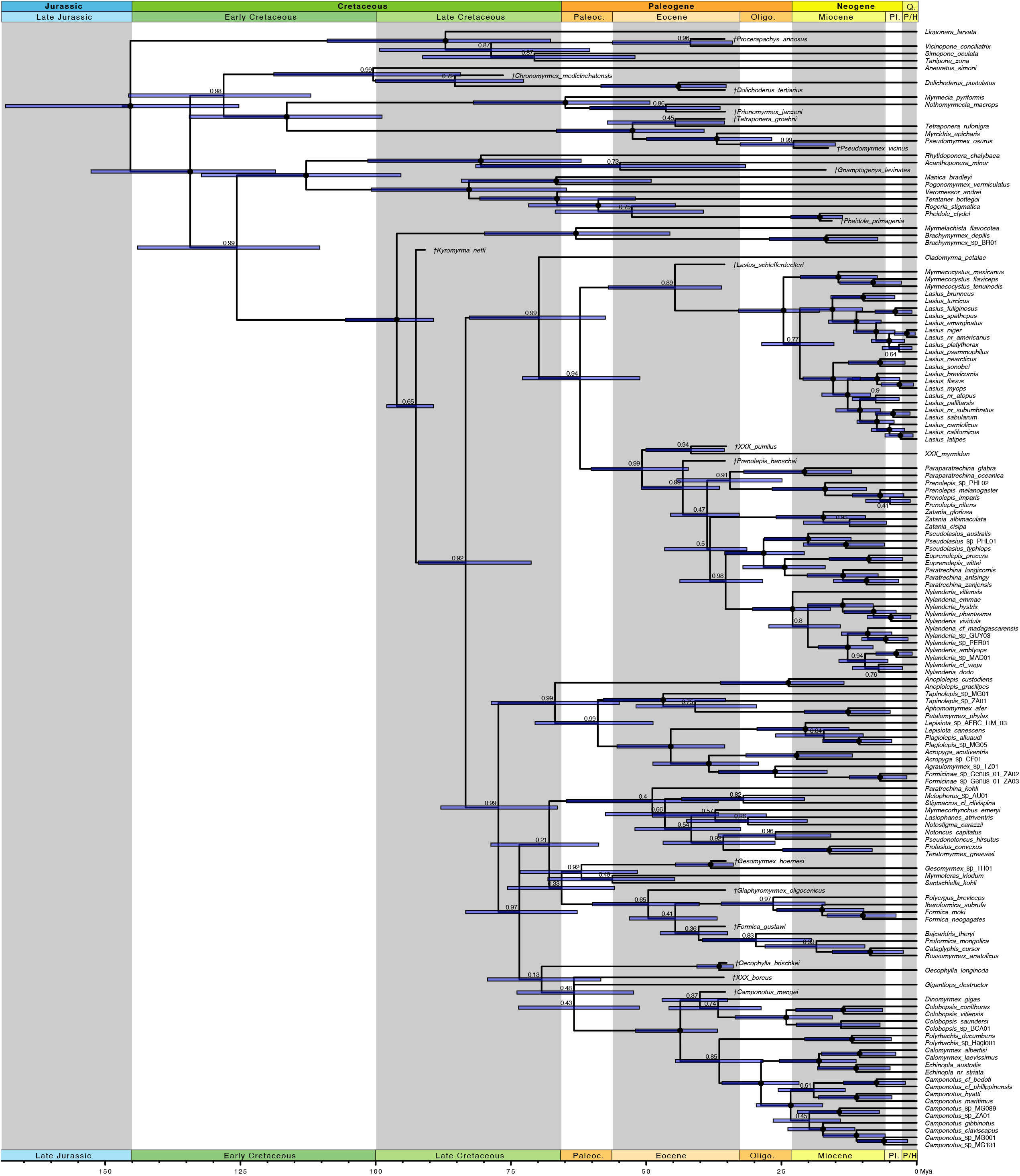
Results from phylogenetic analysis of the Lasiini_w_outgroups_w_fossils_154t combined morphological and molecular dataset of the in MrBayes using the fossilized birth-death branch length prior and the IGR clock variation prior. Fossil taxa are indicated by a dagger preceding the name. Node support values are in Bayesian posterior probability. Filled circles indicate full support. Horizontal blue bars at nodes are 95% highest posterior density (HPD) intervals.

We found that *Lasius*, with the exclusion of *XXX myrmidon*, is divided into two major clades. One clade comprises members of the subgenera *Lasius s. str.* and *Dendrolasius* (here called the “*niger* clade”), while the other includes all other sampled species (here called the “*flavus* clade”). Critically, we found that the *flavus* clade is diagnosable by an enlarged metapleural gland bulla, constituting a new morphological autapomorphy (see *Key to extant genera of Lasiini* below). Within the *niger* clade, the socially parasitic species classified in *Dendrolasius* are nested within taxa currently considered to belong to *Lasius s. str*., while another *Lasius s*. *str*. species, *L*. *pallitarsis*, is a member of the *flavus* clade. The aberrant *Lasius* species near *atopus*, currently classified in *Chthonolasius*, is in the *flavus* clade, sister to *L. pallitarsis* with moderate statistical support. Finally, the four *Cautolasius* sampled here are not monophyletic but instead form two well-supported clades, which we recognize as the *flavus* and *nearcticus* species groups. Because of this phylogenetic arrangement and a continuing trend to abandon subgeneric classification in ants (*e.g.*, Borowiec 2016a), we propose synonymizing *Lasius* subgenera and recognition of informal species groups instead (see *Taxonomy* below).

#### Question 3: What are the relationships of fossil Formicinae?

Answer: Some previously hypothesized relationships are stable, some are sensitive to model choice, and the taxa †*Lasius pumilus*, †*Pseudolasius boreus*, and †*Glaphyromyrmex* require reclassification.

Overall, we find that fossil placement is sensitive the inclusion or exclusion of a clock model. Under “clockless” conditions, fossils were highly nested within the phylogeny (Fig. S3 & S6). However, the topology resulting from clockless morphology-only analysis (Fig. S3) had both very limited support within the Formicinae and was in considerable conflict with previous and present analyses using molecular data. The outgroup fossils were consistent at the subfamily level, while among our formicine fossils, five of the nine Baltic-region fossil taxa were recovered close to their original attributions (Figs. 4, S3, S6–S9): (1) †*Lasius schiefferdeckeri* may be sister to or nested within the *Lasius* genus group; (2) †*Prenolepis henschei* is within the crown of the *Prenolepis* genus group; (3) †*Formica gustawi* is within the crown of the Formicini; (4) †*Oecophylla brischkei* is correctly placed to genus; (5) †*Gesomyrmex hoernesi* is correctly placed to genus; and (6) †*Camponotus mengei* is a member of the crown Camponotini.

In contrast, we find that one generic and two tribal transfers are necessary. †*Pseudolasius boreus* and †*Glaphyromyrmex oligocenicus*, both attributed to Lasiini, are recovered outside of the tribe with very high support across all combined-evidence analyses (0.98–0.99 PP). We transfer the former species to a new genus forming †*XXX boreus* **gen. nov. comb. nov.**, which we consider *incertae sedis* within the subfamily (**tribal transfer**), and we transfer †*Glaphyromyrmex* back to the Formicini (**tribal transfer**; see *Taxonomy* below for discussion). With respect to *Lasius*, †*Lasius pumilus* is recovered outside of the genus, being consistently recovered as sister to *XXX* (Figs. 4, S7–S9); for this reason, we transfer the species to a third and final new genus, forming †*XXX pumilus* **gen. nov. comb. nov.**, which we place in the *XXX* genus group (see *Taxonomy* below for diagnosis).

The placements of two key taxa, †*L. schiefferdeckeri* and †*Kyromyrma neffi*, were sensitive to the choice of clock model. Under IGR (Fig. S7), †*L. schiefferdeckeri* was sister to the *Lasius* genus group (0.90 PP) and †*Kyromyrma* was nested in the crown Formicinae to the exclusion of the Myrmelachistini (0.94 PP). Under TK02 (Figs. S8, S9), †*L schiefferdeckeri* formed a clade with *Lasius* (0.63–0.71 PP) and †*Kyromyrma* was nested in the crown Lasiini (0.91–0.94 PP). Notably, the MCMC performed better under the IGR model relative to the TK02 model. Under TK02, the clock rate parameter sampled poorly, which was reflected in low ESS values for tree height and tree length. This, in addition to the unusually old estimations for the root node (Figs. S8 & S9), led us to prefer the IGR model over TK02. Regardless, we conservatively retain †*L. schiefferdeckeri* in *Lasius* and treat †*Kyromyrma* as *incertae sedis* within the Formicinae, in agreement with Ward et al. (2016) and Grimaldi & Agosti (2000).

### Evolutionary History

Results in this section pertain to *Questions 4* and *5* of Table 1.

#### Question 4: Where and when did the internal clades of the Lasiini originate?

Answer: The Lasiini likely originated and initially diversified in the Eastern Hemisphere, with modern genera present by the mid-Miocene.

Our primary divergence dating findings, based on the FBD analysis with and IGR branch length variation model, are as follows (Figs. 4, 5, Table 5): (1) The deepest split in the Lasiini (*Cladomyrma* + core Lasiini) is Late Cretaceous to Paleocene; (2) the split between *XXX* and the *Prenolepis* genus group crown was probably post-K/Pg and may have been during the Early Eocene Climatic Optimum (part of the Ypresian Epoch, 56.0–47.8 Ma); and (3) the crown *Lasius* genus group is young compared to that of the *Prenolepis* genus group, being probably Oligocene to Early Miocene in origin, while the crown *Prenolepis* genus group arose in the Late Eocene. Our date estimate for *Myrmecocystus* (14.6 mya) concurs with a recent study by van Elst et al. (2021) (14.1 mya in that study), but the crown *Lasius* genus group (24.9 mya), and the crown *Lasius* (21.9 mya) ages are slightly older (18.4 mya and 16.2 mya respectively in van Elst et al. 2021) but fall within the 95% HPDs estimated in that study. Our 95% HPD for *Nylanderia*, however, does not overlap with that of Williams *et al*. (2020), with our range being much younger (∼15–30 vs. ∼45–35 mya). With respect to our biogeographic reconstructions, we find that DIVALIKE is the most likely model for the deep history of the Lasiini (Table 5) although the second-most likely model (DEC) results in largely congruent estimated distributions (see results on Dryad https://doi.org/10.25338/B8B645). Mapping the DIVALIKE inference onto our chronogram and considering the climatological history of Earth (Fig. 5), we propose the following narrative.

**Table 5.** Results of divergence dating and historical biogeography reconstruction analyses. Divergence dating compares 95% HPDs for selected nodes given alternative calibration treatments. All ages reported are in units of mya and are for crown nodes. Biogeography reconstruction provides ranges at key nodes along with their likelihood (I = Indomalayan, Na = Nearctic, P = Palearctic; combinations indicate both regions were occupied).

| node | divergence dating |  |  |  |  |  |  |  | biogeography reconstruction |  |  |  |  |  |
| --- | --- | --- | --- | --- | --- | --- | --- | --- | --- | --- | --- | --- | --- | --- |
|  | FBD IGR clock |  | FBD TK02 clock |  | uniform IGR clock |  | uniform TK02 clock |  | DEC (lnL<br>-136.77) |  | DIVALIKE (lnL<br>-132.24) |  | BAYAREALIKE<br>(lnL -150.98) |  |
|  | mean age (95% HPD) | PP | mean age (95% HPD) | PP | mean age (95% HPD) | PP | mean age (95% HPD) | PP | range | likelihood | range | likelihood | range | likelihood |
| Lasini | **70.0 (57.6-82.7)** | **0.99** | 97.2 (89.6-106.0) | 0.91 | 80.8 (68.6-91.8) | 0.96 | 100.0 (90.9-112.3) | 0.94 | PI | 0.33 | PI | **0.37** | PI | 0.43 |
| core Lasini | **62.3 (51.2-72.9)** | **0.94** | 88.6 (79.8-97.8) | 0.64 | 72.5 (59.0-85.4) | 0.94 | 92.0 (83.5-102.2) | 0.49 | P | 0.42 | P | **0.56** | PI | 0.43 |
| XXX + *Prenolepis* g. grp. | **51.1 (42.2-60.2)** | **0.99** | 71.0 (60.8-81.9) | 1.00 | 59.9 (48.7-72.8) | 0.99 | 75.6 (65.1-86.5) | 1.00 | P | 0.40 | P | **0.53** | PI | 0.48 |
| *Prenolepis* g. grp. | **39.0 (36.5-50.9)** | **0.47** | 53.1 (43.6-62.6) | 1.00 | 50.9 (40.8-61.3) | 1.00 | 57.0 (48.2-66.5) | 1.00 | PI | 0.42 | PI | **0.50** | PI | 0.64 |
| *Lasius* g. grp. | **24.9 (18.0-33.0)** | **1.00** | 67.9 (54.9-81.6) | 0.99 | 31.1 (22.4-40.5) | 0.99 | 73.7 (61.9-88.3) | 0.99 | NaP | 0.53 | Na | **0.55** | NaP | 0.84 |
| crown *Lasius* | **21.9 (15.3-28.6)** | **0.77** | 63.7 (52.4-75.1) | 0.63 | 27.3 (19.4-35.6) | 0.73 | 64.3 (58.4-82.2) | 0.71 | P | 0.72 | P | **0.71** | NaP | 0.87 |
| *niger* clade | **15.8 (10.0-21.6)** | **1.00** | 53.2 (41.2-65.3) | 0.70 | 20.0 (12.7-27.7) | 1.00 | 60.6 (46.1-73.0) | 0.56 | P | 0.87 | P | **0.99** | NaP | 0.62 |
| *flavus* clade | **15.7 (10.4-21.0)** | **1.00** | 49.1 (36.7-61.8) | 1.00 | 19.7 (13.2-26.8) | 1.00 | 55.9 (41.1-69.4) | 1.00 | Na | 0.71 | Na | **0.90** | NaP | 0.91 |
| *Myrmecocystus* | **14.6 (7.3-21.5)** | **1.00** | 47.9 (33.5-63.1) | 1.00 | 17.8 (8.9-26.9) | 1.00 | 55.8 (40.0-69.8) | 1.00 | Na | 0.87 | Na | **0.97** | Na | 0.53 |

**Fig. 5.**
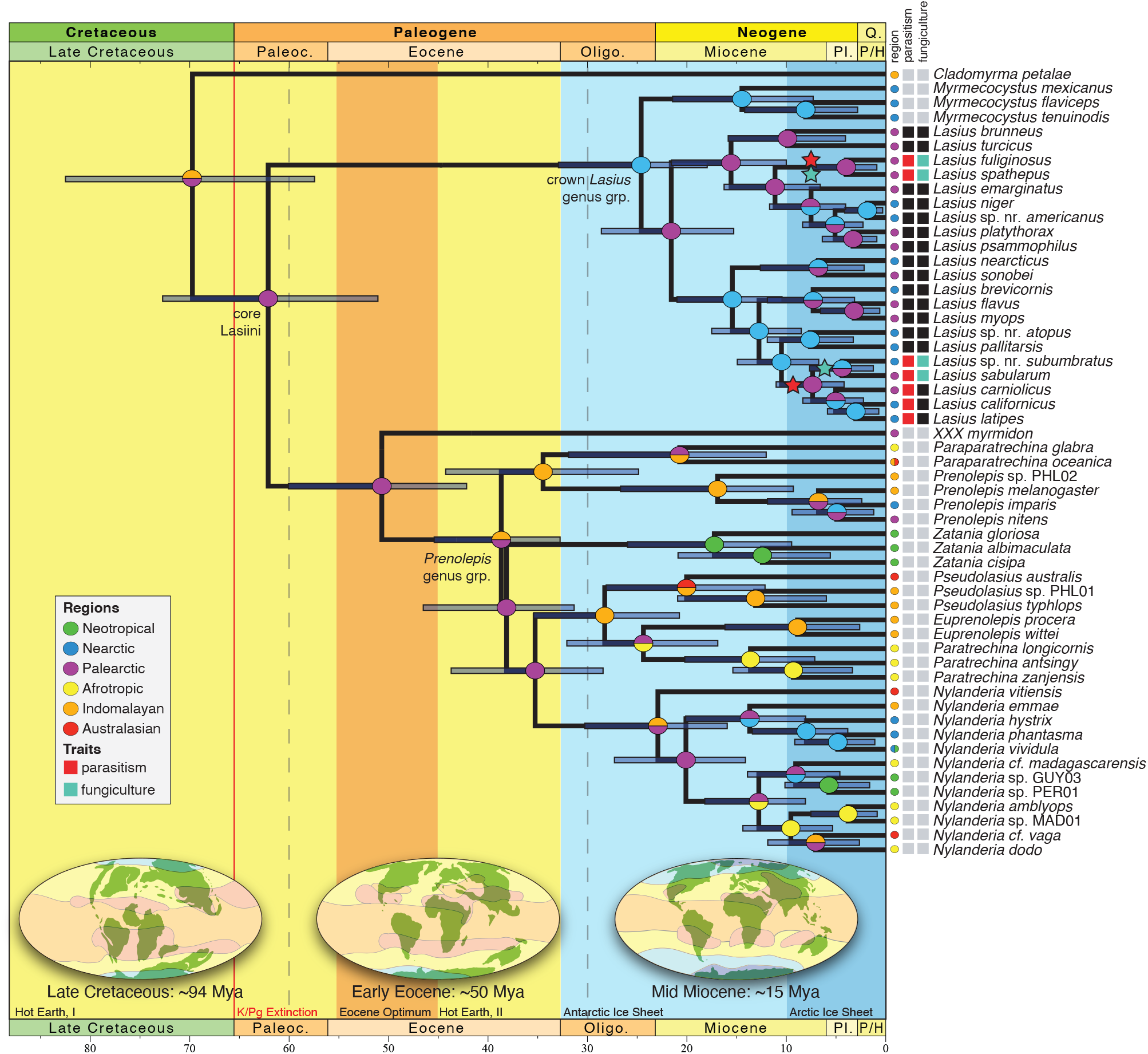
Results of the BioGeoBEARS and life history trait ancestral state estimation analyses on a chronogram pruned from the analysis of the Lasiini_w_outgroups_w_fossils_154t combined morphological and molecular dataset. Horizontal blue bars at nodes are 95% highest posterior density (HPD) intervals from the divergence dating. Colored circles on nodes are inferred inherited biogeographic ranges, with the top half of each corresponding to the upper branch and bottom half with the lower branch; colored circles at branch tips represent biogeographical coding for terminal; dashed lines at 60 and 30 mya indicate the boundaries of the three BioGeoBEARS dispersal rate matrix time periods. Colored boxes at branch tips represent presence or absence of two traits, in *Lasius*: temporary social parasitism (black = absence, red = presence) and fungiculture (black = absence, green = presence). Stars on internal branches indicate the evolution of temporary social parasitism (red) and fungiculture (green) The figure background indicates generalized major climatological regimes during Earth’s history (Royer *et al*. 2004; Zachos *et al*. 2008; Hansen *et al*. 2013), while the inset globes visually represent paleoclimate and landmass configuration modified from Scotese (2010).

The Lasiini originated on the Eurasian landmass around the End Cretaceous, with the core Lasiini originating in the Western or “Palearctic” portion of the ancient continent. Subsequently, the *Lasius* genus group experienced some degree of extinction which pruned the clade down to a single Nearctic ancestor, possibly due to the global cooling of the Oligocene to Pleistocene. Within the crown *Lasius* genus group, *Myrmecocystus* arose as an arid-region-adapted lineage from an extinct *Lasius*-like lineage, as evinced by the ∼35 mya Baltic *Lasius* fossils. The ancestor of *Lasius* dispersed back to the Palearctic, and a complicated series of range expansions and dispersal events occurred throughout the Miocene and Pliocene. An early dispersal event to the Nearctic from the Palearctic led to the origin of the distinctive *flavus* clade, wherein a complex series of dispersal events between the Palearctic and the Nearctic probably occurred across the Bering Land Bridge. Within the *niger* clade, there was at least one dispersal to the Nearctic in the *niger* group.

Whereas the *Lasius* genus group was heavily pruned, the *Prenolepis* genus group began radiating during the second half of the Eocene, with their depauperate sister group surviving to the present day as *XXX* **gen. nov.** The ancestor of the *Prenolepis* genus group either adapted to, or retained an ancestral physiological optimization for tropical climes, having perhaps expanded into the Indomalayan region following the Eocene Optimum. Subsequently, the *Prenolepis* genus group radiated across the global tropics through a complicated history of transcontinental dispersal. Notably, the crown lineages of the *Lasius* genus group have failed to invade tropical zones, whereas at least two invasions of the temperate zone have occurred in the *Prenolepis* genus group. In our analyses, recovered those for the *imparis* clade of *Prenolepis*, and the *vividula* clade of *Nylanderia*. A third invasion is implied by Matos-Maraví *et al*. 2018 for the *Nylanderia fulva* clade. Future studies of the biogeography of *Lasius* may benefit from treating biogeographic region as an observed state (character) in the non-genomic (phenotypic) matrix to extract information from the coarse compression fossils in the Nearctic and Palearctic regions.

#### Question 5: What was the evolutionary sequence of formicine morphology and life history traits?

Answer: Our analyses recover wide-set hind coxae and sepalous proventriculi as synapomorphies of the Formicinae, and show independent, convergent evolution of social parasitism and fungiculture in *Lasius*.

We determined that the polarity of six characters is particularly important for understanding the morphological evolution of the Lasiini and the subfamily Formicinae. In all cases, the equal rates (ER) transition matrix was found to be the most likely model (Table 6). Empirical Bayesian posterior probabilities are presented for all character states at selected nodes in Table 7, and mapped posterior probabilities are presented in Figs. S10–15. Based on our analyses, we predict that the most recent common ancestor of the Formicinae probably had, in addition to the acidopore, the following states: **(1)** Eyes set at or anterior to head midlength [plesiomorphy]; **(2)** wide-set hind coxae [apomorphy]; **(3)** absence of a transverse sulcus posterad the helcial sternite [plesiomorphy]; **(4)** circular propodeal spiracles [plesiomorphy]; **(5)** broadly shouldered tergosternal margins laterad the helcium [plesiomorphy]; and **(6)** a sepalous proventriculus [apomorphy]. Roughly corresponding to expectation (Bolton 2003), slit-shaped spiracles are estimated to be defining apomorphies of the Formicini, Camponotini, and the melophorines *Melophorus* and *Notostigma* (Fig. S12); other results require further consideration.

**Table 6.** Results of ANOVA tests on alternative ancestral state estimation models.

| model | Log lik. | Df | Df change | Resid. Dev | Pr(>\|Chi\|) |
| --- | --- | --- | --- | --- | --- |
| Eyes_ER | -49.43 | 1 | NA | NA | NA |
| Eyes_ARD | -49.12 | 2 | 1 | 0.63 | 0.43 |
| Coxae_ER | -26.97 | 1 | NA | NA | NA |
| Coxae_ARD | -26.24 | 2 | 1 | 1.48 | 0.22 |
| Prov_ER | -27.56 | 1 | NA | NA | NA |
| Prov_ARD | -27.29 | 2 | 1 | 0.53 | 0.47 |
| Spiracle_ER | -28.83 | 1 | NA | NA | NA |
| Spiracle_ARD | -27.64 | 2 | 1 | 2.38 | 0.12 |
| Sulcus_ER | -12.74 | 1 | NA | NA | NA |
| Sulcus_ARD | -12.25 | 2 | 1 | 0.97 | 0.32 |
| AIII_ER | -42.53 | 1 | NA | NA | NA |
| AIII_SYM | -39.71 | 3 | 2 | 5.66 | 0.06 |
| AIII_ARD | -38.36 | 6 | 3 | 2.69 | 0.44 |
| Parasitism_ER | -7.74 | 1 | NA | NA | NA |
| Parasitism_ARD | -7.39 | 2 | 1 | 0.71 | 0.40 |
| Fungiculture_ER | -7.75 | 1 | NA | NA | NA |
| Fungiculture_ARD | -7.73 | 2 | 1 | 0.03 | 0.87 |

**Table 7.**
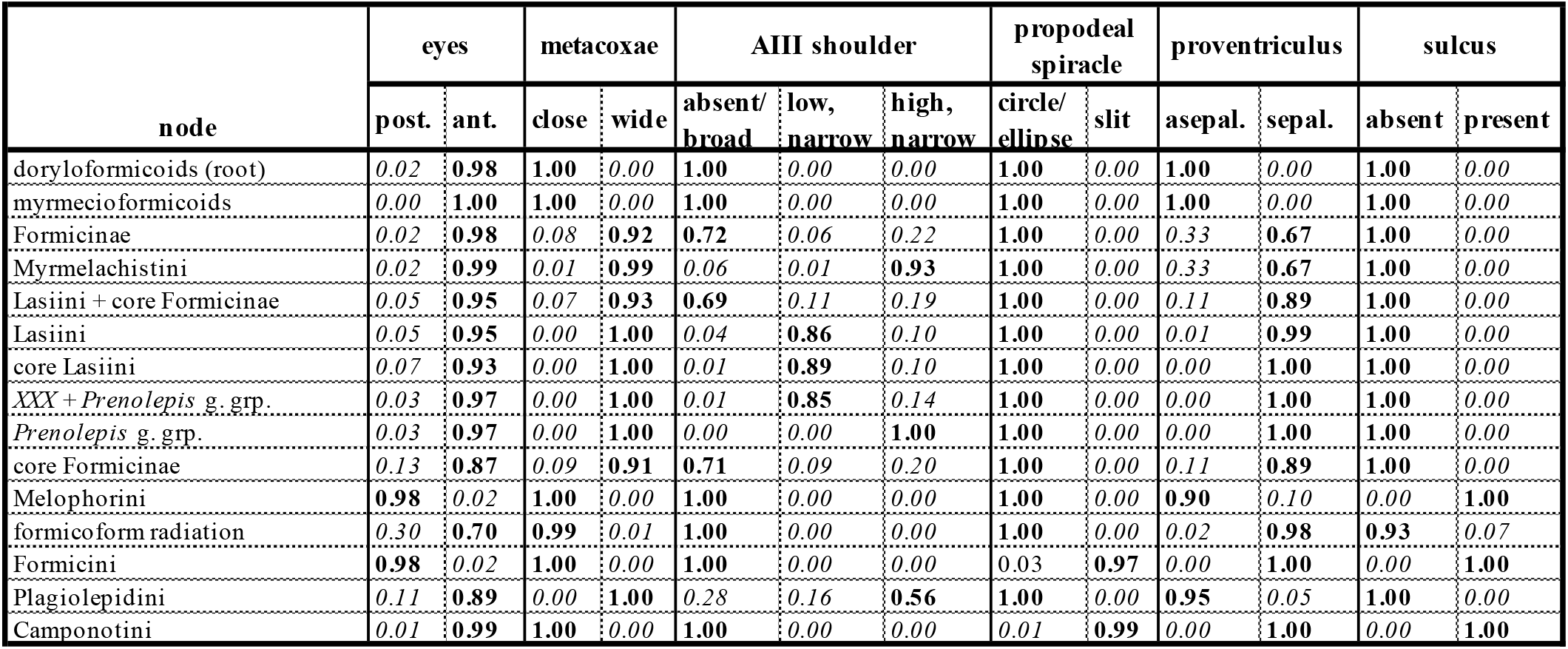
Empirical Bayesian posterior probabilities from ancestral state estimation for selected nodes; numbers may not add up to 1.00 due to rounding error. For ease of reference, the highest probability state is **bolded** and lowest *italicized* for each node.

| node | eyes |  | metacoxae |  | AIII shoulder |  |  | propodeal spiracle |  | proventriculus |  | sulcus |  |
| --- | --- | --- | --- | --- | --- | --- | --- | --- | --- | --- | --- | --- | --- |
|  | post. | ant. | close | wide | absent/<br>broad | low,<br>narrow | high,<br>narrow | circle/<br>ellipse | slit | asepal. | sepal. | absent | present |
| doryloformicoids (root) | 0.02 | 0.98 | 1.00 | 0.00 | 1.00 | 0.00 | 0.00 | 1.00 | 0.00 | 1.00 | 0.00 | 1.00 | 0.00 |
| myrmecioformicoids | 0.00 | 1.00 | 1.00 | 0.00 | 1.00 | 0.00 | 0.00 | 1.00 | 0.00 | 1.00 | 0.00 | 1.00 | 0.00 |
| Formicinae | 0.02 | 0.98 | 0.08 | 0.92 | 0.72 | 0.06 | 0.22 | 1.00 | 0.00 | 0.33 | 0.67 | 1.00 | 0.00 |
| Myrmelachistini | 0.02 | 0.99 | 0.01 | 0.99 | 0.06 | 0.01 | 0.93 | 1.00 | 0.00 | 0.33 | 0.67 | 1.00 | 0.00 |
| Lasiini + core Formicinae | 0.05 | 0.95 | 0.07 | 0.93 | 0.69 | 0.11 | 0.19 | 1.00 | 0.00 | 0.11 | 0.89 | 1.00 | 0.00 |
| Lasiini | 0.05 | 0.95 | 0.00 | 1.00 | 0.04 | 0.86 | 0.10 | 1.00 | 0.00 | 0.01 | 0.99 | 1.00 | 0.00 |
| core Lasiini | 0.07 | 0.93 | 0.00 | 1.00 | 0.01 | 0.89 | 0.10 | 1.00 | 0.00 | 0.00 | 1.00 | 1.00 | 0.00 |
| XXX + *Prenolepis* g. grp. | 0.03 | 0.97 | 0.00 | 1.00 | 0.01 | 0.85 | 0.14 | 1.00 | 0.00 | 0.00 | 1.00 | 1.00 | 0.00 |
| *Prenolepis* g. grp. | 0.03 | 0.97 | 0.00 | 1.00 | 0.00 | 0.00 | 1.00 | 1.00 | 0.00 | 0.00 | 1.00 | 1.00 | 0.00 |
| core Formicinae | 0.13 | 0.87 | 0.09 | 0.91 | 0.71 | 0.09 | 0.20 | 1.00 | 0.00 | 0.11 | 0.89 | 1.00 | 0.00 |
| Melophorini | 0.98 | 0.02 | 1.00 | 0.00 | 1.00 | 0.00 | 0.00 | 1.00 | 0.00 | 0.90 | 0.10 | 0.00 | 1.00 |
| formicoform radiation | 0.30 | 0.70 | 0.99 | 0.01 | 1.00 | 0.00 | 0.00 | 1.00 | 0.00 | 0.02 | 0.98 | 0.93 | 0.07 |
| Formicini | 0.98 | 0.02 | 1.00 | 0.00 | 1.00 | 0.00 | 0.00 | 0.03 | 0.97 | 0.00 | 1.00 | 0.00 | 1.00 |
| Plagiolepidini | 0.11 | 0.89 | 0.00 | 1.00 | 0.28 | 0.16 | 0.56 | 1.00 | 0.00 | 0.95 | 0.05 | 1.00 | 0.00 |
| Camponotini | 0.01 | 0.99 | 1.00 | 0.00 | 1.00 | 0.00 | 0.00 | 0.01 | 0.99 | 0.00 | 1.00 | 0.00 | 1.00 |

Of the reconstructed states, the sepalous form of the proventriculus is newly detected as a synapomorphy of the subfamily—if the asepalous state in *Myrmelachista* is interpreted as an independent reversal, as implied by our results (Fig. S14). This finding contradicts the traditional assumption that the sepalous form arose from the asepalous form within the Formicinae (Emery 1925, Eisner 1957), and hypothesis that the sepalous form is homoplastic within the subfamily (Agosti 1991, Bolton 2003). Because our coding was generalized from the literature, our results must be treated as tentative. We strongly recommend that future studies expand on the sampling of Eisner (1957), particularly to include Leptanillinae, Martialinae, and more poneroids (*sensu* Moreau *et al*. 2006, Brady *et al*. 2006), and to define new traits of the proventriculus for evolutionary analysis because of its “social stomach” function (Eisner & Brown 1958, Wilson 1971). Moreover, a microanatomical study, perhaps using computed tomography, would be a substantial contribution to this question.

Within the Formicinae, posteriorly set eyes are estimated to have been derived nine independent times (Fig. S10), which is a simplistic statement, given that eye position is a continuous and evolutionarily labile trait. This strongly contradicts our intuition that posteriorly set eyes were ancestral, as they are observed in the Leptanillinae (where known), Amblyoponinae, and stem Formicidae (Perrichot *et al*. 2020; Boudinot *et al*. 2020). If posteriorly set eyes are truly retained in the various Formicinae, the Ectatomminae *s. l.*, and the Pseudomyrmecinae, then we must appreciate the low likelihood of this scenario and search for corroborating evidence. Ideally, eye position should be modeled as a continuous trait (Parins-Fukuchi 2018).

Our estimates indicate that wideset (“lasiine”) coxae are an apomorphy of the Formicinae, with a subsequent reversal in the formicoform radiation (the clade that includes Camponotini, Formicini, and Melophorini, along with the monotypic tribes Gesomyrmecini, Gigantiopini, Oecophyllini, and Santschiellini (Fig. S11), with losses in *Lasiophanes* and the ancestor of *Teratomyrmex* and *Prolasius* of the Plagiolepidini, as well as *Myrmoteras*. It may be possible to evaluate the plausibility of this reconstruction through comparative µ-CT analysis of propodeal / petiolar skeletomuscular anatomy. At present, comparative data on “waist” skeletomusculature is limited (*e.g.*, Hashimoto 1996, Perrault 2004).

We detect three independent origins of the “plagiolepidiform” third abdominal segment (*i.e.*, raised, with free sclerites occurring high): Once in the ancestor of the Myrmelachistini, once within the Lasiini (*Prenolepis* genus group), and once in the Plagiolepidini (Fig. S15). Our results support Bolton’s (2003) hypothesis that the lasiine condition (*i.e.*, narrowly shouldered but not raised) was a precursor to the plagiolepidiform condition, as elevated shoulders arose well-within the Lasiini. However, whether the narrow and low shoulders of *Anoplolepis* (Plagiolepidini) are ancestral or a reversal (as observed for *Acropyga*) is uncertain. Narrowed shoulders arose three times within the Melophorini (*Stigmacros, Lasiophanes,* and the ancestor of *Teratomyrmex* + *Prolasius*). Curiously, “high and narrow shoulders” are also observed in Dolichoderinae (*e.g.*, Bolton 1994), thus comparative anatomical studies focusing on this condition are warranted.

Finally, most of the socially parasitic species within *Lasius* included here (“nr. *subumbratus*”, *sabularum*, *carniolicus*, *californicus*, *latipes*) form a strongly-supported monophyletic group within the *flavus* clade (Fig. 2). The other two socially parasitic species, *fuliginosus* and *spathepus*, are in the *niger* clade and are sister to one another. A subset of social parasites, here represented by “nr. *subumbratus*”, *sabularum*, *fuliginosus*, and *spathepus* are also known to culture fungi to bind walls of their nests (Schlick-Steiner *et al*. 2008). These species are widely separated on the phylogeny, and it appears likely that both social parasitism and fungiculture evolved at least twice within the genus (Fig. 5, S16, S17).

### Taxonomy

Here we report the taxonomic implications of our study (*Question 6* of Table 1). Note that probable autapomorphies in the context of the Lasiini are *italicized* in the taxon definitions.

**Tribe Lasiini Ashmead, 1905**

(Figs. 6, 7, 8A–K,9A–J, 10A–D)

**Fig. 6.**
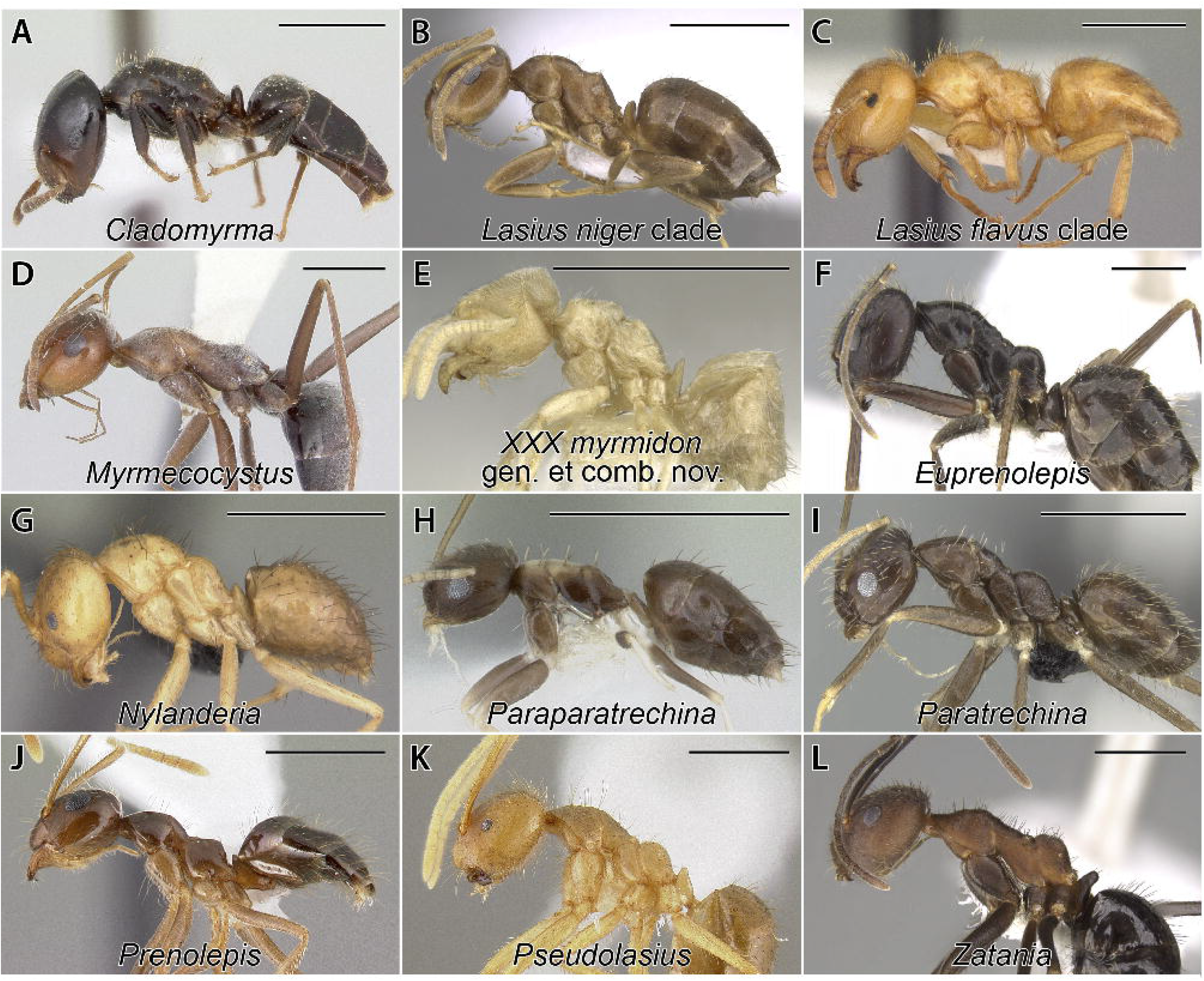
Gross phenotypic synopsis of the Lasiini; all images in profile view; scale bars = 1.0 mm. A, *Cladomyrma hewitti* (CASENT0173906); B, *Lasius lasioides* (CASENT0906077); C, *Lasius citrinus* (CASENT0103542); D, *Myrmecocystus melliger* (CASENT0103518); E, *XXX myrmidon* (CASENT0903666); F, *Euprenolepis procera* (CASENT0906260); G, *Nylanderia amblyops* (CASENT0007735); H, *Paraparatrechina albipes* (CASENT0178759); I, *Paratrechina ankarana* (CASENT0454372); J, *Prenolepis imparis* (CASENT0179615); K, *Pseudolasius amaurops* (CASENT0106005); L *Zatania gibberosa* (CASENT0281461). (Image credits, AntWeb: A, C, G, H = April Nobile; B = Shannon Hartman; D = Jen Fogarty; E, Will Ericson; F, Estella Ortega; I, Michele Esposito; J, Erin Prado; K, Michael Branstetter; L, Ziv Lieberman).

**Fig. 7.**
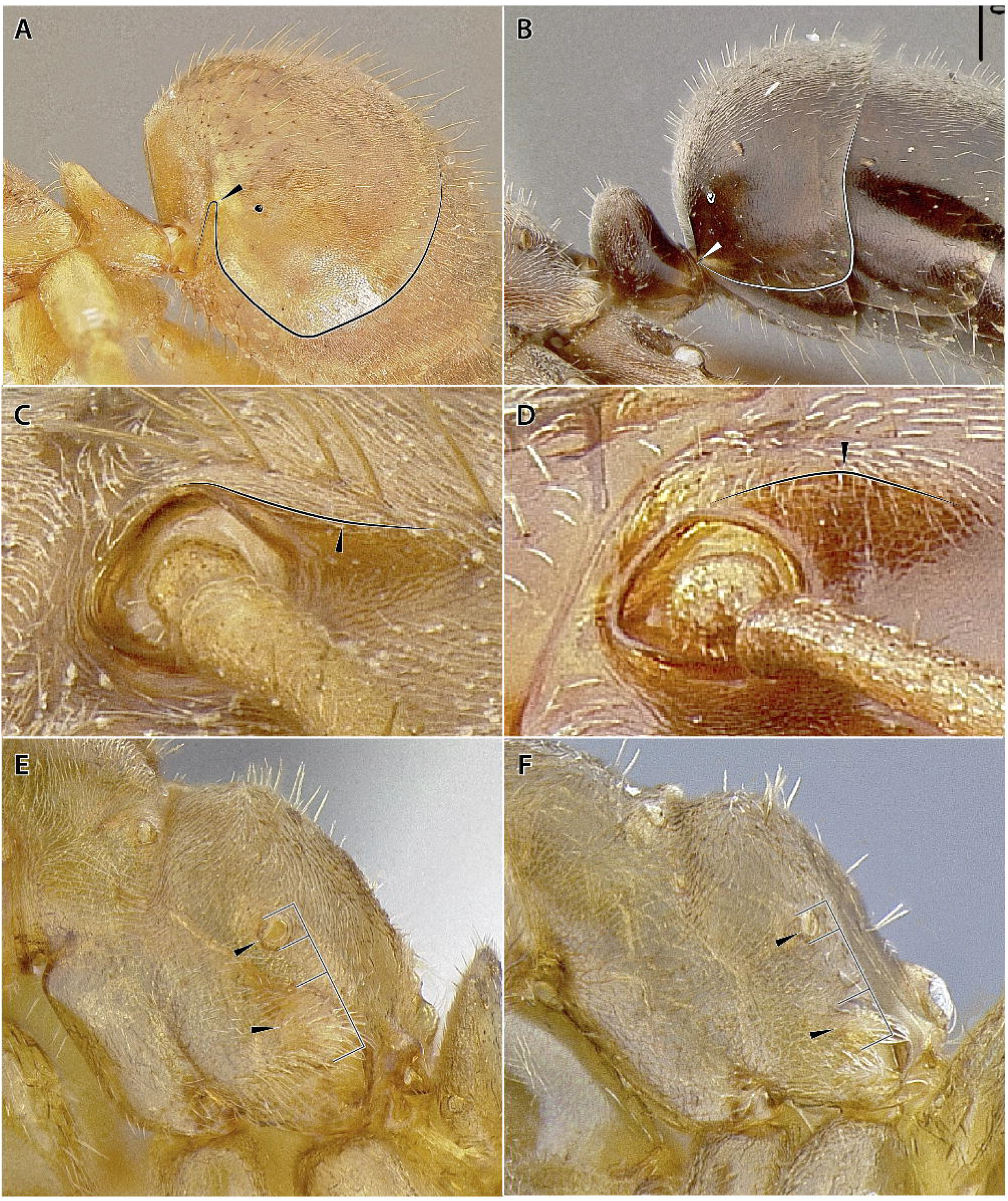
Some key features of the major clades of the Lasiini. A, third abdominal tergite modified with dorsoventrally-oriented groove for reception of entire petiole (*Pseudolasius breviceps*); black line indicates contour of tergum, and black ellipse the spiracle; B, third abdominal tergite receiving only posterior base of petiole (*Myrmecocystus mimicus*); white line indicates contour of tergum, and white ellipse the spiracle; C, frontal protuberance and effaced carinae low relative to face (*Pseudolasius breviceps*); D, frontal protuberance and effaced carinae raised dorsally from face (*Myrmecocystus mimicus*); E, metapleural gland atrium grossly enlarged (*Lasius subumbratus*); F, metapleural gland atrium small (*Lasius turcicus*).

**Fig. 8.**
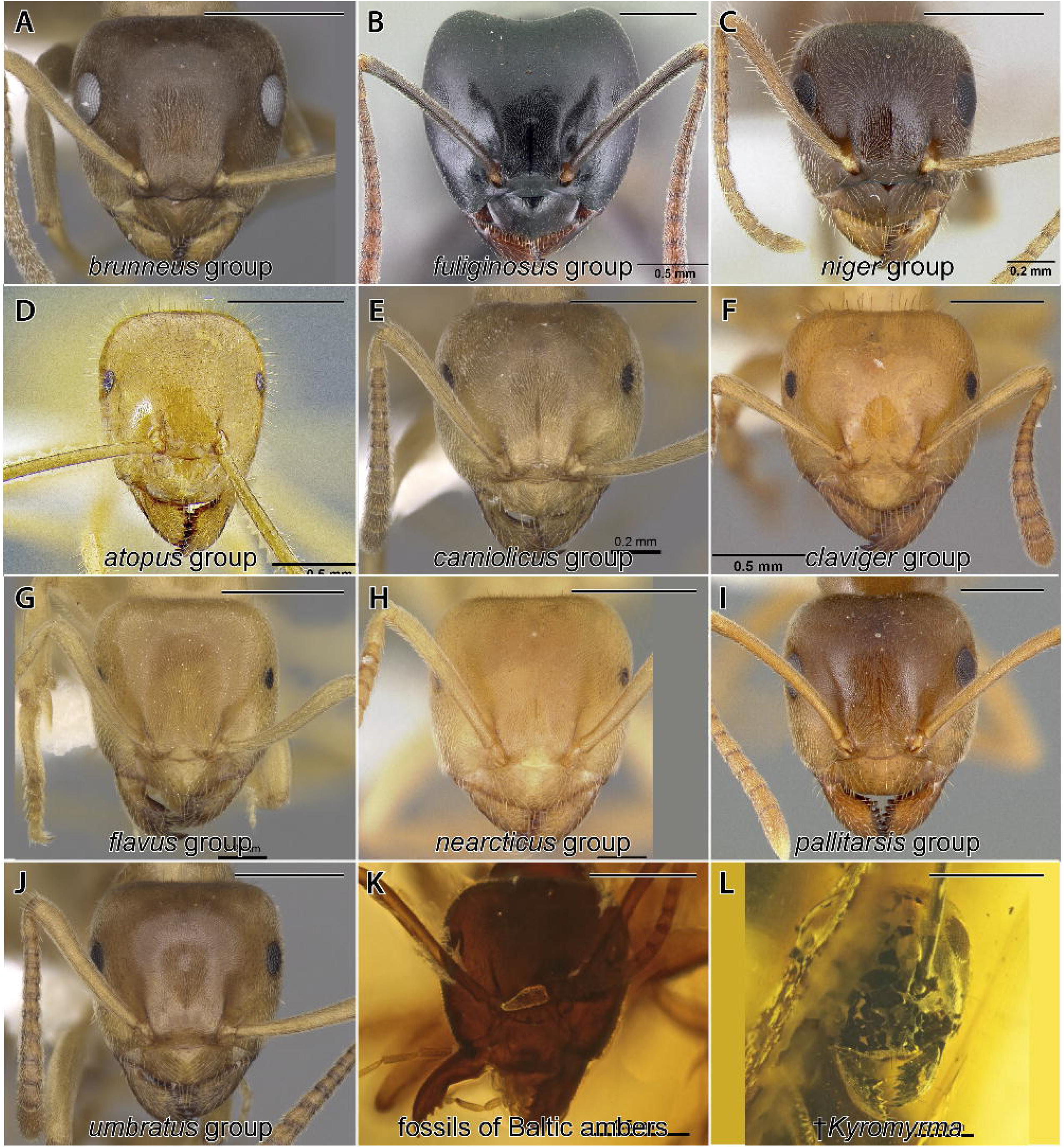
Gross phenotypic synopsis of the extant species groups of *Lasius*, plus two phenotypically similar fossil species; all images in full-face view; scale bars = 0.5 mm. A, *Lasius brunneus* (CASENT0280440); B, *Lasius fuliginosus* (CASENT0179898); C, *Lasius* nr. *niger* (CASENT0106128); D, *Lasius* nr. *atopus* (CASENT0234858); E, *Lasius carniolicus* (CASENT0280471); F, *Lasius claviger* (CASENT0103542); G, *Lasius brevicornis* (CASENT0280456); H, *Lasius nearcticus* (CASENT0104774); I, *Lasius pallitarsis* (CASENT0005405); J, *Lasius aphidicola* (CASENT0280468); K, †*Lasius schiefferdeckeri* (MNHNB25217); L, †*Kyromyrma neffi* (AMNH-NJ1029). (Image credits: D = the authors; AntWeb: A = Will Ericson; B = Erin Prado; C = Michael Branstetter; E, G, J = Shannon Hartman; F, H = April Nobile; I = unknown; K = Vincent Perrichot; L = Dave Grimaldi & Vincent Perrichot.)

**Fig. 9.**
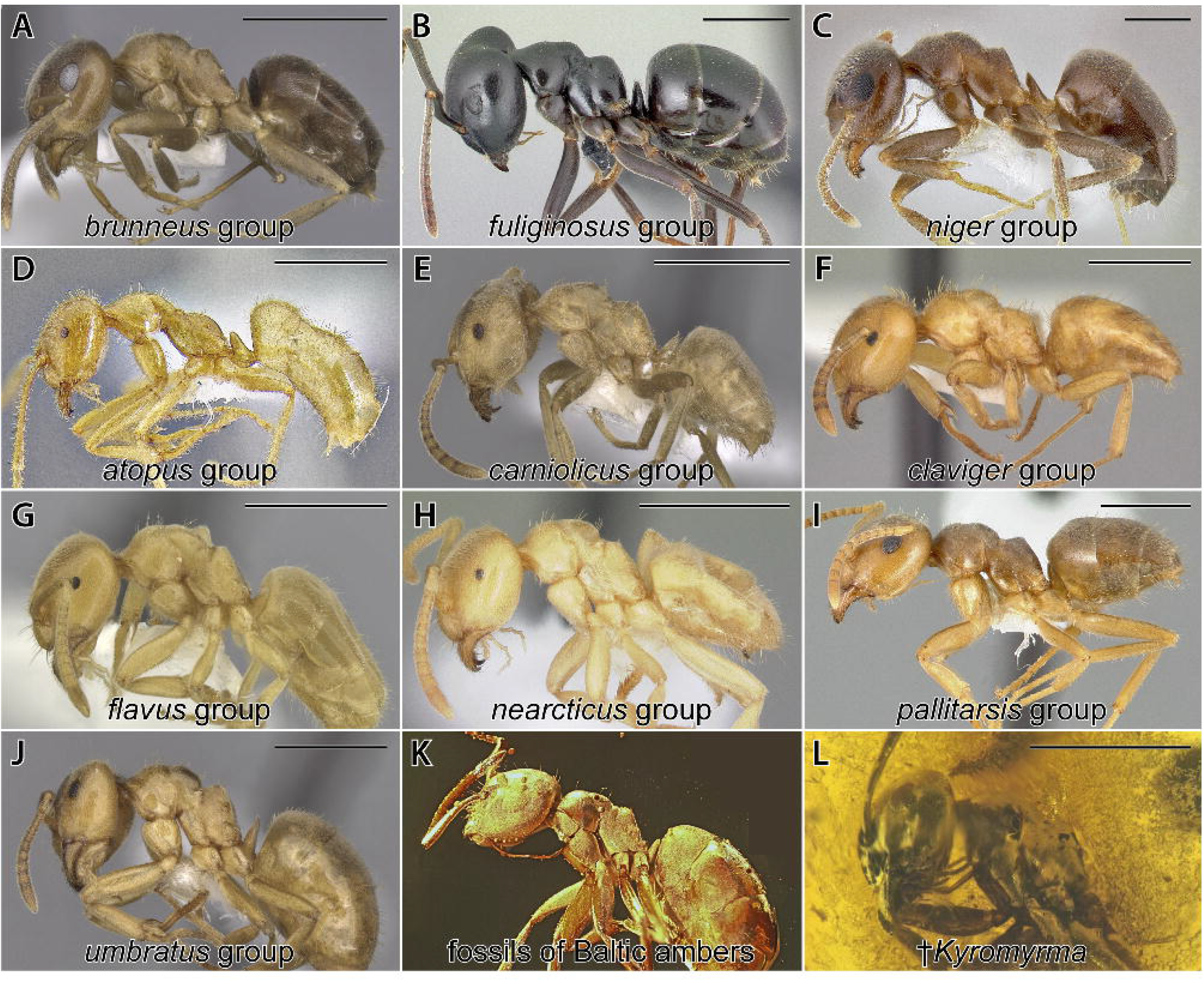
Gross phenotypic synopsis of the species groups of *Lasius*, plus two phenotypically similar fossil species; all images in profile view; scale bars = 1.0 mm. A, *Lasius brunneus* (CASENT0280440); B, *Lasius fuliginosus* (CASENT0179898); C, *Lasius* nr. *niger* (CASENT0106128); D, *Lasius* nr. *atopus* (CASENT0234858); E, *Lasius carniolicus* (CASENT0280471); F, *Lasius claviger* (CASENT0103542); G, *Lasius brevicornis* (CASENT0280456); H, *Lasius nearcticus* (CASENT0104774); I, *Lasius pallitarsis* (CASENT0005405); J, *Lasius aphidicola* (CASENT0280468); K, †*Lasius schiefferdeckeri* (HJF013); L, †*Kyromyrma neffi* (AMNH-NJ1029). (Image credits: D = the authors; AntWeb: A = Will Ericson; B = Erin Prado; C = Michael Branstetter; E, G, J = Shannon Hartman; F, H = April Nobile; I = unknown; K = Vincent Perrichot; L = Dave Grimaldi & Vincent Perrichot.)

**Type genus:** *Lasius*.

**Included genus groups:** *Cladomyrma*, *Lasius* group, *XXX* group, *Prenolepis* group.

**Definition (worker):**

1. With characters of Formicinae (see Bolton 2003 and note 1).
2. Mandible triangular, with 4–11 teeth; third tooth from apex reduced.
3. Dorsal mandibular groove, when present, running along lateral mandibular margin as seen in dorsal view (note 2).
4. Palp formula usually 6,4, less often 3,4.
5. Frontal protuberance mediad antennal toruli with frontal carinae effaced, becoming broadly rounded posteriorly until continuous with face (note 3).
6. Frontal protuberance raised above toruli or not.
7. Antennal toruli near or abutting posterior clypeal margin.
8. Antenna 12-, 11-, or 8-merous.
9. Compound eyes not enormous, may be reduced to absent; long axis of eyes subparallel.
10. Ocelli present or absent (note 4).
11. Metanotum differentiated or not.
12. Metapleural gland present, dorsal rim of metapleural gland curved inward.
13. Propodeal spiracle at or near posterolateral margin of propodeum.
14. Propodeal spiracle circular to elliptical, not slit-shaped.
15. Metacoxae wideset: Distance between mesocoxal bases less than between that between bases of metacoxae with mesosoma in ventral view and coxae oriented at right-angles to long axis of body.
16. Metatibiae without double row of ventral (inner) setae.
17. Petiolar foramen in profile view low, not to barely exceeding dorsal margin of metapleural gland, with or without dorsal margin or lip, but lip, when present, inconspicuous (note 5).
18. Petiolar foramen in ventral view long, anterior margin exceeding metacoxal cavities anteriorly.
19. Petiolar node conspicuously shorter than propodeum, not reaching dorsal surface of propodeum (note 6).
20. Petiolar apodeme (situated anteriorly on tergum) contiguous or nearly contiguous with petiolar node (note 7).
21. Petiolar sternum U-shaped in cross-section (note 8).
22. Abdominal segment III transverse sulcus absent.
23. Base of abdominal segment III with or without complete tergosternal fusion lateral to helcium, free sclerites commencing distantly up segment or near helcium; tergum III overhanging petiole or not.
24. Proventriculus sepalous (note 9).

#### Notes on definition

*Note 1*. The generic composition of the Lasiini has recently been reassessed following Blaimer *et al*. (2015) and Ward *et al*. (2016). The tribal definition proposed here is thus revised relative to Bolton (2003). It includes new characters and recognizes that several former plagiolepidines belong in the Lasiini.

*Note 2*. Previous studies have not focused attention on the dorsal mandibular groove. Here it was observed that the groove, as seen with the mandible in dorsal view, runs along the outer margin of the mandible toward the mandibular apex in the Myrmelachistini, Lasiini, and most Melophorini. This contrasts with the state observed in the “formicoform radiation” (*i.e.*, the clade that includes Camponotini, Formicini, and Melophorini, along with the monotypic tribes Gesomyrmecini, Gigantiopini, Oecophyllini, and Santschiellini; see the section †*Kyromyrma*, an ancestral formicine, under the section “Incertae sedis in the Formicinae” below). When the groove is present in the “formicoform radiation”, it is very close to the basal mandibular margin and is shortened, except in various plagiolepidines. The medial and shortened state may have arisen multiple times of the Formicini, Plagiolepidini, and Camponotini clade.

*Note 3*. The “frontal protuberance” is the region between the antennal toruli which is raised relative to the regions of the face laterad the toruli. To some degree, development of this condition accounts for the “laterally directed” torular condition of Formicidae (Boudinot *et al*. 2020). In other formicine tribes, including the Melophorini (with *Prolasius* as an exception), the frontal protuberance is carinate above the toruli (*i.e.*, “frontal carinae are present”), with these carinae continuing as a sharp ridge until their posterior terminus. Within the Lasiini, the frontal carinae are effaced; they are long in all groups except *Cladomyrma*, while in the Plagiolepidini the carinae are short in all genera except *Anoplolepis*.

*Note 4.* Ocellus presence / absence is a relatively weak character as expression of ocelli may be variable for genera in which ocelli are observed, such as *Lasius*. However, this statement is included as ocellus expression is a traditional character which is easy to evaluate and may have value for future works defining tribes wherein ocelli may be consistently present or absent. Within the Lasiini, ocelli are always absent in *Cladomyrma*, *Euprenolepis*, *Nylanderia* (although infrequently expressed as single, median ocellus), and *Pseudolasius*, variably present among *Lasius*, *Paraparatrechina*, and *Prenolepis* species, and consistently present in *Myrmecocystus*, *Paratrechina* (except *P. kohli*; see comments under *Prenolepis* genus group), *Zatania*.

*Note 5*. The conformation of the dorsal region of the petiolar foramen is newly described here. There is complex variation of the form across the subfamily, but it appears at least that the form observed in the Lasiini is consistent; this form is also observed in the Myrmelachistini. In various lineages within the core Formicinae, *i.e.*, the clade that spans the node between Melophorini and Camponotini, a raised and conspicuously carinate dorsal margin is observed.

*Note 6*. Short petiolar nodes are also observed in the Myrmelachistini and Plagiolepidini (excluding *Anoplolepis*). The node height is variable in the Melophorini, being short in *Prolasius* and *Myrmecorhynchus*.

*Note 7*. The apodeme is clearly separated from the node in most Melophorini, except *Prolasius*.

*Note 8*. A U-shaped petiolar sternum was used by Bolton (2003) to diagnose the lasiine tribe group. This trait also occurs in the Myrmelachistini, Myrmoteratini, Plagiolepidini, and four Melophorini (*Lasiophanes*, *Prolasius*, *Stigmacros*, *Teratomyrmex*); see “Ancestral state estimation” results above and Fig. S15.

*Note 9*. The form of the proventriculus forms a natural division between the Plagiolepidini and the “plagiolepidiform” *Prenolepis* genus group, as observed by Emery (1925). Due to uncertainty about the polarity of the sepalous condition—and whether this form arose multiple times (Eisner 1957; Agosti 1990, 1991)—Bolton (2003) lumped the two plagiolepidiform clades, plus the Myrmelachistini, into the Plagiolepidini.

#### Exclusion of †*Glaphyromyrmex* from the Lasiini

The Baltic amber formicine, †*Glaphyromyrmex* Wheeler, was placed in the Formicini until recently (Wheeler 1915, Donisthorpe 1943, Dlussky 1967, Dlussky & Fedoseeva 1988, Bolton 1994), when Dlussky (2008) transferred the genus to the Lasiini. Placement of †*Glaphyromyrmex* in the Lasiini is counterintuitive given presence of a ventral double-row of setae on its metatibiae, a state which does not occur in any lasiine. Intuitive placement of †*Glaphyromyrmex* based on morphology is challenging, however. †*Glaphyromyrmex* differs from members of all formicine tribes in which double seta rows occur (Formicini, Camponotini, some Melophorini). Specifically, †*Glaphyromyrmex* differs from the Formicini in eye position (eyes set at about head midlength), from the Camponotini in having antennal toruli which abut the posterior clypeal margin (vs. widely separated), and from the Melophorini, which have well-defined dorsal and ventral flaps surrounding the metapleural gland, a conformation that is apparently absent in †*Glaphyromyrmex*. Recognizing these differences, however, we transfer †*Glaphyromyrmex* back to the Formicini (**tribal transfer**) based on our combined analyses (Fig. 4) and recommend revised study of the fine-scale external anatomy of the fossil taxon.

#### On the identity of †*Protrechina*

Wilson (1985) described a genus putatively close to *Paratrechina sensu lato* (∼ *Prenolepis* genus group) from mid-Eocene Claiborne amber (Arkansas, 40.4–37.2 Ma; Saunders *et al*. 1974). This genus, †*Protrechina*, supposedly differs from *Paratrechina s. l.*, *Lepisiota*, and *Brachymyrmex*—among other, unstated formicines—by absence of standing macrosetae on the mesosomal dorsum, a state similar to that observed for †*XXX* and *XXX* as noted below. The genus has been variably treated as a lasiine (Bolton 1994, 1995; LaPolla & Dlussky 2010), a “prenolepidine” (Hölldobler & Wilson 1990), or as *incertae sedis* in the subfamily where it remains at present (Wilson 1985; Dlussky & Fedoseeva 1988; Bolton 2003; Ward *et al*. 2016). The type specimen of †*Protrechina carpenteri*, at the Museum of Comparative Zoology (Harvard), should be reexamined to evaluate its tribal, and perhaps generic, placement. This would be particularly valuable given the approximately mid-Eocene origin of the *Prenolepis* genus group here inferred (Fig. 4; Table 5); such a study should be facilitated by use of micro-CT (*e.g.*, Barden *et al*. 2017b; Hita-Garcia *et al*. 2017), given the poor condition of the specimen reported by LaPolla & Dlussky (2010).

#### Key to extant genera of Lasiini and primary clades of *Lasius* (workers)

**1.** Antenna with 8 antennomeres. Mesonotum, metanotum, and propodeum more-or-less continuous, and metanotum undifferentiated (Fig. 6A). Propodeal spiracle distinctly situated in dorsal third of propodeum. Frontal carinae inconspicuous, being extremely short (< 0.5 × anteroposterior antennal torulus diameter), or absent … ***Cladomyrma*** (see Agosti 1991, Agosti *et al*. 1999)

- Antenna with 11 or 12 antennomeres. Mesonotum, metanotum, and/or propodeum discontinuous and metanotum differentiated (Figs. 6A–G, I–L), rarely mesosomal dorsum continuous and metanotum undifferentiated (few *Paraparatrechina*) (Fig. 6H). Propodeal spiracle situated somewhat below or above midheight of propodeum, but not in dorsal third. Frontal carinae conspicuous (≥ 1.0 × anteroposterior antennal torulus diameter) … **2**

**2.** Petiole posteriorly elongated. Node reduced, apex not or barely projecting dorsally above metapleural gland when observed in profile view (Fig. 6F–L). Abdominal tergum IV with anterolateral corners directed anterodorsally from helcium and forming lateral margins of concavity which receives entire posterior region of petiole when gaster raised (Fig. 7A); petiole largely obscured by abdominal tergum IV in dorsal view. Facial region between antennal toruli flat, effaced frontal carinae not raised above toruli (Fig. 7C) … ***Prenolepis* genus group** (see LaPolla *et al*. 2012, LaPolla & Fisher 2014)

- Petiole without posterior elongation. Node squamiform, not reduced, apex projecting dorsally above metapleural gland when observed in profile view (Fig. 6B–E). Abdominal tergum IV with anterolateral corners weakly or not raised anterodorsally from helcium, corners not forming lateral margins of concavity, and not concealing entire posterior region of petiole when gaster raised (Fig. 7B); petiole visible in dorsal view, not obscured by abdominal tergum IV. Facial region between antennal toruli bulging to concave, frontal carinae raised above toruli (Fig. 7D) … **3**

**3.** Eyes situated in anterior half of head as measured in full-face view (Fig. 10A). Metapleural gland orifice reduced, with opening directed posteriorly. Propodeal spiracle situated in lower half of propodeum in profile view. Antennomere III broader than long. Hypostoma lacking carina along lateral margins … ***XXX* gen. n.** (compare also †*XXX* **gen. nov.**, below)

**Fig. 10.**
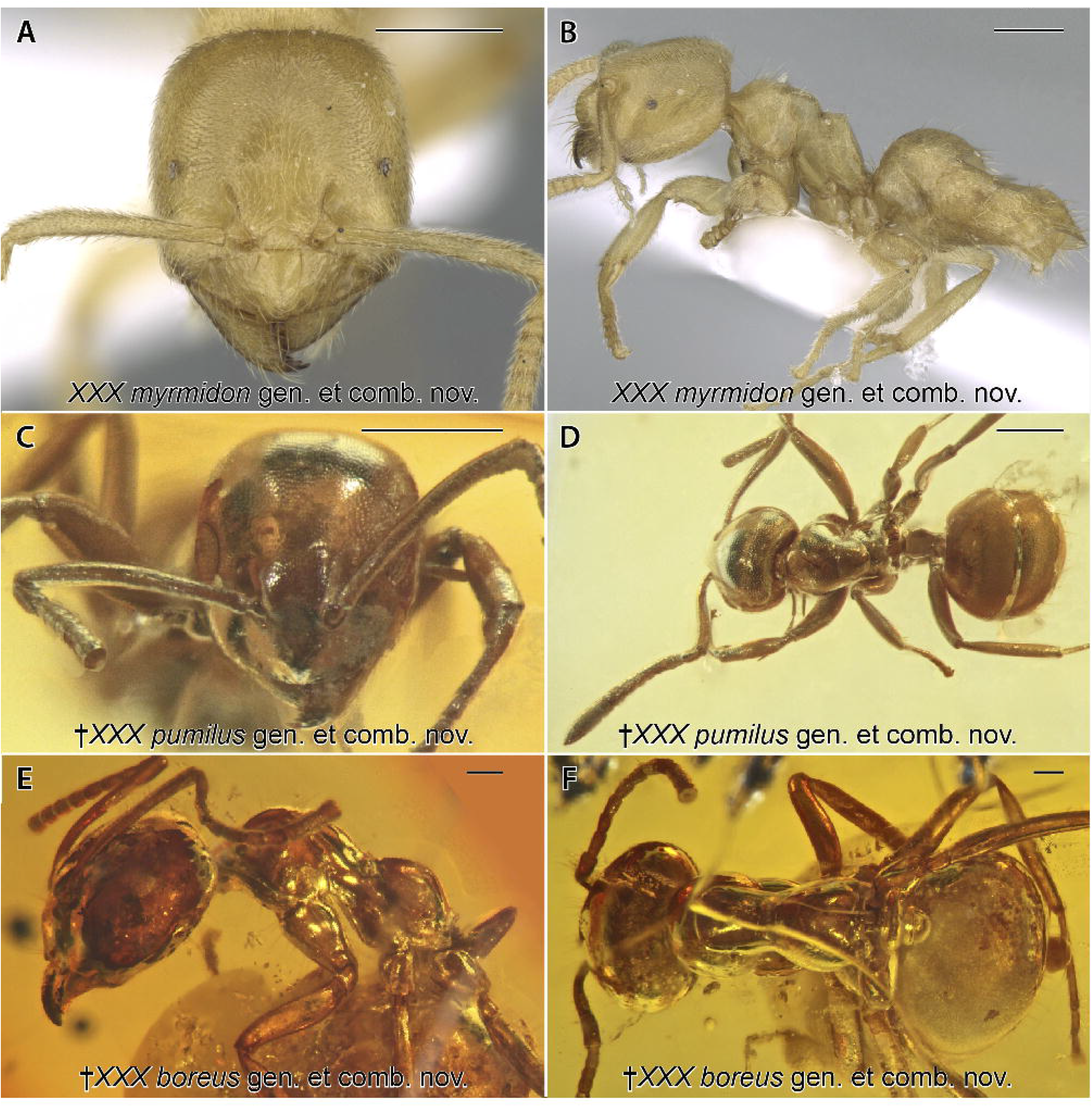
Habitus images for the three new genera recognized in this study; scale bars = 0.25 mm. A, B, *XXX myrmidon* **gen.** and **comb. n.** (CASENT0906115); C, D, †*XXX pumilus* **gen.** and **comb. n.** (SMFBE1226); E, F, †*XXX* **gen.** and **comb. n.** (GZG-BST04646). Views: A = full-face; B, E = profile; C = laterally oblique facial; D, F = dorsal. Observe: (1) The compound eyes of *XXX* and †*XXX* are mid-set, and (2) †*XXX* **gen. n.** has a distinctly-carinate propodeal foramen, vertical petiolar node, and non-grooved (lateromedially convex) third abdominal tergite.

- Eyes situated in posterior half of head as measured in full-face view (Fig. 9). Metapleural gland orifice small to very large, opening laterally as well as posteriorly. Propodeal spiracle situated at or above midheight of propodeum in profile view. Antennomere III usually longer than broad. Hypostoma with carina along lateral margins … **4 (*Lasius* genus group)**

**4.** Maxillary palpomeres III and IV strongly flattened and greatly elongated, length of each exceeds length of apical antennomere. Ventral surface of head with psammophore, *i.e.*, margined by long setae (Fig. 6D). Mesothorax and metathorax elongated, mesonotum meeting metanotum at very low angle (Fig. 6D). Compound eyes set at or near extreme posterior margin of head in full-face view; eyes usually separated by < 1 eye diameter … ***Myrmecocystus*** (see Snelling 1976, 1982)

- Maxillary palpomeres III and IV circular to elliptical in cross-section and not elongated, length of each is less than length of apical antennomere. Ventral surface of head without psammophore (Fig. 6B, C). Mesothorax not elongated and metathorax very rarely elongated, mesonotum usually meeting metanotum at steep angle (Fig. 6B, C). Compound eyes set further away from posterior head margin in full-face view; eyes usually separated by > 1 eye diameter … **5 (*Lasius*)**

**5.** Metapleural gland enlarged internally, atrium (internal chamber of gland) bulging (forming bulla) and conspicuously visible from external view in profile (Fig. 7E)*. Usually, distance between atrium and propodeal spiracle < 2 spiracular diameters as viewed with spiracle and gland in same plane of focus. Compound eyes usually somewhat- to very-reduced; **if** compound eye large, **then** offset basal tooth present on mandible **and** body yellowish brown … ***Lasius flavus* clade**

- Metapleural gland small, atrium flat to concave, inconspicuous from external view in profile (Fig. 7F). Usually, distance between atrium and propodeal spiracle > 2 spiracular diameters as viewed with spiracle and gland in same plane of focus. Compound eyes relatively large **and** offset basal tooth absent; **if** offset basal tooth present, **then** body jet black … ***Lasius niger* clade**

*Note: The enlarged state of the metapleural gland bulla is a defining synapomorphy of the *Lasius flavus* clade. A slightly enlarged bulla is observable in the *fuliginosus* species group, but these jet-colored ants are otherwise highly distinct.

#### Genus group of *Lasius*

(Figs. 6B, C, D, 7B–F, 8A–K, 9A–J)

**Included genera:** *Lasius*, *Myrmecocystus*.

**Definition (worker):**

1. With characters of Lasiini.
2. Mandible with 4–11 teeth.
3. Palp formula 6,4 or 3,4.
4. Basal and masticatory mandibular margins meeting at a weakly oblique angle.
5. Frontal carinae conspicuous, > 0.5 × anteroposterior antennal torulus diameter.
6. Frontal region of head, between antennal toruli bulging, with frontal carinae raised above toruli (note 1).
7. Antenna 12-merous.
8. Third antennomere usually longer than broad (note 2).
9. Compound eye set in posterior half of head (note 3).
10. Ocelli usually present (note 4).
11. Dorsum of head with at least some standing setae in addition to those on posterolateral head corners.
12. Mesonotum, metanotum, and/or propodeum discontinuous, metanotum differentiated.
13. Anterolateral margin of mesopleural area, near posteroventral pronotal margin, with longitudinally oriented bosses subtending transverse groove or not (note 5).
14. Metapleural gland small to very large, orifice opening laterally to posterolaterally, not directed posteriorly.
15. Propodeal spiracle situated in lower 2/3 of propodeum.
16. Petiole with raised squamiform node, without posterior elongation.
17. Abdominal tergum III vertical, without deep groove for receiving petiole; not concealing petiole when gaster raised.
18. Tergosternal suture of abdominal segment III not raised dorsally before unfusing near spiracle, rather anterior margin of abdominal sternum III directed anteriorly away from helcium before narrowly curving posteriorly.
19. Pubescence of abdominal terga VI and VII absent, or present and linear to weakly curved.

#### Notes on definition

*Note 1*. The polarity of this character is uncertain. Both the *Lasius* and *XXX* genus groups have raised frontal regions, but neither *Cladomyrma* nor the *Prenolepis* genus group do.

*Note 2*. Antennomere III is longer than broad in most *Lasius*, except for the *flavus* species group (former *Cautolasius*).

*Note 3*. Although we suspect that posteriorly set eyes are a plesiomorphy for the crown Formicidae, our ancestral state estimation results indicate that the posteriorly set eyes of the *Lasius* genus group are derived, and that they are homoplastic with respect to *Prenolepis* + *Zatania* and *Paratrechina* (Fig. S10).

*Note 4*. Ocelli are not expressed in various species of the *flavus* clade. Generally, ocellus presence is a highly variable state, even within species, thus is a weak defining feature for worker ants.

*Note 5*. The development of the boss subtending the transverse mesosternal groove appears to be an apomorphy of the *Lasius* genus group or *Lasius* itself, with loss in *Myrmecocystus* and reduction in the *L. fuliginosus* group. The groove and shoulder (boss) are present in a reduced form in *XXX* and are variable in the *Prenolepis* genus group. Specifically, they are reduced such that no rim is present on the anterior mesosternal margin (*e.g.*, *Nylanderia*), or the groove is elongated (*e.g.*, *Zatania*, *Prenolepis*, *Paratrechina*); some *Paraparatrechina* have either the groove (narrow) or the shoulder.

#### Genus *Lasius* Fabricius, 1804

= *Donisthorpea* Morice & Durrant, 1915

= *Acanthomyops* Mayr, 1862 **syn. rev.**

= *Austrolasius* Faber, 1967 **syn. n.**

= *Cautolasius* Wilson, 1955 **syn. n.**

= *Chthonolasius* Ruzsky, 1912 **syn. n.**

= *Dendrolasius* Ruzsky, 1912 **syn. n.**

(Figs. 6B, C, 7E, F, 8A–K, 9A–J)

**Type species:** *Formica nigra* Linnaeus, 1758 (= *Lasius niger*)

#### Subgeneric classification remarks

The modern body of taxonomic work on *Lasius* was initiated by Wilson’s revision of the genus (Wilson 1955), which was classified into four subgenera at the time: *Cautolasius*, *Chthonolasius*, *Dendrolasius*, and *Lasius s. str.* In this work, Wilson provided a phylogeny of *Lasius* (Fig. 1A), but this treatment was an intuitive account of what he considered trends in the evolution of the morphology and biogeography of the genus. Subsequently, the subgenus *Austrolasius* was erected for a few socially parasitic species (Faber 1967) and the former genus *Acanthomyops* was included in *Lasius* as the sixth subgenus (Ward 2005).

The first inference based on molecular data was presented by Hasegawa (1998), who used COI to investigate relationships of four *Lasius* species (phylogeny not figured here). Since then, two major attempts at resolving the phylogeny were presented by Janda *et al*. (2004) and Maruyama *et al*. (2008) (Figs. 1B, C). Both studies used a combination of morphological characters and molecular data, including mitochondrial (COI, COII, tRNA-Leu) and ribosomal (16S) markers. A more recent effort focused on the phylogeny of European species related to *Lasius niger* (*Lasius s*. *str*.) and included nuclear genes *LW Rh* and *wg* in addition to 16S and COI (Talavera *et al*. 2015).

To date, the monophyly of the subgenera has not been questioned except for the nominotypical subgenus (Janda *et al*. 2004, Maruyama *et al*. 2008). Recently, Seifert (2020, p. 21) stated that there is clear justification for elevating the subgenera to generic status. Such an action would considerably complicate the classification of the *Lasius* genus group because of the robustly supported paraphyly of the subgenera that we have uncovered here (Figs. 1D, 2, S1, Table 4). Specifically, in order to retain monophyly at the genus rank, four new genera would need to be erected for the species groups of *brunneus*, *nearcticus*, *atopus*, and *pallitarsis*. An additional issue would be the placement of the species which have not been sequenced, particularly those of the *niger* species group and, for example, the recently described *L. brevipalpus* which is *incertae sedis* in the *niger* clade. The strongest morphological reorganization at the generic or subgeneric levels would be to recognize the reciprocally monophyletic *niger* and *flavus* clades as *Lasius* and *Acanthomyops*, respectively, but we refrain from doing so here (for our rationale, see “Species group classification of *Lasius*” below). **Comments on extant species:**

We determine that one species, *Lasius escamole* Reza, 1925, should be excluded from *Lasius* and considered a junior synonym of the dolichoderine *Liometopum apiculatum* Mayr, 1870, **syn. n.** Reza (1925) described *Lasius escamole* in the context of a cultural study on the eponymous traditional Mexican dish known to be made from the larvae of *L. apiculatum* (Hoey-Chamberlain *et al*. 2013). Although Reza’s description and illustrations are extremely vague, it is possible to see details that point to a dolichoderine identity. In the figures of the original description, the mandibles have long masticatory margin and small, sharp, even denticles, the ventral metasoma is shown as a slit-like anal opening rather than a formicine-like acidopore, and various figures display the fine, dense, appressed pilosity characteristic of *Liometopum*, but no erect setae as expected for *Lasius*.

Many species have been added to *Lasius* since Wilson’s revision, mostly in Europe and Mediterranean, while North American taxa have largely remained untreated except for a thorough revision of *Acanthomyops* (Wing 1968). Careful research has revealed multiple Palearctic *Lasius* species that show only subtle morphological differentiation from close relatives (Seifert 1983, 1990, 1991, 1992, 2020; Schlick-Steiner *et al*. 2003). There is no reason to believe that North America does not harbor a diverse fauna of such “cryptic species”. For example, the question of the putative Holarctically distributed *Lasius* species was resolved by Schär *et al*. (2018) who elevated to species the American representatives of *L. alienus*, *L. flavus*, and *L. umbratus*, recognizing the following revived taxa, in order: *L. americanus*, *L. brevicornis*, and *L. aphidicola*. Renewed focus on the Nearctic fauna is necessary, as is expanded sequencing at the global scale.

#### Comments on extinct species

Without having scored the Baltic *Lasius* fossils other than †*L. schiefferdeckeri*, *i.e.*, †*L. punctulatus* and †*L. nemorivagus*, we are unable to quantitatively address their placement. Historically, †*L. punctulatus* and †*L. schiefferdeckeri* were considered to be members of *Lasius s. str.* (Wilson 1955, Dlussky 2011), while the queen-based †*L. nemorivagus* was placed in *Chthonolasius* (Wilson 1955) later to be implicitly considered *incertae sedis* in the genus (Dlussky 2011).

Our combined-evidence dating analyses recover †*Lasius schiefferdeckeri* as sister to or within the *Lasius* genus group (Figs. 4, S7–S9). As the specific relationship of the fossil to the extant species of the *Lasius* genus group is uncertain, we conservatively consider the fossil *incertae sedis* in *Lasius*. There remains the possibility that †*L. schiefferdeckeri* is ancestral to the extant *niger* clade and is indicative of low rates of phenotypic transformation, as suggested by Mayr (1868), Wheeler (1915), and Wilson (1955). The placement of †*L. schiefferdeckeri* may be refined in future study by scoring characters which are explicitly derived from comparison of the *brunneus* and *niger* groups within the *niger* clade.

The two differentiating traits proposed for the *brunneus* and *niger* subclades are, on average, < 8 mandibular teeth in the *brunneus* subclade (Seifert 1992; some *niger* subclade species with < 8), and presence of a subapical cleft in the mandibles of *brunneus* subclade males (Wilson 1955). While †*L. schiefferdeckeri* demonstrates both a tooth count of < 8, and presence of a subapical cleft, the latter character is probably plesiomorphic of the *Lasius* genus group, and the polarity of the former is uncertain. Notably, Wilson (1955) observed that the male mandibles of †*L. schiefferdeckeri* are observed to vary from the “*brunneus* form” to the derived “*niger* form”. With these three traits in mind, it does seem reasonable that †*L. schiefferdeckeri* is stem to or directly ancestral to the *niger* clade.

#### Note on biology

Despite the interest in this genus, however, basic natural history remains unknown for many species, including the morphologically aberrant *Lasius atopus* and the species we sequence here, “*L.* nr. *atopus*”. Only a handful of recently published studies have addressed the behavior of some of the more rarely encountered species (*e.g*., Raczkowski & Luque 2011), indicating that more effort is needed to elucidate the biology of *Lasius*.

### Species group classification of *Lasius*

To avoid following the undesirable trend where taxonomy is divorced from molecular phylogenetics (Steiner *et al*. 2009, Ward 2011), we propose an informal species-group classification of *Lasius* which reflects the phylogenetic structure within the genus. As the ICZN rules do not apply to informal species groups, this arrangement is also more adaptable to future refinements and new phylogenetic findings, thus contributing to taxonomic stability. Two main clades are recognized, those of *L. niger* (70 spp. + 1 ssp.) and *L. flavus* (54 spp.), with three and seven species groups, respectively. Additionally, all 21 valid extinct species are formally considered *incertae sedis* in the genus, while 14 extant species are considered *incertae sedis* in the genus and unidentifiable.

Should the resurrection of subgenera be deemed advisable and desirable, we recommend dividing the genus into *Lasius s. str.* and *Acanthomyops* along the boundaries of the *niger*- and *flavus*-clade split. Resurrection of *Acanthomyops* may be justified, as it is readily recognizable by the autapomorphic condition of its metapleural gland (see couplet 5 of key to lasiine genera above, Fig. 7E & F, also Appendix S1), and the relatively old split between the *flavus* and *niger* clades (Figs. 4 & 5). We have elected not to do so here because: (1) our molecular sampling is still fractional at the species level (9/70 *niger* clade, 12/54 *flavus* clade); (2) transfer of *Lasius* subgenera to *Acanthomyops* may be disruptive to the practicing systematic community; (3) Sanger data are being rapidly superseded by UCE data for ant systematics; and (4) morphological investigation is necessary to diagnose the newly recovered subclades. In either case, the exceptional morphometric data being generated for *Lasius* (*e.g.*, Seifert 1982, 1992, 2020; Seifert & Galkowski 2016) will be valuable for model-based phylogenetic analysis combining phenotypic and genotypic data (*e.g.*, Barden *et al*. 2017a, Prebus 2017), particularly in the light of the greater information content of continuous trait data relative to discretized characters (Parins-Fukuchi 2018). Ultimately, our goal is to encourage the consilience of morphological and molecular data, and we view both as critical for a complete understanding of evolution and systematics.

The following inter-group transfers are made: (1) Members of non-monophyletic former *Lasius s. str.* are dispersed among *Lasius brunneus*, *L. niger*, and *L. pallitarsis* groups; (2) *Lasius atopus* is removed from the *L. umbratus* group (former *Chthonolasius*); and (3) *L. nearcticus*, and *L. sonobei* are removed from the *L. flavus* group (former *Cautolasius*). For diagnosis of the *niger* and *flavus* clades, see the key to extant genera of Lasiini above. The task of morphologically distinguishing two pairs of clades, those of *brunneus* and *niger* and of *flavus* and *nearcticus*, is difficult. Future phylogenetic studies are expected to refine the boundaries of the of these species-groups. Note that, in the list below, [e] indicates that specimen(s) of the taxon have been examined while empty brackets, [], indicate that specimens were unavailable and placed based on study of the literature. Species that were included in our molecular sampling are **bolded**. Genus group type species are marked with asterisks (*). Overall, with the exclusion of *L. myrmidon* and *L. escamole*, 118 of the 125 valid, extant species (124) and subspecies (1) were examined either directly or indirectly (via AntWeb). For authorities and taxonomic synopses, refer to Bolton (1995) or AntCat (Bolton 2021). Finally, it should be noted that some groups contain additional, undescribed species.

#### I. Clade of *niger*

##### 1. Species group of *brunneus* (*L. s. str.* part 1/3)

Constituent species (11): *austriacus* [e], ***brunneus*** [e], *excavatus* [e], *himalayanus* [e], *israelicus* [e], *lasioides* [e], *neglectus* [e], *precursor* [e], *silvaticus* [e], *tapinomoides* [e], ***turcicus*** [e].

*Distribution*: Palearctic.

*Note*: Within the *brunneus* species group as circumscribed here, two complexes are recognized by Seifert (2020): that of *brunneus* comprises *brunneus*, *excavatus*, *lasioides*, *himalayanus*, and *silvaticus* (5 spp. total); that of *turcicus* comprises *austriacus*, *israelicus*, *neglectus*, *precursor*, *tapinomoides*, and *turcicus* (6 spp. total).

##### 2. Species group of *fuliginosus* (= *Dendrolasius*)

Constituent species (8): *buccatus* []*, capitatus* [e]*, fuji* []*, **fuliginosus\**** [e]*, morisitai* []*, nipponensis* [e]*, orientalis* [e]*, **spathepus*** [e].

*Distribution*: Palearctic.

##### 3. Species group of *niger* (*F. s. str.* part 2/3)

Constituent species (50), subspecies (1): *alienus* [e]*, **americanus*** [e], *balearicus* [], *bombycina* [e], *casevitzi* [e], *chinensis* [e], *cinereus* [e], *coloratus* [e], *creticus* [e], *cyperus* [e], *crypticus* [e], ***emarginatus*** [e], *flavescens* [e], *flavoniger* [e], *grandis* [e], *hayashi* [e], *hikosanus* [], *hirsutus* [], *illyricus* [e], *japonicus* [e], *kabaki* [e], *karpinisi* [e], *koreanus* [e], *kritikos* [e], *lawarai* [e], *longipalpus* [e], *magnus* [e]*, maltaeus* [e], *mauretanicus* [e], *neoniger* [e]*, **niger\**** [e]*, niger pinetorum* [], *nigrescens* [e], *obscuratus* [e], *paralienus* [e], *persicus* [e], *piliferus* [e], ***platythorax*** [e], *productus* [e]*, **psammophilus*** [e], *sakagamii* [e], *schaeferi* [e]*, schulzi* [e], *sichuense* [e], *sitiens* [e]*, tebessae* [e], *tunisius* [e], *uzbeki* [e], *vostochni* [e], *wittmeri* [e], *xerophilus* [e].

*Distribution*: Holarctic.

*Note 1*: Our molecular sample for *L. americanus* represents an undescribed species.

*Note 2*: Within the *niger* species group as circumscribed here, Seifert (2020) recognizes two morphometrically diagnosable complexes: that of *obscuratus* comprises *criticus*, *obscuratus*, *piliferus*, and *psammophilus*; that of *paralienus* comprises *bombycina*, *casevitzi*, *kritikos*, and *paralienus*.

##### 4. Species *incertae sedis* in the clade of *niger*

Species (1): *brevipalpus* [e].

*Note*: The species *brevipalpus* was described by Seifert (2020) who remarked that its relationship to other “*sensu stricto*” *Lasius* is uncertain due to a unique character combination, at least within the Asian fauna. We confirm placement of this species in the *niger* clade.

#### II. Clade of *flavus*

##### 1. Species group of *atopus* (= *Chthonolasius* part 1/2)

Constituent species (1): ***atopus*** [e].

*Distribution*: Restricted to western North America (California).

*Notes*: This species was classified in the former subgenus *Chthonolasius*; our sequenced specimen represents a new species near *atopus*.

##### 2. Species group of *carniolicus* (= *Austrolasius*)

Constituent species (2): ***carniolicus\**** [e]*, reginae* [e]. *Distribution*: Palearctic.

##### 3. Species group of *claviger* (= *Acanthomyops*)

Constituent species (16): *arizonicus* [e]*, bureni* [e]*, **californicus*** [e]*, claviger\** [e]*, colei* [e]*, coloradensis* [e]*, creightoni* [e]*, interjectus* [e]*, **latipes*** [e]*, mexicanus* [e]*, murphyi* [e]*, occidentalis* []*, plumopilosus* [e]*, pogonogynus* [e], *pubescens* [e]*, subglaber* [e].

*Distribution*: Nearctic.

##### 4. Species group of *flavus* (= *Cautolasius* part 1/2)

Constituent species (7): *alienoflavus* [e]*, **brevicornis*** [e], *elevatus* [e]*, fallax* [e]*, **flavus\**** [e]*, **myops*** [e]*, talpa* [e].

*Distribution*: Holarctic.

*Notes*: This group corresponds to the former subgenus *Cautolasius*, in part. The inclusion of *alienoflavus*, *elevatus*, *fallax*, and *talpa* is provisional; molecular data ought to be generated for these species, and a critical reanalysis of phenotypic traits conducted.

##### 5. Species group of *nearcticus* (= *Cautolasius* part 2/2)

Constituent species (2): ***nearcticus*** [e], ***sonobei*** [e].

*Distribution*: Holarctic.

##### 6. Species group of *pallitarsis* (= *Lasius s. str.* part 3/3)

Constituent species (1): ***pallitarsis*** [e].

*Distribution*: Nearctic.

*Note*: This species was classified in the former subgenus *Lasius s. str.* and has previously been recovered elsewhere in the phylogeny by Janda *et al*. (2004) and Maruyama *et al*. (2008).

##### 7. Species group of *umbratus* (= *Chthonolasius* part 2/2)

Constituent species (25): *aphidicola* [e], *balcanicus* [e]*, bicornis* [e]*, citrinus* [e]*, crinitus* [e]*, distinguendus* [e]*, draco* []*, humilis* [e]*, jensi* [e]*, longiceps* []*, meridionalis* [e]*, mikir* [e]*, minutus* [e]*, mixtus* [e]*, nevadensis* [e]*, nitidigaster* [e]*, przewalskii* []*, rabaudi* [e]*, **sabularum*** [e]*, speculiventris* [e]*, **subumbratus*** [e]*, tibialis* [], *umbratus\** [e]*, vestitus* [e]*, viehmeyeri* [e].

*Distribution*: Holarctic.

*Note*: This group corresponds to the old genus *Chthonolasius,* with the exclusion of *L. atopus*.

One of our molecular terminals of this group is an undescribed species near *subumbratus*.

#### III. Incertae sedis in genus

##### 1. Impression fossils

Species (18), subspecies (1): *<u>West Palearctic</u>*: (28.4–23.0 mya, <u>Aix-en-Provence</u>): †*epicentrus* [e]; (12.7–11.6 mya, <u>Radoboj</u>): †*anthracinus* [e], †*globularis* [],†*longaevus* [e], †*longipennis* [e], †*occultatus* [e], †*occultatus parschlugianus* [], †*ophthalmicus* [e]; (12.7–11.6 mya, <u>Berezovsky</u>): †*tertiarius* []; (11.6–5.3 mya, <u>Schnossnitz</u>): †*oblongus* []; (8.7–7.2 mya, <u>Joursac</u>): †*crispus* []; (8.7–2.6 mya, Lake <u>Chambon</u>): †*chambonensis* [e]; *<u>East Palearctic</u>*: (20.4–16.0 mya, <u>Shanwang</u>): †*inflatus* [], †*mordicus* [],†*truncatus* [], †*validus* []; (16.0–11.6 mya, <u>Vishnevaya</u>): †*vetulus* []; *<u>Nearctic</u>*: (48.6–40.4 mya, <u>Kishenehn</u>): †*glom* [e]; (37.2–33.9 mya, <u>Florissant</u>): †*peritulus* [e].

*Note*: All impression fossils attributed to *Lasius* are here considered to be *incertae sedis* within the genus. The detail preserved for all examined specimens is insufficient to challenge generic placement but does match the gross *gestalt* of *Lasius*. Critical study of these fossils, especially †*L. glom* and †*L. peritulus*, should be undertaken.

##### 2. Amber fossils

Species (3): †*nemorivagus* [], †*punctulatus* [e], †*schiefferdeckeri* [e].

*Deposits*: *Palearctic*: Baltic, Bitterfield, Danish-Scandinavian, and Rovno (Ukrainian).

*Note*: These species, from coniferous ambers (Wolfe *et al*. 2016), are considered stem *Lasius* as per the “Comments on extinct species” section above.

#### IV. Extant species

Species (14): *alienoniger* Forel, 1874, *emeryi* Ruzsky, 1905, *exulans* Fabricius, 1804, *longicirrus* Change & He, 2002, *minimus* (Kuznetsov-Ugamsky, 1928), *monticola* (Buckley, 1866), *mixtoumbratus* Forel, 1874, *monticola* Buckley, 1866, *nigerrimus* (Christ, 1791), *nitidus* (Kuznetsov-Ugamsky, 1927), *pannonica* Röszler, 1942, *rubiginosa* Latreille, 1802, *ruficornis* Fabricius, 1804, *transylvanicus* Röszler, 1932.

*Note*: The species *exulans*, *monticola*, *rubiginosa*, and *ruficornis* were rendered *incertae sedis* by Bolton (1995); we are doubtful that these names will be resolved. Recently, Seifert (2020) considered *alienoniger*, *emeryi*, *longicirrus*, *minimus*, *nigerrimus*, *nitidus*, *pannonica*, and *transylvanicus* to be *nomina nuda*. The species *nigerrimus* is unidentifiable to subfamily (Seifert 2020).

**Genus group of *XXX***

(Figs. 6E, 10A–D)

**Genera included: †*XXX* gen. n., *XXX* gen. n**.

**Definition (worker):**

1. With characters of Lasiini.
2. Mandible with 6–8 teeth.
3. Palp formula 6,4.
4. Basal and masticatory mandibular margins meeting at a weakly oblique angle.
5. Clypeus modified for reception of labrum (specifically, clypeus with anterolateral notches; note 1).
6. Frontal carinae conspicuous, > 0.5 × anteroposterior antennal torulus diameter.
7. Frontal region of head, between antennal toruli strongly bulging, frontal carinae raised above toruli (note 2).
8. Antenna 12-merous.
9. *Third antennomere broader than long* (note 3).
10. Compound eyes set at about head midlength.
11. Ocelli absent.
12. *Dorsum of head completely without standing setae or with only a few small setae around each posterolateral head corner* (note 4).
13. Mesonotum, metanotum, and/or propodeum discontinuous, metanotum undifferentiated medially.
14. Mesopleural anterodorsal margin, near posterolateral region of pronotum, inconspicuously bulging, this weak shoulder a very narrow groove which traverses the mesosternum.
15. Metapleural gland very small, orifice directed almost completely posteriorly (note 5).
16. Propodeal spiracle situated in lower 2/3 of propodeum.
17. Legs entirely or almost entirely devoid of standing setae.
18. Petiole with raised squamiform node, without posterior elongation.
19. Abdominal tergum III vertical, without deep groove for receiving petiole; not concealing petiole when gaster raised (note 6).
20. Tergosternal suture of abdominal segment III not raised dorsally before unfusing near spiracle, rather anterior margin of abdominal sternum III directed anteriorly away from helcium before narrowly curving posteriorly (note 6).
21. Pubescence of abdominal terga VI and VII linear.

##### Notes on definition

*Note 1*. Modification of the anterior clypeal margin for reception of the labrum is here interpreted as a synapomorphy of *XXX* plus the core *Prenolepis* genus group. The modification is most easily observed in anterior or anterodorsal view. In most Formicinae, the perceived anterior clypeal margin with head in full-face view is usually a carina which runs across the clypeus from the lateral clypeal margins. This carina may be raised or otherwise modified; regardless, the carina itself or the anterior region of the clypeus is produced anteriorly, concealing the clypeolabral articulation. In the *Prenolepis* and *XXX* genus groups, the clypeolabral articulation is usually exposed or nearly exposed laterally by notches in the anterior clypeal carina, although it may be exposed medially where the anterior clypeal carina is absent.

*Note 2*. See note 1 for the *Lasius* genus group.

*Note 3*. Surprisingly, this distinguishes members of the *XXX* genus group from most formicine genera, with the exception of some *Myrmelachista*, various plagiolepidines, and *Lasius* (*Cautolasius*). *Cladomyrma* species have a cone-shaped third antennomere which is usually longer than the diameter of its base (and often apex), except in *C. dianeae* for which length and basal diameter are subequal. Some *Pseudolasius* and *Paraparatrechina* approach having subequal diameter and length.

*Note 4*. Mei (1998), in his diagnosis of *XXX myrmidon*, indicated that these standing setae were absent except for one specimen.

*Note 5*. This state may be a synapomorphy for the clade consisting of *XXX* plus the *Prenolepis* genus group. It was not possible to evaluate this with confidence for †*XXX*, but from the dorsal view of specimen SMFBE1226 on AntWeb, the gland does appear to be small.

*Note 6*. These characters could not be evaluated with confidence for †*XXX* with the available material or descriptions.

**Comments:** Although we could not evaluate all of these characters for †*XXX*, the genus is consistently recovered as sister to *XXX* in combined analysis (Figs. 4, S7–S9). The *XXX* genus group is most easily differentiated from either the *Lasius* or *Prenolepis* genus groups by the combination of mid-set eyes, the posteriorly directed and reduced metapleural gland orifice, raised frontal carinae, and almost complete lack of standing setae on the head. Additionally, both genera have broad third antennomeres and unusually small bodies relative to *Lasius*, being < 2 mm in total length (Mei 1998, Dlussky 2011).

Genus *XXX* GEN. NOV.

(Figs. 6E, 10A, B)

**Type and included species:** *Lasius myrmidon* Mei, 1998: 177, figs. 1–11 (w.). Original designation.

**ZooBank LSID:** http://zoobank.org/urn:lsid:zoobank.org:act:AC5A1489-C805-4035-A1C0-3DDBC6E1206D

**Definition (worker):**

1. With characters of *XXX* genus group.
2. *Dorsal mandibular groove absent* (note 1).
3. Ventromedial base of mandible without trough or impression (note 2).
4. Maxillary palps short, exceeding hypostomal margin but not reaching postgenal bridge midlength.
5. Maxillary palpomere 3 longest, 4 shorter but as long as 5 and 6 together (note 3).
6. Clypeus, in lateral view, convex and weakly bulging anteriorly (note 3).
7. Anterior tentorial pit situated lateral to midlength of the epistomal suture.
8. *Lateral hypostomal carina absent* (note 4).
9. *Compound eyes absent, reduced, or vestigial, with at most 9 ommatidia* (note 5).
10. *Propodeal spiracle situated distinctly in lower half of propodeum* (note 6).
11. Legs almost entirely devoid of standing setae (note 3).
12. Petiolar node weakly inclined anteriorly, node well-developed, squamiform (note 7).

##### Notes on definition

*Note 1*. The dorsal mandibular groove is discernable in all examined *Lasius* and *Prenolepis* genus group taxa.

*Note 2*. Presence of a trough on the ventromedial base of the mandible is a newly detected synapomorphy of the core *Prenolepis* genus group. The impression is enhanced when the ventromedial mandibular margin is carinate and/or produced medially, and best seen when the mandibles are open and with the head in lateral anteroventral view. The trough may be reduced or absent in some species.

*Note 3*. Previously included in the original diagnosis of the species *XXX myrmidon* by Mei (1998).

*Note 4*. The lateral hypostoma is usually delimited by a carina which is discontinuous with the medial hypostomal lamina. Among formicines, the lateral hypostomal carina is absent only in *Acropyga* (some species) and *Brachymyrmex*, both genera outside of Lasiini.

*Note 5*. The specimens which were available for examination had 5–8 ommatidia; the maximum ommatidium count is from Mei (1998, p. 178).

*Note 6*. A lowered propodeal spiracle appears sporadically replicated in only a few *Prenolepis* genus group members.

*Note 7*. The petiolar node is strongly inclined anteriorly in the core *Prenolepis* genus group. **Etymology:** A combination of the Greek “meta-” (with, across, after) and “lásios” (hairy), in reference to the former placement of the type species. Homophonously forming “metal-Asius”, invoking the death of the Trojan leader Asius during the assault on the Archaean wall of Troy. Masculine.

**Comments:** *XXX* is uniquely identified among the Lasiini by absence of the dorsal mandibular groove and lateral hypostomal carina, short and broad third antennomere, mid-set compound eyes (when present) which are reduced to at most 9 ommatidia, and near complete absence of standing setae on the head. High magnification may be required to evaluate the lateral hypostoma.

**Genus †*XXX* GEN. NOV.**

(Figs. 10C, D)

**Type and included species:** †*L. pumilus* Mayr, 1868. Original designation.

**ZooBank LSID:** http://zoobank.org/urn:lsid:zoobank.org:act:3E019A26-D68E-41DE-86C9-A26EA22D20CE

**Definition (worker):**

1. With characters of *XXX* genus group (note 1).
2. Maxillary palps long, reaching occipital foramen (note 2).
3. Compound eyes well-developed, with > 20 ommatidia.
4. Mesosomal dorsum devoid of setae.
5. Legs entirely devoid of standing setae.
6. Petiolar node weakly inclined anteriorly, node squamiform.

##### Notes on definition

*Note 1*. Several characters could not be evaluated for †*XXX*, including the ventromedial mandibular groove, palpomere proportions, clypeal profile, and lateral hypostomal carina.

*Note 2*. From Dlussky (2011).

##### Etymology

Combination of the Greek words “ēlektro-” (amber) and “lásios” (hairy), indicating the geological provenance and prior taxonomic placement of these lasiine ants. Masculine.

##### Comments

We place †*XXX* with *XXX* together based on the results of our phylogenetic analyses, and we interpret presence of the broad third antennomere and highly reduced cranial setation as synapomorphies of this clade. Although we have not examined the neotype of †*XXX pumilus*, designated by Dlussky (2011) and deposited in Muzeum Ziemi Polskiej Akademii Nauk in Warsaw, the specimen we have studied is unlikely to be misidentified as it exhibits unique diagnostic traits of the species among the Lasiini, let alone of the Baltic amber fauna, including absence of setae on the head dorsum, well-developed eyes, short and broad third antennomere, and very small body size (< 2 mm). Dlussky (2011) describes the eyes of both †*Lasius schiefferdeckeri* and †*E. pumilus* as “shifted somewhat posteriorly so that the length of [the] gena [is] more than [that of the] maximum diameter of [the] eyes”. This very general statement is true of both species, however the eyes of †*L. schiefferdeckeri* are distinctly set in the posterior head half whereas those of †*E. pumilus* are situated at head midlength, distinguishing it from all *Lasius* genus group members. †*XXX pumilus* differs from *M. myrmidon* by the following: (1) compound eyes large; (2) maxillary palps long, reaching occipital foramen; and (3) standing setae completely absent from head dorsum and mesosoma.

**Genus group of *Prenolepis***

(Figs. 6F–L)

**Genera included:** Euprenolepis, *Nylanderia*, *Paraparatrechina*, *Paratrechina*, *Prenolepis*, *Zatania*.

Definition (worker):

1. With characters of Lasiini.
2. Mandible with 4–7 teeth.
3. Palp formula variable: 6,4; 3,4; 5,3; 4,3; 3,3; 2,3; 2,2 (note 1).
4. *Basal and masticatory mandibular margins usually meeting at a strongly oblique angle*.
5. Clypeus modified for reception of labrum (specifically, clypeus with anterolateral notches; note 1 of *XXX* genus group).
6. Frontal carinae conspicuous, > 0.5 × anteroposterior antennal torulus diameter.
7. Frontal region of head between antennal toruli flat, frontal carinae not raised above toruli.
8. Antenna 11- or 12-merous.
9. Third antennomere usually longer than broad (see note 3 of *XXX* genus group).
10. Compound eyes usually at about head midlength, sometimes set posteriorly (see note 3 of *Lasius* genus group).
11. Ocelli present or absent.
12. Dorsum of head with standing setae.
13. Mesonotum, metanotum, and/or propodeum discontinuous, metanotum usually differentiated (note 2).
14. Mesopleuron anterodorsal margin, near posterolateral region of pronotum, usually without raised boss nor with narrow or broad groove traversing the mesosternum (note 4 of *Lasius* genus group).
15. Metapleural gland very small, orifice directed completely to nearly completely posteriorly (note 5 of *XXX* genus group).
16. Propodeal spiracle situated in lower 2/3 of propodeum.
17. Legs with or without standing setae.
18. *Petiole with very low node and posteriorly elongated*.
19. *Abdominal tergum III inclined anteriorly, with deep groove for reception of petiole, concealing most of petiole in dorsal view when gaster raised*.
20. *Abdominal tergum III with complete tergosternal fusion adjacent to helcium, and fusion raised anterodorsally, with sclerites becoming free high up on segment*.
21. *Pubescence of abdominal terga VI and VII usually dense, forming a uniform layer of short and strongly curved setae* (note 3).

##### Notes on definition

*Note 1*. The palp formula of most genera is 6,4; *Euprenolepis* has a palpomere count of 3,4 and in *Pseudolasius* it is variable but not 3,4 (Bolton 2003).

*Note 2*. The metanotum of many but not all *Paraparatrechina* is undifferentiated.

*Note 3*. These setae may or may not completely cover tergum VI. Absence of this characteristic pubescence occurs in and may be a synapomorphy of *Euprenolepis* and *Paratrechina*.

##### Comments

*Paratrechina* has recently expanded from a single species to five (LaPolla *et al*. 2010, 2013; LaPolla & Fisher 2014). One of these recently added species, *P. kohli* (Forel, 1916) was originally described in *Prenolepis*, transferred to *Nylanderia* (as a subgenus of *Paratrechina*), back to *Prenolepis*, and finally to *Paratrechina* (*sensu stricto*) (LaPolla & Fisher 2014). This species stood out from the remainder in its surface sculpturing, eye positioning, color, and mandibular dentition. It was observed during this study that *P. kohli* has a petiolar node which exceeds propodeal spiracle height, lacks the raised tergosternal fusion of abdominal segment III and posterior elongation of the petiole, and closely resembles species of *Anoplolepis*, specifically *A. carinata* (Emery, 1899) and *A. tenella* (Santschi, 1911). Additionally, *P. kohli* has 11-merous antennae, an undifferentiated metanotum, has posteriorly set eyes, and lacks ocelli, as is characteristic of *Anoplolepis* (Fisher & Bolton 2016). We find in combined analyses that *P. kohli* always falls outside of the Lasiini (Figs. 4, S7–S9). For these morphological and phylogenetic reasons, we transfer the species *Paratrechina kohli* to *Anoplolepis* in the Plagiolepidini, forming *A. kohli* (Forel, 1916) **comb. n.** (tribal transfer). Expanded sampling of male characters may improve placement of the fossil taxon †*Prenolepis henschei* with respect to the remainder of the genus group.

#### *Incertae sedis* in the Formicinae

##### Genus †Kyromyrma

Comparative morphological study of †*Kyromyrma* (∼92 mya, New Jersey amber; Grimaldi & Agosti 2000) at the gross (Figs. 8L, 9K) and fine scales reveals considerable morphological affinity to *Lasius* (holotype examined at AMNH). In the original description of †*Kyromyrma*, the authors did not address the problem of within-subfamily placement, merely noting that “the fossil bears an overall resemblance to *Prolasius*, mostly by virtue of the generalized morphology” (Grimaldi & Agosti 2000, p. 13681). Our combined evidence analyses resulted ambiguous support for the placement of †*Kyromyrma.* Results placed †*Kyromyrma* as sister to the Lasius genus group (Fig. S5), sister to the core Lasiini (Figs. S8 & S9), sister to all Lasiini (Fig. S7), or sister to Formicinae exclusive of Myrmelachistini (Fig. 4). Statistical support for these placements was uniformly low.

**Genus †*XXX* GEN. NOV.**

(Figs. 10E, F)

**Type and included species:**

†*Pseudolasius boreus* Wheeler, 1915. Original designation.

**ZooBank LSID:** http://zoobank.org/urn:lsid:zoobank.org:act:BFF515E7-9966-443F-8A34-555A4B41906B

**Definition (worker):**

1. With characters of the Formicinae (note 1).
2. *Cranium size variable, width somewhat broader than to about twice that of mesosoma* (note 2).
3. Mandible triangular, with 7–8 teeth.
4. Dorsal mandibular groove extending along lateral mandibular margin in dorsal view.
5. Palps reduced, not reaching midlength of postgenal bridge.
6. Frontal carinae raised above antennal toruli.
7. Antennal toruli abutting posterior clypeal margin.
8. Antennomere count unknown (note 3).
9. Antennomere III longer than broad.
10. Compound eyes situated posterior to head midlength.
11. Compound eyes not enormous and long axes subparallel.
12. Ocelli minutely present or absent.
13. Promesonotum domed, raised well above propodeum.
14. Metanotal groove well-developed.
15. Petiolar foramen in profile view clearly raised and margined with conspicuous thickened carina or lip.
16. Petiolar node squamiform, tall; dorsoventral height equal to or possibly exceeding dorsum of propodeum.
17. Abdominal segment III without raised tergosternal fusion.

##### Notes on definition

*Note 1*. Several characters were not possible to evaluate. These include the following: Reduction of third tooth from mandibular apex; palp formula; the metapleural gland; ventral length of the petiolar foramen; relative separation of the meso- and metacoxae; presence or absence of setal double-row on ventral tibial surfaces; petiolar apodeme conformation; cross-sectional shape of the petiolar sternum; and presence or absence of the post-helcial sulcus.

*Note 2*. The massive head of †*P. boreus* led Wheeler (1915) to erroneously assume that the specimens were majors of *Pseudolasius*; Wheeler did note the posteriorly set eyes of †*P*. *boreus*, one of the distinctions between it and *Pseudolasius*, but vacillated based on the insufficient knowledge of the latter genus at that time.

*Note 3*. The examined specimens are missing their terminal antennomeres, and no previous descriptions of †*P*. *boreus* include an antennomere count.

**Etymology:** A simplification of “pseudo-*Pseudolasius*” inspired by Phil Collins. Masculine.

**Comments:** †*XXX boreus* **comb. n.** was previously placed in the Lasiini (Emery 1925, Dlussky & Fedoseeva 1988, Bolton 1994), and has been considered to be a member of the genus *Pseudolasius* (Wheeler 1915, LaPolla & Dlussky 2010). Based on an examination of a syntype (GZG-BST04646) and a non-type (NHMW1984-31-211) worker, as well as the original description from Wheeler (1915), we reject both placements. Neither specimen can be mistaken due to the unique combination of characters they display, including setation, high petiolar nodes, massive crania, and mesosomal form. †*XXX boreus* does not display the modifications of the third abdominal segment characteristic of the *Prenolepis* genus group, also rejecting its original placement in *Pseudolasius* (see diagnostic note 2 above). †*XXX* is excluded from the Lasiini overall by two additional characters: (1) The petiolar node, which is completely vertical and very tall dorsoventrally, is as tall as or possibly taller than the propodeum; and (2) the raised and conspicuously lipped petiolar foramen. No lasiine, extant or extinct, has such a tall petiolar node, nor do any species display the petiolar foramen conformation observed in †*XXX*. We cannot confidently place †*XXX* in any extant tribe, therefore, we consider †*XXX incertae sedis* in the Formicinae. As with other fossil taxa which are of uncertain placement in the Formicinae, micro-CT scans could be used to evaluate the form of the helcium and third abdominal sternum.

## Conclusion

Revisionary works have traditionally scrutinized morphological systems for the systematic classification of a given group of interest. Here, we integrate traditional morphology with molecular phylogeny to understand the ant tribe Lasiini in the context of the subfamily Formicinae. Of particular interest to us are the suite of phenotypic characteristics which are used to define taxa. After reevaluating the fossil record of the Formicinae, we were able to estimate the geological age of phylogenetic branching events in the subfamily; with this chronogram, we formally estimated the probability that key diagnostic characters are ancestral or derived for any given clade. To this end, we have tested polarity hypotheses for traits used by Bolton in his crucial morphological reclassification of the Formicidae (Bolton 2003), as well as the form of the proventriculus—a structure used by taxonomists for well over a century to classify the Formicinae and other subfamilies (*e.g.*, Forel 1874, 1878; Emery 1888, 1925). Additionally, we find that two life history traits, temporary social parasitism and fungiculture, have both evolved twice in the genus.

One key question arising from our work is how to treat the two genera of the *Lasius* genus group relative to the fossils of *Lasius*, given the problem of anagenesis and the apparent high degree of morphological conservativism of the nominotypical genus. This conservativism is evidenced by the striking similarity of the *L. niger* clade with the Baltic amber fossils and the Cretaceous genus †*Kyromyrma*. Should the Baltic amber taxa be split, even though there are not enough morphological features to differentiate them from *Lasius* as currently delimited? Or should the phenotypically distinct genus *Myrmecocystus*, like *Acanthomyops* before it, be synonymized with *Lasius*? Perhaps it will be necessary to split the Baltic taxa and to revive *Acanthomyops* for the *Lasius flavus* clade. For now, we acknowledge the difficulty of justifying our choice to retain the Baltic amber fossils in the genus *Lasius* given the principle that all higher taxa are to be monophyletic. We also emphasize that the unexpectedly young age of the crown *Lasius* genus group is not likely to be an artifact of dating method or taxon sampling. We hypothesize that the overall short branch lengths of the *Lasius* genus group are a result of increased species turnover associated with colonization of and radiation into the recently emerged North temperate ecosystems (Economo *et al*. 2018). The young crown group age of *Lasius* and *Myrmecocystus* in turn likely contributes to lack of morphological divergence and makes these ants known examples of taxonomically difficult genera (*e.g.*, Snelling 1976, Seifert & Galkowski 2016).

With respect to our evolutionary questions, we reconceive the ancestral formicine as having originated around the end of the Early Cretaceous or beginning of the Late Cretaceous with anteriorly set eyes, narrowly set coxae, circular propodeal spiracles, lacking a sulcus posterad the helcial sternite, and having low anterolateral corners of abdominal tergum III. Crucially, we find support for the “sepalous” condition of the proventriculus as an autapomorphy of the Formicinae, suggesting that greater control of the crop valve may have been a key innovation in the origin and subsequent radiation of the subfamily during the Mesozoic. In this evolutionary morphological context, the Lasiini derived wide-set coxae and raised gastral corners independently of the Plagiolepidini and some Melophorini, the former in the Late Cretaceous and the latter probably during the Tertiary period. We find that the new genus, *XXX*, is a probable relict of an ancient clade which split from its common ancestor with the tropicopolitan radiation of the *Prenolepis* genus group during the Late Cretaceous.

To date, our study provides the best-supported hypothesis of phylogenetic relationships among the major lineages of *Lasius* and, in addition to revising the classification of the Lasiini, contributes to a greater understanding of the tempo and mode of formicine evolution. From the phylogenetic perspective, we conclude that inclusion of a clock model in analyses of morphology results in substantial topological improvement, and that divergence dating analyses should, if possible, include both morphological and molecular data in the tip-dating framework. Despite the advances we propose, much work remains to be done for the Lasiini. Extended morphological data are required to resolve the placements of fossils more completely. Such analyses may benefit from the inclusion of males (*e.g.*, Barden *et al*. 2017a) and characters specific to the *Prenolepis* genus group. Our work has revealed that two fossil taxa were misplaced at genus rank (†*Lasius pumilus*, now in †*XXX* **gen. nov.**, and †*Pseudolasius boreus*, now †*XXX* **gen. nov.**), as were two fossil taxa misplaced to tribe rank (†*Glaphyromyrmex* and †*XXX* **gen. nov.**). We have great hope for the future of holistic phylogenetic studies as ever more genomic and phenomic data are made available.

## Supporting information

All data sets, as well as input and output of all the analyses mentioned in the *Methods* section are deposited on Dryad (https://doi.org/10.25338/B8B645).

Table S1. List of primers used to generate new sequence data.

Table S2. GenBank accession numbers for all previously and newly published sequences.

Table S3. PartitionFinder “best scheme” for the 9-gene matrix.

Appendix S1. Morphometric differentiation of *Lasius*, *XXX*, and †*XXX*.

Appendix S2. Morphological state definitions used for the 114-character matrices.

Fig. S1. Constraint schemes used for the stepping-stone topology tests in MrBayes.

Fig. S2. The results of the 55-taxon concordance factor analysis.

Fig. S3. The results of the 154-taxon morphology-only MrBayes analysis.

Fig. S4. The results of the 135-taxon molecular-only MrBayes analysis.

Fig. S5. The results of the 135-taxon molecular-only IQ-TREE analysis.

Fig. S6. The results of the 154-taxon combined morphology & molecular MrBayes analysis without a clock model.

Fig. S7. The results of the 154-taxon combined morphology & molecular MrBayes analysis with a uniform IGR clock.

Fig. S8. The results of the 154-taxon combined morphology & molecular MrBayes analysis with a uniform TK02 clock.

Fig. S9. The results of the 154-taxon combined morphology & molecular MrBayes analysis with a FBD IGR clock.

Fig. S10. Ancestral state estimation results for relative compound eye position.

Fig. S11. Ancestral state estimation results for coxal separation.

Fig. S12. Ancestral state estimation results for ventral sulcus posterad helcium.

Fig. S13. Ancestral state estimation results for propodeal spiracle shape.

Fig. S14. Ancestral state estimation results for proventriculus form.

Fig. S15. Ancestral state estimation results for abdominal segment III conformation.

Fig. S16. Ancestral state estimation results for temporary social parasitism.

Fig. S17. Ancestral state estimation results for fungiculture.

Fig. S18. Dot plot of relative eye position data among *Lasius*, *XXX*, and †*XXX*.

Fig. S19. Plots of eye position indices among *Lasius*, *XXX*, and †*XXX*.

Fig. S20. Plot of relative eye size contrasting *Lasius* clades with *XXX* and †*XXX*.

## Acknowledgments

We extend our primary gratitude to Phil Ward. Phil provided us with funding, specimens, sequences, constructive discussion, and attention to the manuscript; as well, he recommended the name *XXX*, which we appreciate. We also appreciate the critique of an anonymous reviewer of our first manuscript, as this led to our discovery of the placement of *myrmidon* and morphological reevaluation of the Lasiini. Thanks to Dave Grimaldi for granting access to the amber collection at the AMNH. For his exquisite automontage images of Baltic amber material and for sharing his insights into the fossil record of the Formicidae, we thank Vincent Perrichot. Brian Fisher and Michele Esposito: Thank you for AntWeb and for maintaining Bolton’s catalog and Ward *et al*.’s AntBib on AntCat. We thank Barry Bolton and Brian Fisher for commenting on the manuscript prior to submission. We thank Bonnie Blaimer for discussion of the Formicinae and for sending us the UCE single-locus gene trees for coalescence analysis. We thank Adrian Richter for discussing head characters. Jill Oberski caught some mistakes in the manuscript and translated Mayr’s writing for us; thank you. Finally, we thank Lech Borowiec and Sebastian Salata for contributing specimens of *XXX myrmidon* and *Lasius carniolicus*, respectively.

## Supplementary material

**Table S1.** Primers used for amplification and sequencing *abdominal-A* (*abdA*), *Arginine Kinase* (*ArgK*), *rudimentary* (*CAD*), *elongation factor 1* α *F2 copy* (*EF1*α*F2*), *long-wavelength rhodopsin* (*LW Rh*), *Topoisomerase I* (*Top1*), *Ultrabithorax* (*Ub*), *wingless* (*wg*). Blank spaces every three nucleotides delimit codons, except for intronic sequences, which are arbitrarily presented with spaces every three nucleotides. Most frequently used primers denoted by asterisk.

**Table S2.** GenBank data for all sequences used in this study.

**Table S3.** PartitionFinder best schemes for the Lasiini_w_outgroups_135t and Lasiini_55t datasets.

## Appendix S1. Morphometric differentiation of Lasius, XXX, and †XXX

Because eye position and size are important for the classification of the Lasiini, we measured six variables for representatives of all *Lasius* species groups plus *XXX myrmidon* and †*XXX pumilus*. Measurements were taken using a digital dual-axis stage micrometer (Mitituyo) beneath a Leica MZ7.5 (50 × maximum magnification); two specimens of *M. myrmidon* and the specimen of †*E. pumilus* were measured from digital micrographs using Adobe Illustrator. All measurements, except for those of †*E. pumilus*, were recorded to three significant figures. An effort was made to measure two individuals of each species. In total, measurements were taken for 104 specimens representing 53 species of *Lasius*, four specimens of *XXX*, and one specimen of †*XXX*. From the six measurements recorded for each specimen, we calculated two metrics each of eye length, eye midlength location, plus the averaged eye length. With the measurements and transformations, seven indices were calculated in order to control for size variation among the measured taxa. We used *ggplot* in R to visualize the results of these analyses.

### Measurements

HL1 *Head length 1*. Length of head in full-face view from posteromedian margin of head to an imaginary line drawn between the anterolateral clypeal corners.

HL2 *Head length 2*. Length of head in full-face view from posteromedian margin of head to anteromedian point of clypeus.

ED1 *Eye distance 1*. Distance from an imaginary line drawn between the anterolateral clypeal corners and the anterior margins of the compound eyes.

ED2 *Eye distance 2.* Distance from an imaginary line drawn between the anterolateral clypeal corners and the posterior margins of the compound eyes.

ED3 *Eye distance 3.* Distance from the anteromedian point of the clypeus to the anterior margins of the compound eyes.

ED4 *Eye distance 4*. Distance from the anteromedian point of the clypeus to the posterior margins of the compound eyes.

### Transformations

EL1 *Eye length 1.* ED2-ED1.

EL2 *Eye length 2*. ED4-ED3.

ELx *Eye length average*. (EL1+EL2)/2.

EML1 *Eye midlength 1*. ELx+ED1.

EML2 *Eye midlength 2*. ELx+ED2.

### Indices

ESI *Eye size index*. ELx/HL2*100.

ClEaI *Lateroclypeus to eye anterior index*. ED1/HL1×100.

ClEpI *Lateroclypeus to eye posterior index.* ED2/HL1×100.

ClEmI *Lateroclypeus to eye midlength index*. EML1/HL×100.

CaEaI *Anteroclypeus to eye anterior index.* ED3/HL2 ×100.

CaEpI *Anteroclypeus to eye posterior index*. ED4/HL2 ×100.

CaEmI *Anteroclypeus to eye midlength index*. EML2/HL2 ×100.

### Discussion

We observe that the compound eyes of *XXX* and †*XXX* are clearly set in a more-anterior position relative to *Lasius* (Fig. S18). Our calculated indices of eye position (Fig. S19) demonstrate that the midlength of the compound eyes of *XXX* and †*XXX* are set anterior to head midlength as measured from the anterolateral clypeal margins (Fig. S19F) and at about midlength as measured from the anterior clypeal margin (Fig. S19G). Because of variation in eye length, we observe overlap in indexed measurements to the posterior eye margin between *XXX* and *Lasius* (Figs. S19A, C) and near overlap of indexed measurements to the anterior eye margin between †*XXX* and *Lasius* (Figs. S19B, D). We also observe that although the two clades of *Lasius* approach *XXX* with respect to relative eye size, they do not overlap, in contrast to the relatively large-eyed †*XXX*. Finally, we observe that the compound eyes of the *L. niger* clade are proportionally larger than those of the *L. flavus* clade (Fig. S18, S20), with some exceptions (*e.g.*, *L. pallitarsis*, very small individuals). Having reviewed the patterns, we conclude that the morphometric data provide quantitative support for our qualitative definitions of the three genera recognized in our work, as well as the two primary clades of *Lasius*.

## Appendix S2. Character list used in combined phylogenetic analysis

All character states were scored as Boolean values, with states defined as TRUE (1) or FALSE (0). Characters were composed and scored independently from that of Maruyama *et al*. (2008), a study which itself expanded on Janda *et al*. (2004). Note that character descriptions of the head assume prognathy.

### Body size proxy

1. Head width ≥ 1 mm. — *Character derived from Peeters & Ito (2015) wherein head width was used as a proxy for body size in argumentation that miniaturization was a key innovation in the early evolutionary history of the Formicidae*.

Compound eyes and ocelli.

2. Medial eye margins strongly converging, rather than parallel. — *The state of anteriorly converging compound eyes has evolved a number of times in the Formicidae, including among the formicoids the genera* Turneria, Santschiella, Opisthopsis, Gesomyrmex*, and* Myrmoteras. *Used previously by Bolton (2003) to define the Gesomyrmecini, including* Gesomyrmex *and* Santschiella.
3. Eye situated at or posterior to head midlength (if FALSE, then situated anterior to head midlength). — *Note that the scoring for this state is different from that used for ancestral state estimation (Fig. S10)*.
4. Eye hypertrophied (if FALSE, then small, reduced, or absent). — *Eyes scored as enlarged for the pseudomyrmecine terminals (although not the case for the whole subfamily),* Gesomyrmex, Gesomyrmex*, and* Myrmoteras; *this state also applies for* Opisthopsis*, which is not in the matrix*.
5. All three ocelli present. — *Ocellar presence is a variable state, even among taxa where they usually occur. For this analysis, if all three ocelli were observable in a worker specimen, the state was scored as TRUE*.

### Mandibles

6. Mandibular teeth in form of massive triangles or spines. — *Scored as TRUE for* Myrmoteras*; teeth enlarged and nearly spine-like in* Myrmecia.
7. At least some mandibular teeth in form of fine serration. — *Applies primarily to Dolichoderinae, which were not heavily sampled*.
8. Smaller teeth uniformly intercalated among larger teeth along margin. — *Applies primarily to Dolichoderinae, which were not heavily sampled*.
9. Mandible, third tooth from apex size as large as other teeth (*i.e*., not reduced). — *Bolton (2003) inferred that a small third tooth from the apex was probably an ancestral state for the Formicinae, with an enlarged tooth being a synapomorphy for the Camponotini*.
10. Basal mandibular tooth offset from rest of teeth on masticatory margin. — *The tooth at the juncture of the basal and masticatory margins is either oriented similarly to other teeth on the masticatory margin or is set more proximally or is at a distinct angle; the latter two states constitute “offset”, as used by Wilson (1955) for classification in* Lasius.
11. Basal mandibular margin with at least one tooth. — *Excludes basal tooth of masticatory margin. A tooth or teeth on the basal mandibular margin occur with some frequency in the myrmechoderine clade (myrmeciomorphs + dolichoderomorphs). Such teeth occur sporadically in the Formicinae, and only in the formicoform radiation*.
12. Mandible elongate, with lateral and masticatory margins parallel or nearly parallel. — *This state describes the sampled Myrmeciinae*.
13. Masticatory mandibular margins meeting in opposition, *i.e.*, not overlapping when mandibles fully closed. — *Mandibles with non-overlapping masticatory margins are observed in* †Prionomyrmex, Nothomyrmecia, *and various trap-jaw ants (Bolton 1999; character 10 of Ward & Brady 2003)*.
14. Mandible with basal and masticatory margins rounding evenly into one another. — *This state occurs sporadically among the sampled core formicoids and has been used previously by Shattuck (1995) as his character 14 for the Dolichoderinae*.
15. Mandibular basal and masticatory margins meeting at a strongly oblique angle. — *Observed among sampled taxa primarily among the* Prenolepis *genus group and some Plagiolepidini, where the juncture between the basal and masticatory margins is clearly defined as an angle*.
16. Groove present on dorsal mandibular surface. — *A dorsal mandibular groove is widespread in the Aculeata (Buren 1970; Hermann* et al. *1971) and is known from various Formicidae (Ettershank 1966; Gotwald 1969; Hermann* et al. *1971). Here, the groove is observed principally in Formicinae*.

### Other mouthparts

17. Prementum concealed by labrum anteriorly, maxillary stipes laterally. — *Autapomorphy of the Dorylinae (as the “dorylomorph subfamilies” in Bolton 2003)*.
18. Maxillary palps elongate, exceeding occipital foramen. — *Scoring differs here from that of character 11 in Maruyama et al. (2008)*.

### Perioral sclerites

19. Hypostoma with lateral flanges which are conspicuous in profile view. — *A defining feature of* Dolichoderus *(Shattuck, 1992, 1995)*.
20. Clypeus, in full face view, with longitudinal carinae at the lateral margins. — *Described by Ward & Brady (2003) as the lateral clypeal carina in their character 6, after Baroni Urbani (2000, character 3). Such carinae are characteristic of †*Prionomyrmex *and* Nothomyrmecia.
21. Pleurostomal condyle conspicuously large and rectangular. — *A newly defined state which is characteristic of the dolichoderomorphs (Aneuretinae + Dolichoderinae)*.

### Anterior clypeal margin

22. Medioclypeus (= “middle lobe” of clypeus, or “discal clypeus” *sensu* Serna & Mackay 2010, Serna *et al*. 2011) produced anteriorly relative to lateroclypeus (= “lateral lobes” of clypeus, or “premalar space”, *sensu* Serna & Mackay 2010, Serna *et al*. 2011). — *This state is observed variably across the Formicidae*.
23. Anterior clypeal margin with distinct, lateromedially narrow lobate anteromedian process, which is with or without additional processes such as teeth. — *Anterior clypeal processes have evolved a number of times in the crown Formicidae, including the Pseudomyrmecinae, various Myrmicinae, and various Formicinae*.
24. Anterior portion of clypeus broadly produced anteriorly, whether convex or truncate. —
25. Anterior clypeal margin crenulate (i.e., with > 2 teeth). — *Crenellation is observed in various Formicinae,* Tetraponera*, and Ponerinae*.
26. Anterior clypeal margin with median notch or emargination. — *Notches or emargination of the anterior clypeal margin occurs with some frequency throughout the family*.
27. Anterolateral notches for reception of labrum present on anterior clypeal margin (visible in full-face view). — *Observed in the* Prenolepis *genus group*.

### Frontoclypeal complex and toruli

28. Antennal toruli directed dorsally (if FALSE, then toruli directed more-or-less laterally). ‒ *When the antennal insertion is treated as a plane defined by the medial and lateral torular arches, the insertion is directed dorsally, rather than laterally in the Dorylinae, which is also more-or-less the case for the Leptanillinae*.
29. Clypeus conspicuously extending posteriorly between antennal toruli by at least 1/4 anteroposterior torulus length. — *States 24, 25, and 26 could be treated as three discrete categories in a single multistate character which would be a measurement of the distance of the antennal toruli relative to the posterior clypeal margin. State 24, here, is observed generally in the Dorylinae, Dolichoderinae, and Myrmicinae. The toruli of most Formicinae either indent the clypeus (state 25) or are considerably distant from the posterior clypeal margin (state 26). Within the Formicinae, some Plagiolepidini have the clypeus extending between the toruli*.
30. Antennal toruli nearly contacting, contacting, or indenting posterior clypeal margin. — *Characterization of this state drawn from Bolton (1994, 2003)*.
31. Antennal toruli considerably distant from posterior clypeal margin (if FALSE, then closely approximated, abutting, or indenting). — *As for character 25, characterization of this state is drawn from Bolton (1994, 2003) for definition of the Camponotini. This state is also observed in* Myrmoteras, Oecophylla*, and* Notostigma.
32. Antennal sockets in full-face view: fully exposed. — *States 33, 34, and 35 could be treated as three discrete categories in a single multistate character. When viewed from dorsal aspect, the antennal sockets may be concealed to some degree by either the medial torular arch (state 34) or the frontal carinae (state 35). Among sampled taxa, fully exposed sockets are observed in* Vicinopone, †Procerapachys*, and a* Polyrhachis *species*.
33. Antennal sockets in full-face view: partially to completely concealed by laterally produced medial torular arches (or fused carinae + medial torular arches). — *Most taxa observed in this study had antennal sockets which were partially concealed by the medial torular arches*.
34. Antennal sockets in full-face view: partially to completely concealed anteriorly produced frontal carinae, which also conceal the medial torular arch (this state represents presence of true “frontal lobes”). — *Most Dorylinae scored for this study had expanded frontal carinae which conceal, at least in part, the antennal sockets. Such expansion is also observed in various Myrmicomorpha (ectaheteromorphs + Myrmicinae) and a few Formicinae*.

### Facial surfaces

35. Frontal carina present. — *Frontal carinae are present in the majority of formicoids and, among taxa included in this study, are absent for a few Formicinae*.
36. Frontal carina conspicuous, being > 0.5 × anteroposterior antennal torulus diameter. — *Relatively long frontal carinae occur in most sampled taxa. In addition to* Cladomyrma*, a number of core Formicinae have reduced carinae*.
37. Frontal carina carinate, continuing as a sharp ridge until posterior terminus (if FALSE, then carina ecarinate). — *Rounded frontal carinae occur among the Lasiini and various other Formicinae here sampled. Reduction of the length of the frontal carinae (state 37, above) is not always associated with loss of carination*.
38. Frontal carinae very closely approximated on face. — *Among taxa sampled in the present study, very close-set frontal carinae are only observed in* Pseudomyrmex osurus *and †*Pseudomyrmex vicinus.
39. Parafrontal carina (genal carina laterad antennal torulus) present. — *Among the formicoids, parafrontal carinae are observed in Dorylinae and the myrmicine* Terataner.

### States of relative antennomere length

40. Scape length (SL) < 0.5 × head width (HW). — *In addition to geniculation, one of the traditional diagnostic features of the Formicidae is scapes which are long relative to the flagellum (*e.g.*, Wilson* et al. *1967a,b; Bolton 2003). Here, we evaluate the overall length of the scape by comparison to the cranium, and relative length to the flagellum separately. The former continuous is divided into three mutually exclusive, additive states*.
41. Scape length (SL) > 0.5 × head width (HW), < 1.0 x HW or head length (HL). — *See character 40*.
42. Scape length (SL) > head width (HW) or head length (HL). — *See character 40*.
43. Scape length > 0. 5 × funiculus (= pedicel + flagellum) length. — *See character 40*.
44. Antennomere 3 longer than antennomere 4. — *Variable among the Formicidae, with especially long third antennomeres observed in stem Formicidae (*e.g.*, †*Sphecomyrminae).
45. Antennomere 3 broader than long. — *Observed in* XXX *and †*XXX.

### Sensilla basiconicum on flagellum

46. Flagellum: Sensilla basiconicum socket raised above antennomere (if FALSE, then flush). — *Recorded as an autapomorphy of the myrmeciomorphs (Myrmeciinae, Pseudomyrmecinae) by Bolton (2003)*.

### Cranial setation

47. Coarse, paired macrosetae present on face. — *This setational pattern is observed in various* Prenolepis *genus group taxa, as well as* Anoplolepis gracilipes.
48. *Standing* setae present on dorsum of head, excluding posterolateral corners and clypeus. ‒ *This is equivalent to character 26 of Maruyama et al. (2008). Our scoring differs for the* L. fuliginosus *group: We observe dorsal setae near the posterior margin of the head in* L. fuliginosus *and* L. spathepus*, whereas Maruyama* et al. *recorded these setae as absent for these two species*.
49. Ventral head surface margined laterally and posteriorly by a continuous line of long, curved setae (psammophore present). — *Ventral cranial psammophores have evolved a number of times in the Formicidae, notably in* Pogonomyrmex *and* Veromessor *(Myrmicinae),* Melophorus *(Formicinae), and* Myrmecocystus.

### Mesosomal shape

50. Pronotum with dorsolaterally situated longitudinal margination. — *Pronotal margination has evolved several times across the Formicidae*.
51. Promesonotal articulation immobile (sutured, whether suture externally visible or not). ‒ *Among sampled taxa, promesonotal fusion is observed in Dorylinae, Ectatomminae, and Myrmicinae. See also Bolton (2003) and Bolton in Fisher & Bolton (2016)*.
52. Mesosomal profile diagonal, with anterior portion (pronotum and mesonotum) raised, often convex or strongly humped (if FALSE then mesosoma more-or-less linear, with a boxy shape and profile). — *The TRUE state is modified from Bolton’s (2003) conception of the “domed” promesonotum, used to define his Pheidolini, particularly in contrast with the “boxy” form used to define his Myrmicini*.
53. Mesonotum in dorsal view conspicuously anteroposteriorly elongate (if FALSE then reduced to narrow, transverse bar, or absent). — Most *sampled taxa have elongate mesonota*.
54. Mesonotum petiolate, being constricted and separating pronotum distinctly from mesopleural area. — *This “petiolation”, or constriction, results in the “hourglass” shape of various* Prenolepis *genus group taxa, as well as a few other Formicinae (*e.g., Oecophylla).
55. Mesoscutellum conspicuous, bar like (if FALSE then reduced). — *The workers of some species have secondarily derived a well-developed mesoscutellum*.
56. Metanotum present as distinct, well-defined, transverse sclerite (if FALSE then may be represented by groove or irregular ridge). — *Although most crown Formicidae lack a well-developed metanotum, as noted by Barden & Grimaldi (2016), the metanotum is secondarily well-developed in some genera*.
57. Metanotal groove distinct and well-developed. Metapleural glands.
58. Metapleural gland present. — *As with scape-to-flagellum length, presence of the metapleural gland is a canonical defining state of the Formicidae (Wilson* et al. *1967a,b; Brothers, 1975; Bolton 2003). The gland has been variably lost in males and has been lost multiple times in the Camponotini*.
59. Metapleural gland with conspicuously enlarged atrium and bulla. — *Newly observed apomorphy of the* Lasius flavus *clade*.

### Propodeum

60. Propodeum armed with spines or other distinct protuberances. — *Propodeal armature has arisen independently among various Formicidae*.
61. Propodeal spiracle: slit-shaped (if FALSE then rounded or elliptical). — *Used by Bolton (2003) to define his formicine tribe group. See also Agosti (1990) and Agosti & Bolton (1990)*.
62. Propodeal spiracle situated at or above 2/3 dorsoventral height of propodeum in lateral view. — *Propodeal spiracle location is informative for defining higher groups of the Formicidae, including the family itself (Bolton 2003, Boudinot 2015). Among the sampled taxa, a “high” spiracle is observed in the Myrmechoderines, the Ectatomminae* sensu lato*, some Myrmicinae, and various Formicinae*.
63. Propodeal spiracle situated at or near posterolateral margin of propodeum (if FALSE, then spiracle situated more anteriorly. — *Although not maximally consistent, a posteriorly-situated spiracle is observed in Camponotini, and Lasiini among other taxa*.
64. Propodeum produced posterodorsally as shelf overhanging posterior face. — *Observed in* Dolichoderus*, and some* Camponotus.
65. Propodeal foramen with thick, dorsal carina. — *This state roughly distinguishes the “lasiiform grade” (Myrmelachistini, Lasiini) from the “formicoform radiation” (Melophorini through Camponotini)*.

### Legs

66. Metacoxa with posterodorsal, proximal spine. — *Scored as an autapomorphy of †*Gnamptogenys levinates*, although such spines also occur in other* Gnamptogenys*. Surprisingly, similar posterodorsal process are also observed in Myrmosidae (*e.g.*, Brothers 1975)*.
67. Metacoxae wideset, with petiolar foramen extending to mesocoxal foramina. — *One of the defining states of Bolton’s (2003) lasiine tribe group, along with a U-shaped ventral petiolar cross-section*.
68. Metatibia with double row of ventral (inner) setae. — *Among sampled taxa, these setae are observed categorically in the Formicini and sporadically in the Camponotini and Melophorini*.
69. Anterior mesotibial spur well-developed (if FALSE spur formula 1 or 0). — *Reduction of the anterior meso- and metatibial spurs has occurred numerous times in the Formicidae (see Appendix 2 of Bolton 2003)*.
70. Anterior metatibial spur well-developed (if FALSE spur formula 1 or 0). — *See comment for prior character*.

### Flexibility of mesosoma

71. Metasoma capable of flexion over mesosoma. — *This state is a defining feature of* Oecophylla *and the myrmicine* Crematogaster.

### Abdominal segment II (“petiole”)

72. Petiole (abdominal segment II) with dorsolateral longitudinal carinae anterior to the node, and which may continue onto node. — *Observed in poneroids, the Dorylinae, some Ectatomminae* sensu lato *and Myrmicinae*.
73. Petiole (abdominal segment II) anterior foramen completely sessile. — *Pedunculation of the petiole is described by this and the following two characters. Sessile petioles are observed in most Formicinae, excluding* Myrmoteras *and* Oecophylla.
74. Petiole (abdominal segment II) anterior foramen subsessile (anterior foramen narrowed, giving petiole an anteriorly extended look in lateral or dorsolateral view).
75. Petiole (abdominal segment II) anterior foramen completely pedunculate (anterior region of segment long, in comparison with posterior region, anterior region narrow).
76. Petiole (abdominal segment II) posterior foramen pedunculate.
77. Petiole strongly inclined anteriorly or with posterior portion elongate. — *This is a feature of those groups which have the “gaster” overhanging the petiole, including the Tapinomini (Dolichoderinae), Plagiolepidini, and* Prenolepis *genus group*.
78. Petiolar node relatively high, reaching or exceeding propodeal spiracle in lateral view. ‒ *While a reduced petiolar node is characteristic of groups with the “prenolepidine syndrome” (*i.e., *those mentioned for the prior state), the petiolar node varies in length throughout the Formicidae. For this reason, TRUE for this state means “tall” in distinction to “short” or “very short”*.
79. Dorsal apex of petiolar node transversely carinate or pointedly angular, whether anteriorly, posteriorly, or at midlength. — *Transverse carination appears sporadically across the Formicidae but is by no means consistent*.
80. Petiolar node anteroposteriorly narrow, thus scale-like. — *The “squamiform” node of various Formicidae*.
81. Petiolar node with dorsal armature (spine or spines present). — *Dorsal armature has arisen infrequently*.
82. Petiolar laterotergite present (if false, then reduced to inconspicuousness or absent altogether). — *This is probably an ancestral feature of the Formicidae which has been lost in parallel several times. Such loss is observed during in taxa with tergosternal fusion*.
83. Petiole (abdominal segment II) tergosternal fusion present. — *This represents the first step toward complete tergosternal fusion*.
84. Petiole (abdominal segment II) tergosternal fusion complete (if FALSE, then may be partial or absent).

### Abdominal segment III

85. Abdominal segment III, transverse sulcus present posterad helcial sternite. — *Bolton (2003) recorded the sulcus as present for the Camponotini, “Notostigmatini”, Formicini, and some “Melophorini”*.
86. Differentiated presternite of abdominal segment III (helcial sternite), in profile, concealed by tergite. — *This and the following two states were described by Bolton (1990a) in his crucial study of the abdominal characters. This state is probably ancestral for the Formicidae*.
87. Differentiated presternite of abdominal segment III (helcial sternite), in profile, exposed, bulging ventrally and overlapped by tergite. — *An autapomorphy of the Dorylinae discovered by Bolton (1990a,b)*.
88. Differentiated presternite of abdominal segment III (helcial sternite), in profile, exposed, continuous with tergite. — *An autapomorphy of the Myrmicinae discovered by Bolton (1990a,b)*.
89. Anterior articulatory surfaces of abdominal segment III infraaxial, i.e., below segment midheight. — *The characterization of this and the following two states was derived from Keller (2011). The state of having an infraaxial helcium is widespread among the crown Formicidae*.
90. Anterior articulatory surfaces of abdominal segment III axial, i.e., at segment midheight. — *In the present taxon set, this state is observed in the Dorylinae and all of the myrmeciomorphs except* Myrmecia.
91. Anterior articulatory surfaces of abdominal segment III supraaxial, i.e., above segment midheight. — *Although supraaxiality occurs in various poneroids, such as most Amblyoponinae, and some Myrmicinae,* Aneuretus *is the only formicoid in present matrix for which the supraaxial state is true*.
92. Abdominal sternum III with raised ridge or bosses (= prora) anteriorly.
93. Abdominal segment III tergum and sternum fused. — *In addition to tergosternal fusion of abdominal segment IV (character 101), tergosternal fusion of segment III is a feature of metasomal “tubulation”,* sensu *Taylor (1978). See also Bolton (1994, 2003) and Fisher & Bolton (2016)*.
94. Abdominal segment III dorsoventrally petiolated (reduced in size, forming “postpetiole”). — *“Petiolation” is a syndrome of modifications which are frequently referred to, but rarely defined. Here, we score Dorylinae,* Myrmecia, *Pseudomyrmecinae, and Myrmicinae as having petiolation of the third abdominal segment. Characters 95 and 96 also pertain to petiolation of AIII*.
95. Abdominal segment III moderately lateromedially narrowed relative to segment IV in dorsal view (*i.e*., AIII width > 0.5 × but < 0.90 × AIV width). — *Lateromedial narrowing of the third abdominal segment relative to the fourth is another feature observed in taxa which display the “petiolization” syndrome. A narrow third segment is observed in the Myrmeciomorpha (Pseudomyrmecinae, Myrmeciinae), and is scored as such. Note that this applies to* Nothomyrmecia*, despite the fact that the third abdominal segment of this genus is conical and lacks a muscular node*.
96. Abdominal segment III strongly lateromedially narrowed relative to segment IV in dorsal view (*i.e*., AIII width < 0.5 × AIV width). — *This is the extreme state of lateromedial narrowing of abdominal segment III associated with the “petiolization” syndrome. This was scored as TRUE for the Myrmicinae*.
97. Abdominal segment III tergosternal margins forming narrow shoulder laterad helcium. ‒ *“Shouldering” of abdominal segment III was used by Bolton (2003) to diagnose various groups of the Myrmicinae (*e.g.*, comment iii of the solenopsidine tribe group), but is also observed in Formicinae, such as the* Prenolepis *genus group and the Plagiolepidini*.
98. Abdominal segment III tergosternal margins raised high above helcium. — *This state, plus the preceding, were used by Bolton (2003) to define his Plagiolepidini*.
99. Abdominal segment III elongate, comprising most of gaster (segments III+) in dorsal view. — *Observed in various Camponotini, particularly* Polyrhachis *(Bolton 1994). This state also applies to groups which were not sampled in the present study, such as various Myrmicinae*.

### Metasoma, posterior to segment III

100. Cinctus of abdominal segment IV defining presclerites (correlated with formation of postpetiole) present. — *In many Formicidae, a given tergum or sternum may be divided into pre- and post-sclerites by a transverse sulcus (cinctus) (Bolton 1990)*.
101. Abdominal segment IV tergosternal fusion present (whether partial or complete). — *A probable synapomorphy of the poneroid clade, with reversal in* Adetomyrma*, tabulated by Bolton in Fisher & Bolton (2016); see also Bolton (2003)*.
102. Spiracles of abdominal segments V–VII exposed (if false, then concealed by preceding tergite). — *An autapomorphy of the Dorylinae (Bolton 1990, 1994, 2003)*.
103. Stridulitrum of abdominal segment IV tergum present. — *Stridulitra occur, with some frequency, on the fifth abdominal tergum throughout the Formicidae*.
104. Stridulitrum of abdominal segment IV sternum present. — *The stridulitrum of the fifth abdominal sternum is a unique derivation of* Nothomyrmecia.
105. Abdominal terga VI and VII with dense, uniform layer of short and strongly curved setae.
106. Sting present. — *A functional sting has been reduced in various Formicidae (Kugler 1978, 1979), including the Attini (*sensu *Ward* et al. *2015) and Crematogastrini (*sensu *Blaimer* et al. *2018). The sting is also absent in the Formicinae and Dolichoderinae, as has been recognized since the classical era of ant taxonomy (Mayr 1868, 1869; Forel 1878)*.
107. Acidopore and coronula present. — *The seventh abdominal sternum in Formicinae is characteristically curved, forming the acidopore, and is usually surmounted by a well-defined ring of setae, the coronula. See Hung & Brown (1966) and Keller (2011)*.

### Larval phenotype

108. Larval food pocket (trophothylax) present. — *Recorded as an autapomorphy of the Pseudomyrmecinae by Bolton (2003)*.

### Polyphenism

109. Major (soldier) caste present. — *Pronounced allometry has evolved many times in the Formicidae. Here, majors were recorded for* Aneuretus, Pheidole*, most Camponotini*, Gesomyrmex, *most Melophorini, and* Pseudolasius.
110. At least some workers with phragmotic face. — *Among sampled formicoids, phragmotic faces occur only in* Colobopsis *(see Ward et al. 2016 for distinguishing features of* Colobopsis).

### Characters used in previous studies

Most setational characters of Maruyama *et al*. (2008; 18–20, 23–29, 34, 35, 64–66, 102) not included, except 26 which relates to presence of cranial setae. Additionally, body color, used to define the *fuliginosus* group (M08: 1, 2) not included.

### *A note on* Santschiella

Bolton (pers. comm.) recently dissected a specimen of the bizarre formicine *Santschiella* and noted that the definition of this genus in Fisher & Bolton (2016) needs to be updated (character states quoted verbatim): **(1)** Mandible has 7 teeth, with tooth 3 reduced (smaller than 4). **(2)** Maxillary palpi are very long and extend back almost to the occipital foramen. **(3)** Metacoxae are closely approximated. **(4)** Propodeal foramen is short. **(5)** Tergite of helcium is entire, as in *Echinopla* and most *Polyrhachis*—the characteristic formicine notch is absent. **(6)** Post-helcial sulcus is present on abdominal segment III.

## Figure legends for supplementary files

**Fig. S1.** Schematics of the constraints used in the topology tests for Bayes factor analysis of the Lasiini_55t dataset. (A) H_0_: *XXX* and *Lasius* genus group monophyletic; H_1_: *XXX* and *Prenolepis* genus group monophyletic (B) H_0_: *Lasius* s. str. monophyletic; H_1_: *Lasius* s. str. paraphyletic (C) H_0_: *Chthonolasius* monophyletic; H_1_: *Chthonolasius* polyphyletic (D) H_0_: *Cautolasius* monophyletic; H_1_: *Cautolasius* paraphyletic.

**Fig. S2.** Results of the concordance factor analysis inferred from the Lasius_55t dataset in IQTREE. Node support values are reported as gene concordance factor (gCF) vs. site concordance factor (sCF).

**Fig. S3.** Results of analysis of the Lasiini_w_outgroups_w_fossils_morphology_154t dataset in MrBayes. Node support values are reported as Bayesian posterior probabilities (PP).

**Fig. S4.** Results of analysis of the Lasiini_w_outgroups_135t dataset in MrBayes. Node support values are reported as Bayesian posterior probabilities (PP).

**Fig. S5.** Results of analysis of the Lasiini_w_outgroups_135t dataset in IQTREE. Node support values are reported as maximum likelihood bootstrap (BS).

**Fig. S6.** Results of analysis of the Lasiini_w_outgroups_w_fossils_154t dataset in MrBayes without a clock model. Node support values are reported as Bayesian posterior probabilities (PP).

**Fig. S7.** Results of analysis of the Lasiini_w_outgroups_w_fossils_154t dataset in MrBayes with a uniform branch length prior and IGR clock rate prior. Node support values are reported as Bayesian posterior probabilities (PP). Horizontal blue bars at nodes are 95% highest posterior density (HPD) intervals.

**Fig. S8.** Results of analysis of the Lasiini_w_outgroups_w_fossils_154t dataset in MrBayes with a uniform branch length prior and TK02 clock rate prior. Node support values are reported as Bayesian posterior probabilities (PP). Horizontal blue bars at nodes are 95% highest posterior density (HPD) intervals.

**Fig. S9.** Results of analysis of the Lasiini_w_outgroups_w_fossils_154t dataset in MrBayes with a FBD branch length prior and TK02 clock rate prior. Node support values are reported as Bayesian posterior probabilities (PP). Horizontal blue bars at nodes are 95% highest posterior density (HPD) intervals.

**Fig. S10.** Ancestral state estimation (*ace*) results from analysis of a 135-taxon chronogram for compound eye location under the equal rates assumption. In the pie charts, red = eyes anteriorly set and blue = eyes set at or posterior to head midlength.

**Fig. S11.** Ancestral state estimation (*ace*) results from analysis of a 135-taxon chronogram for coxal separation under the equal rates assumption. In the pie charts, red = close-set and blue = wide-set.

**Fig. S12.** Ancestral state estimation (*ace*) results from analysis of a 135-taxon chronogram for sulcus presence posteroventral to the helcium. In the pie charts, red = sulcus absent and blue = sulcus present.

**Fig. S13.** Ancestral state estimation (*ace*) results from analysis of a 135-taxon chronogram for propodeal spiracle shape. In the pie charts, red = spiracle circular to elliptical and blue = spiracle slit-shaped.

**Fig. S14.** Ancestral state estimation (*ace*) results from analysis of a 135-taxon chronogram for proventricular form. In the pie charts, red = asepalous, blue = sepalous.

**Fig. S15.** Ancestral state estimation (*ace*) results from analysis of a 135-taxon chronogram for tergosternal suture conformation of abdominal segment III. In the pie charts, pink = without narrow shoulder, blue = low narrow shoulder, red = high narrow shoulder.

**Fig. S16.** Ancestral state estimation (*ace*) results from analysis of a 21-taxon chronogram representing *Lasius* for temporary social parasitism. In the pie charts, red = absence, blue = presence.

**Fig. S17.** Ancestral state estimation (*ace*) results from analysis of a 21-taxon chronogram representing *Lasius* for fungiculture. In the pie charts, red = absence, blue = presence.

**Fig. S18.** Dot plot of longitudinal distance from anterolateral corners of clypeus to posterior margins of compound eyes (ED2) by maximum head length (HL2) contrasting the two primary clades of *Lasius* with the genera *XXX* and †*XXX*, and demonstrating that the compound eyes of the latter two taxa are clearly set more-anteriorly than those of *Lasius*.

**Fig. S19.** Box and whisker plots of values for six eye position indices, showing differentiation among *Lasius*, *XXX*, and †*XXX*.

**Fig. S20.** Box and whisker plot of relative eye size (ESI), the index of average eye length (ELx) proportional to maximum head length (HL2), contrasting the two primary clades of *Lasius* with the genera *XXX* and †*XXX*.

## Notes

### Competing Interest Statement

The authors have declared no competing interest.

