## Supplementary figures and images for "Phylogeny, evolution, and classification of the ant genus *Lasius*, the tribe Lasiini, and the subfamily Formicinae (Hymenoptera: Formicidae)"

### Fig. S1

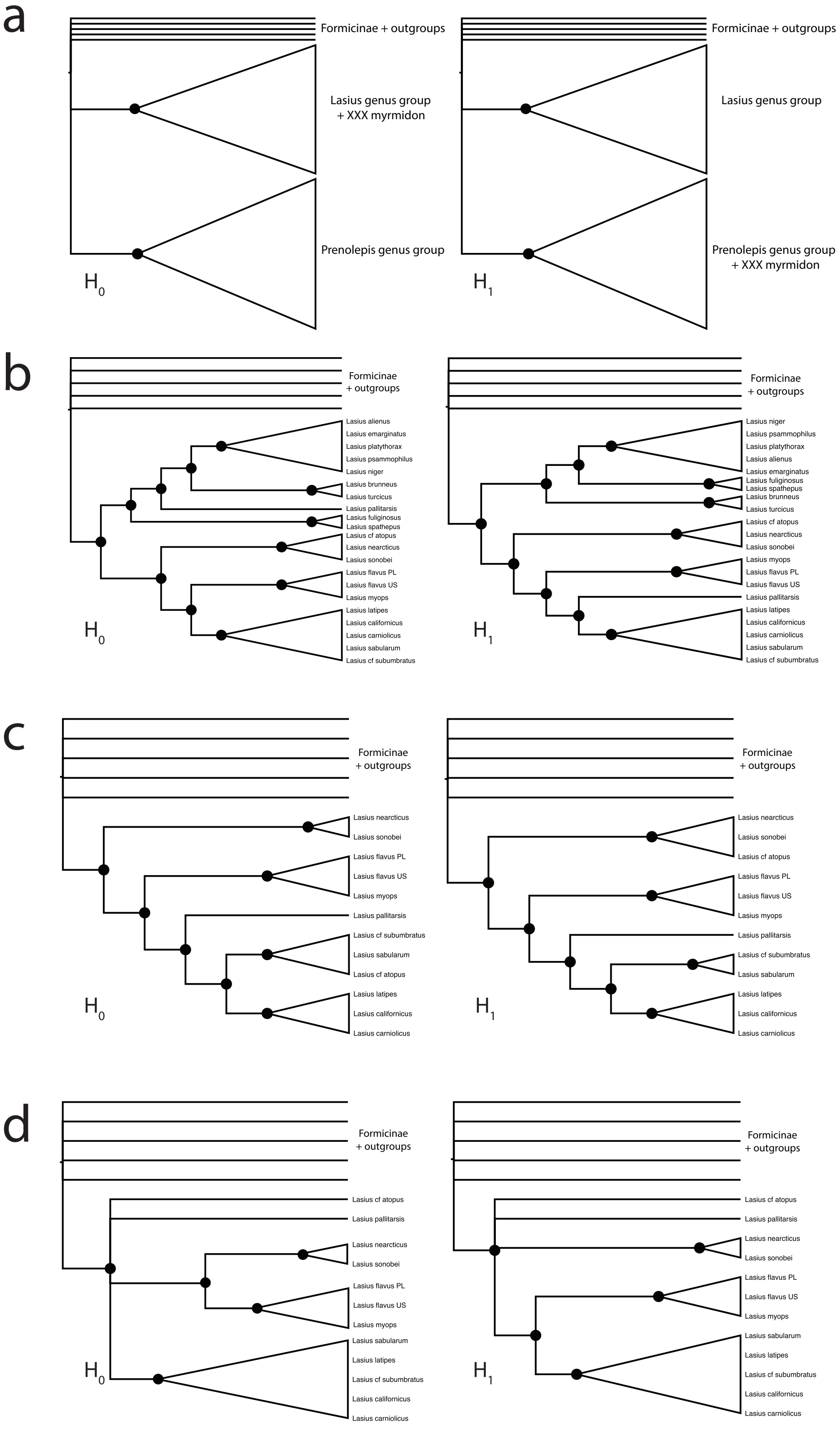

### Fig. S2

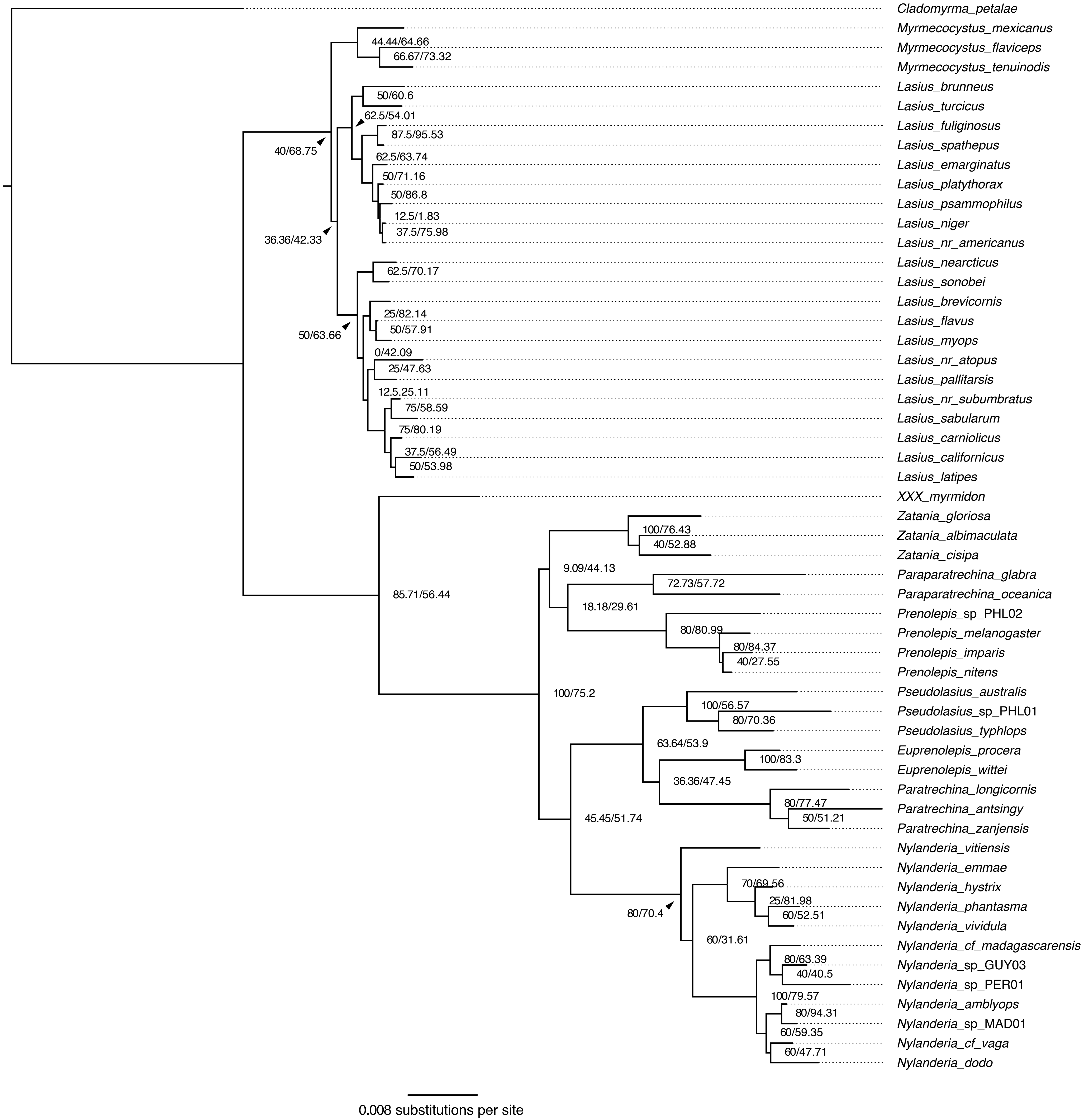

### Fig. S3

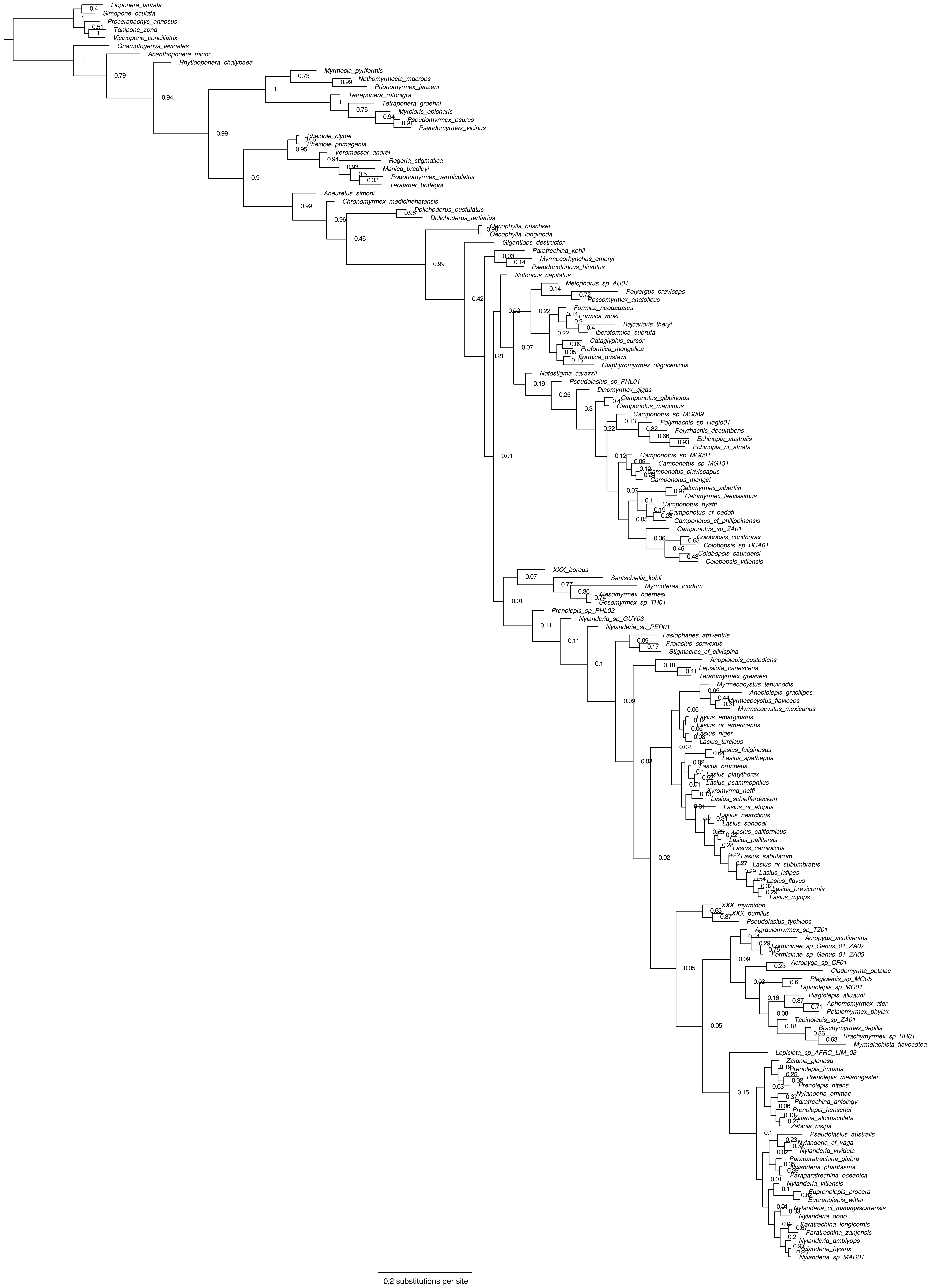

### Fig. S4

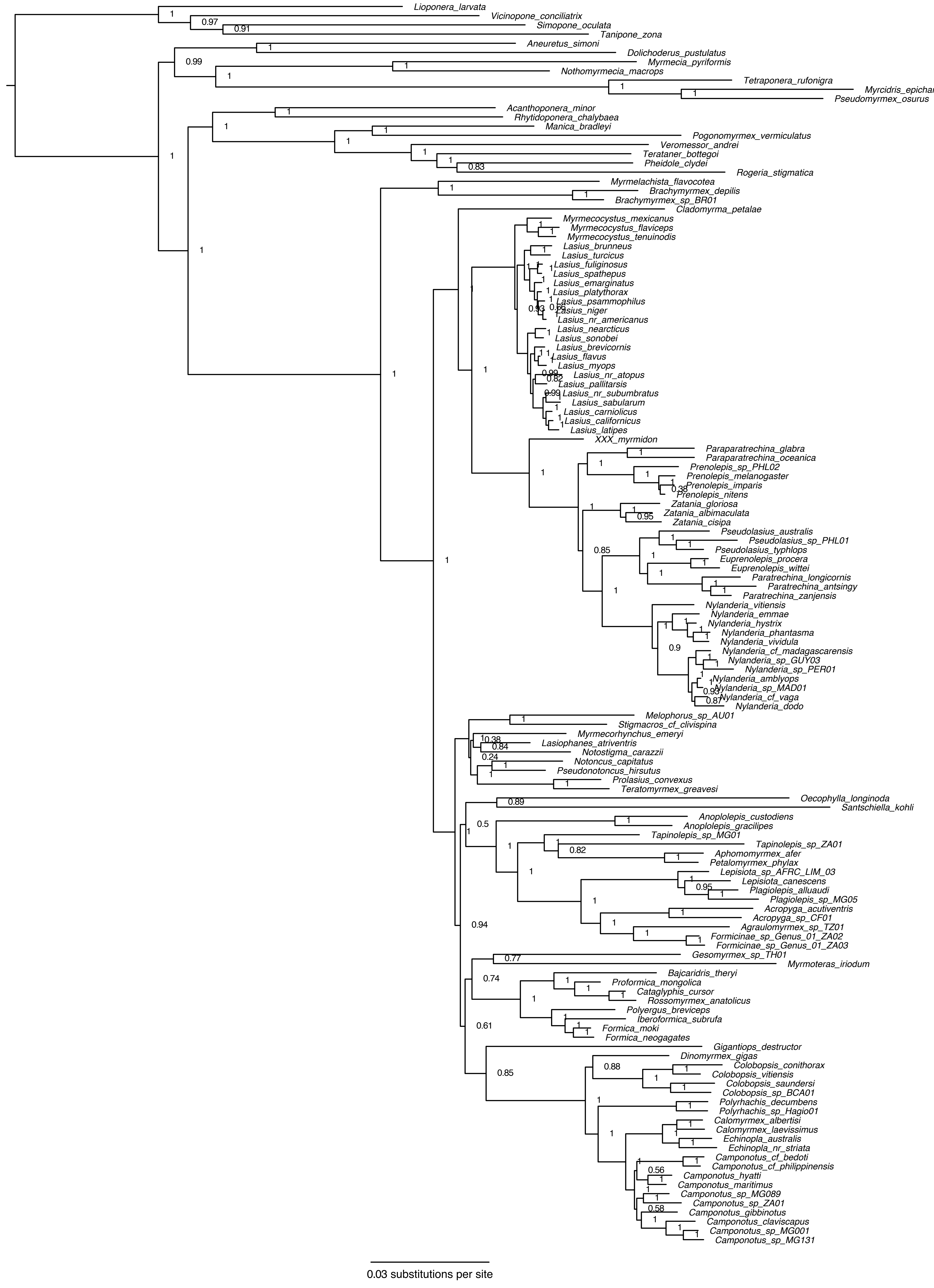

### Fig. S5

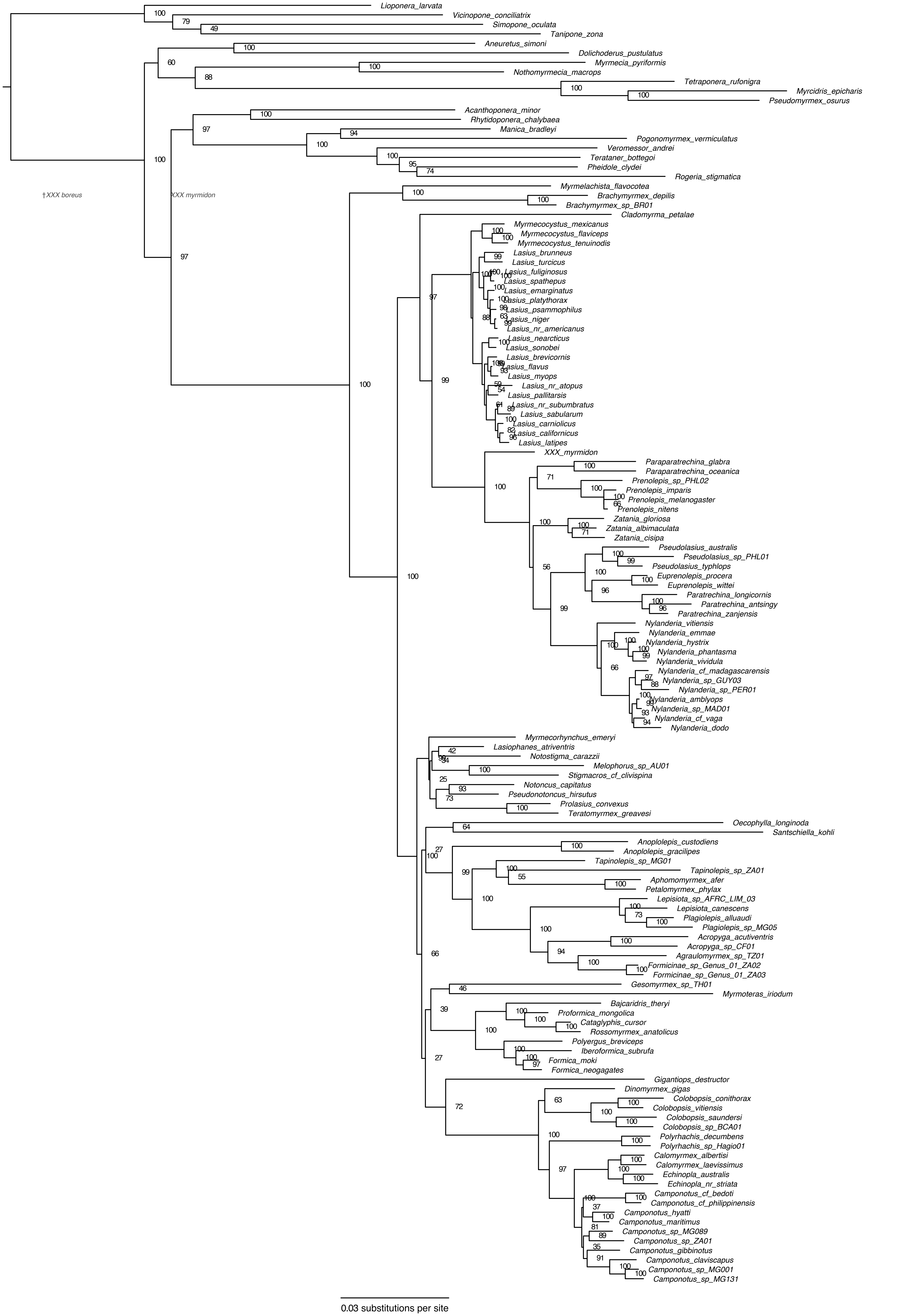

### Fig. S6

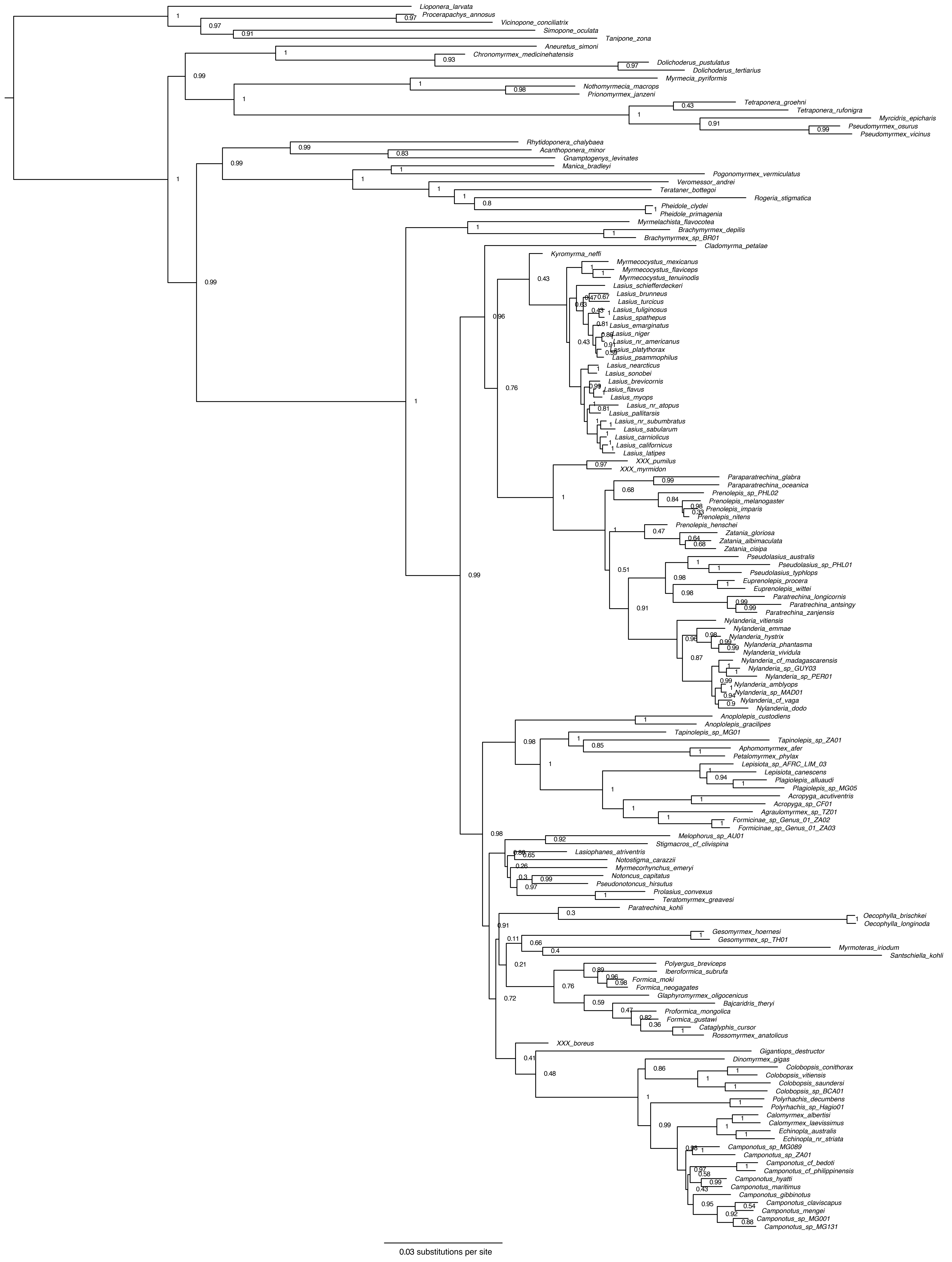

### Fig. S7

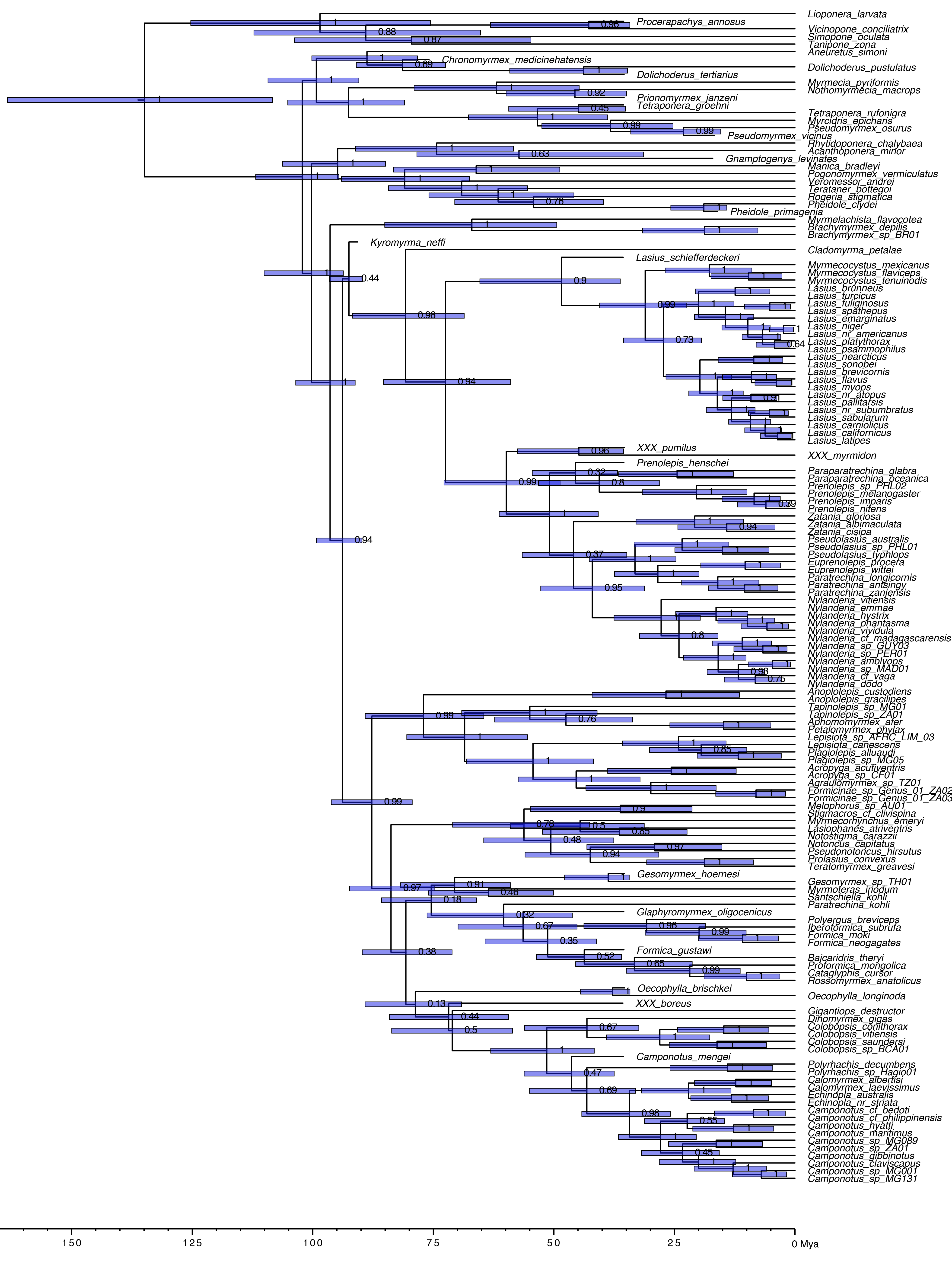

### Fig. S8

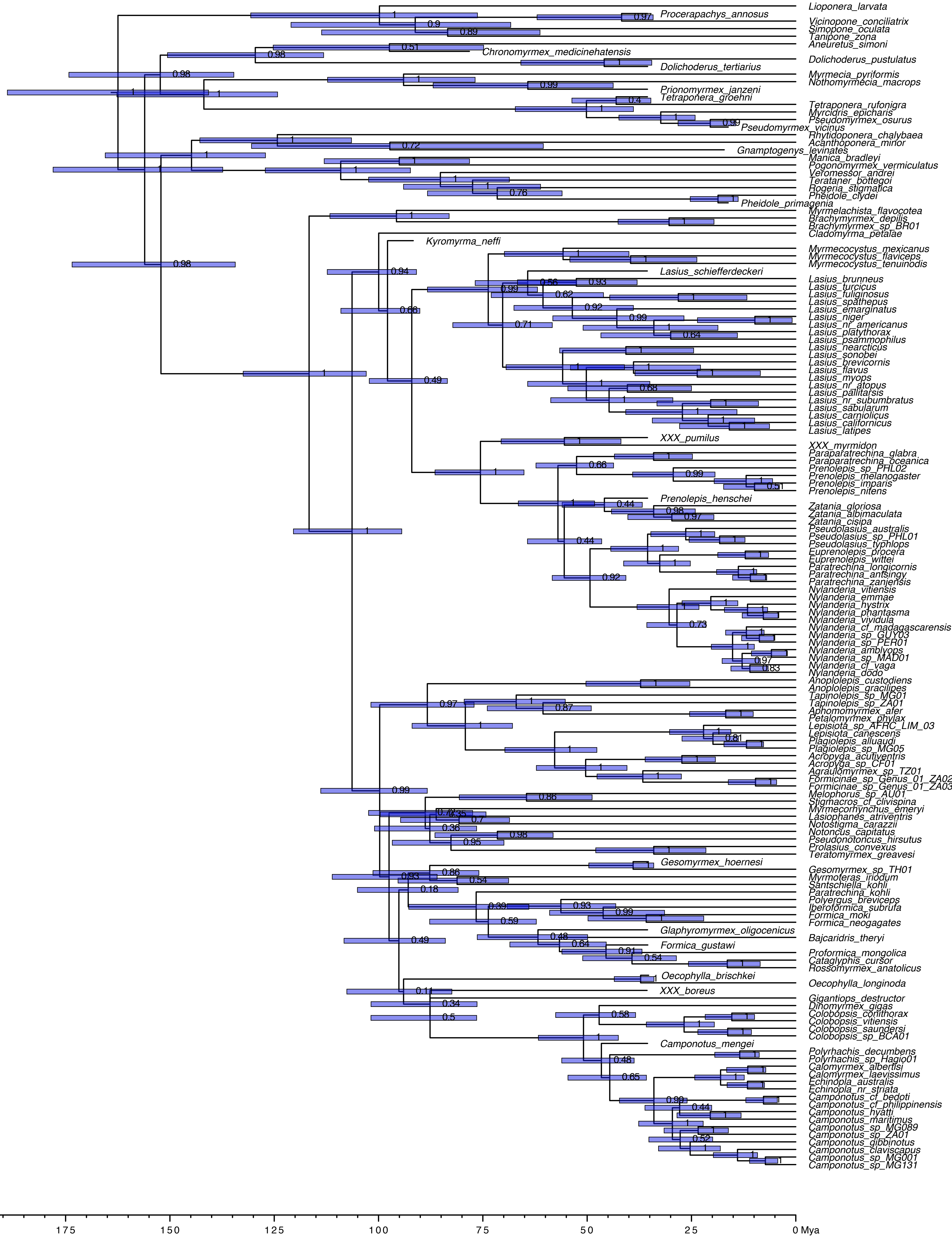

### Fig. S9

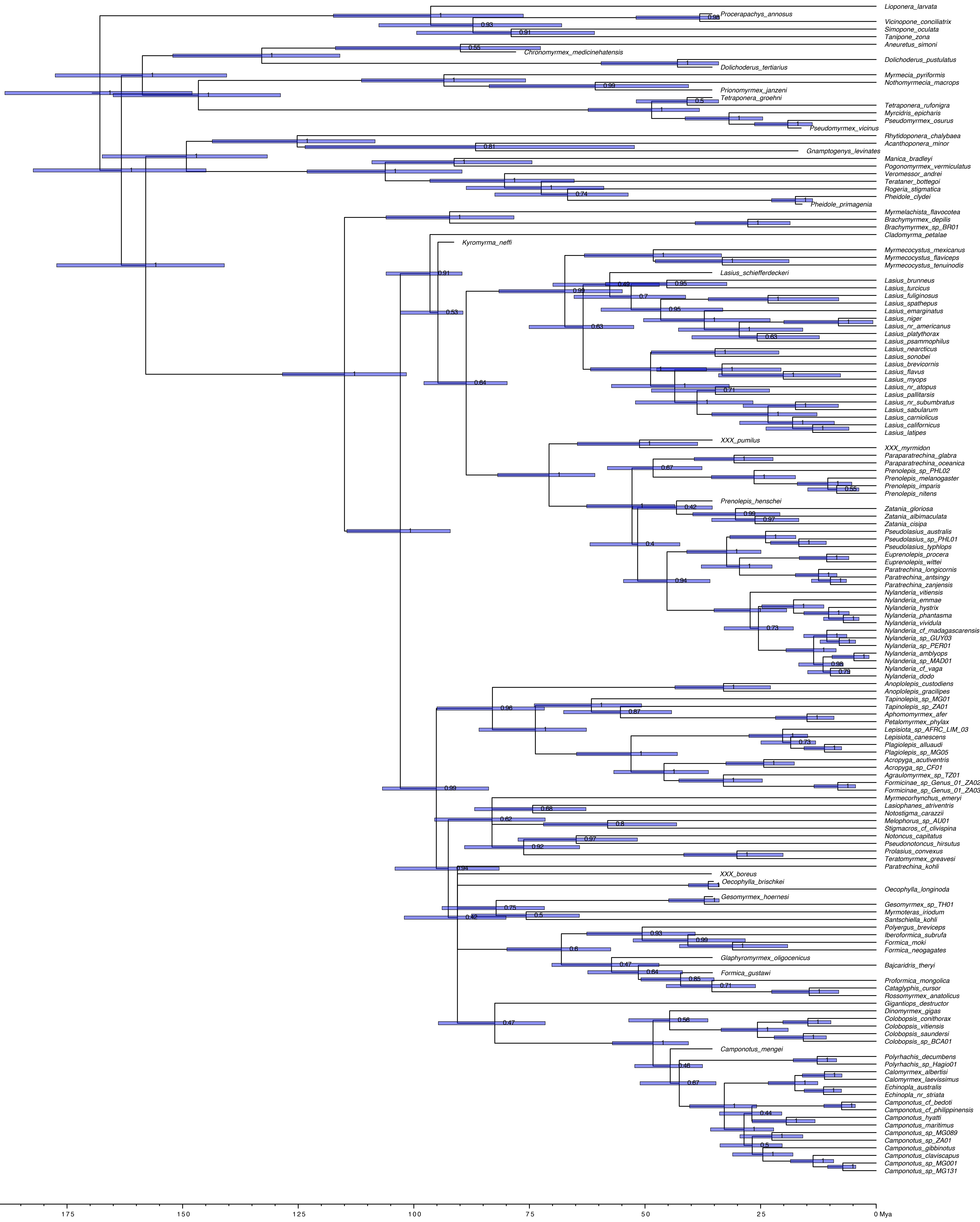

### Fig. S10

#### Lasiini & outgroups 135t Eyes ER ASR

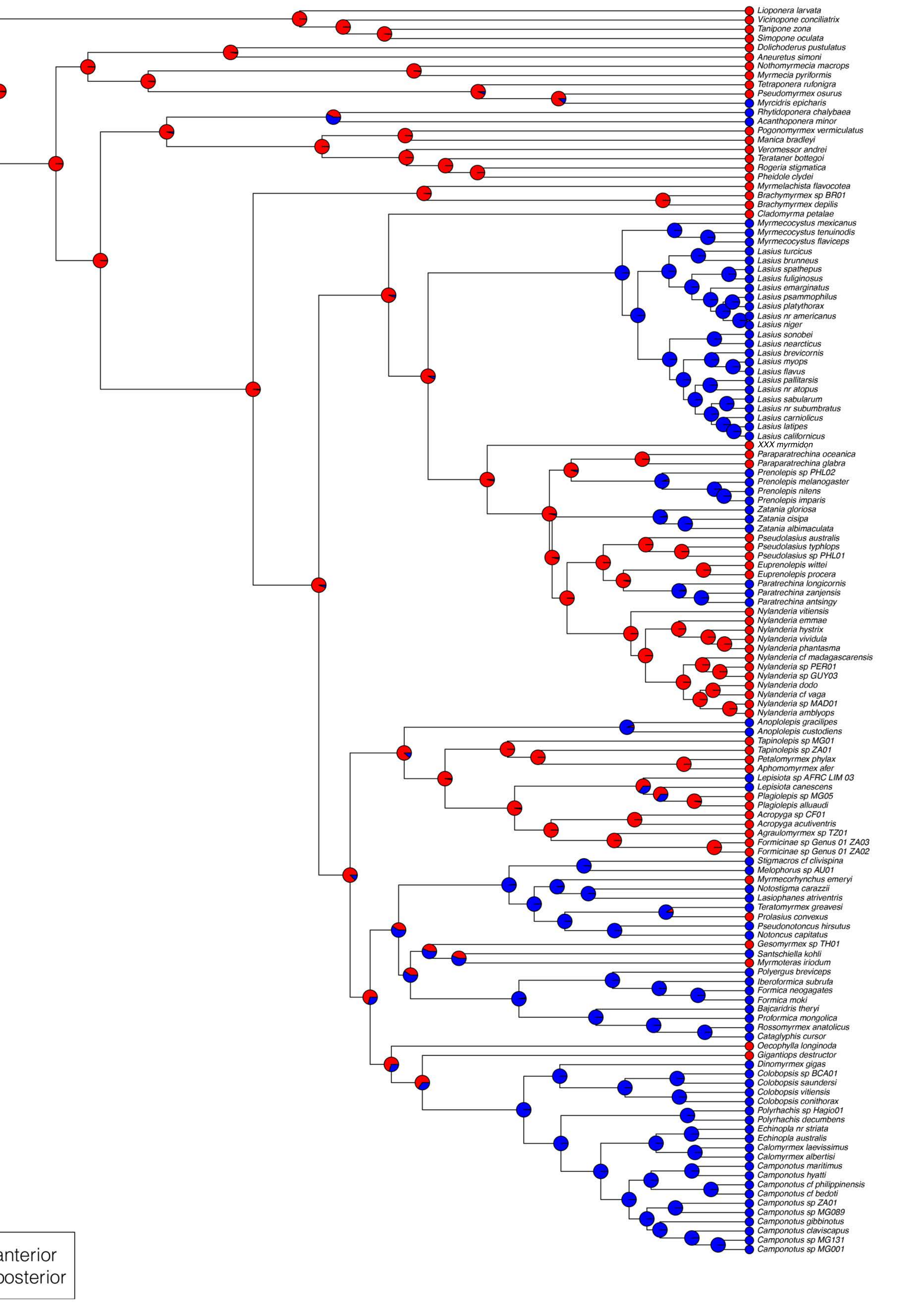

### Fig. S11

#### Lasiini & outgroups 135t Coxae ER ASR

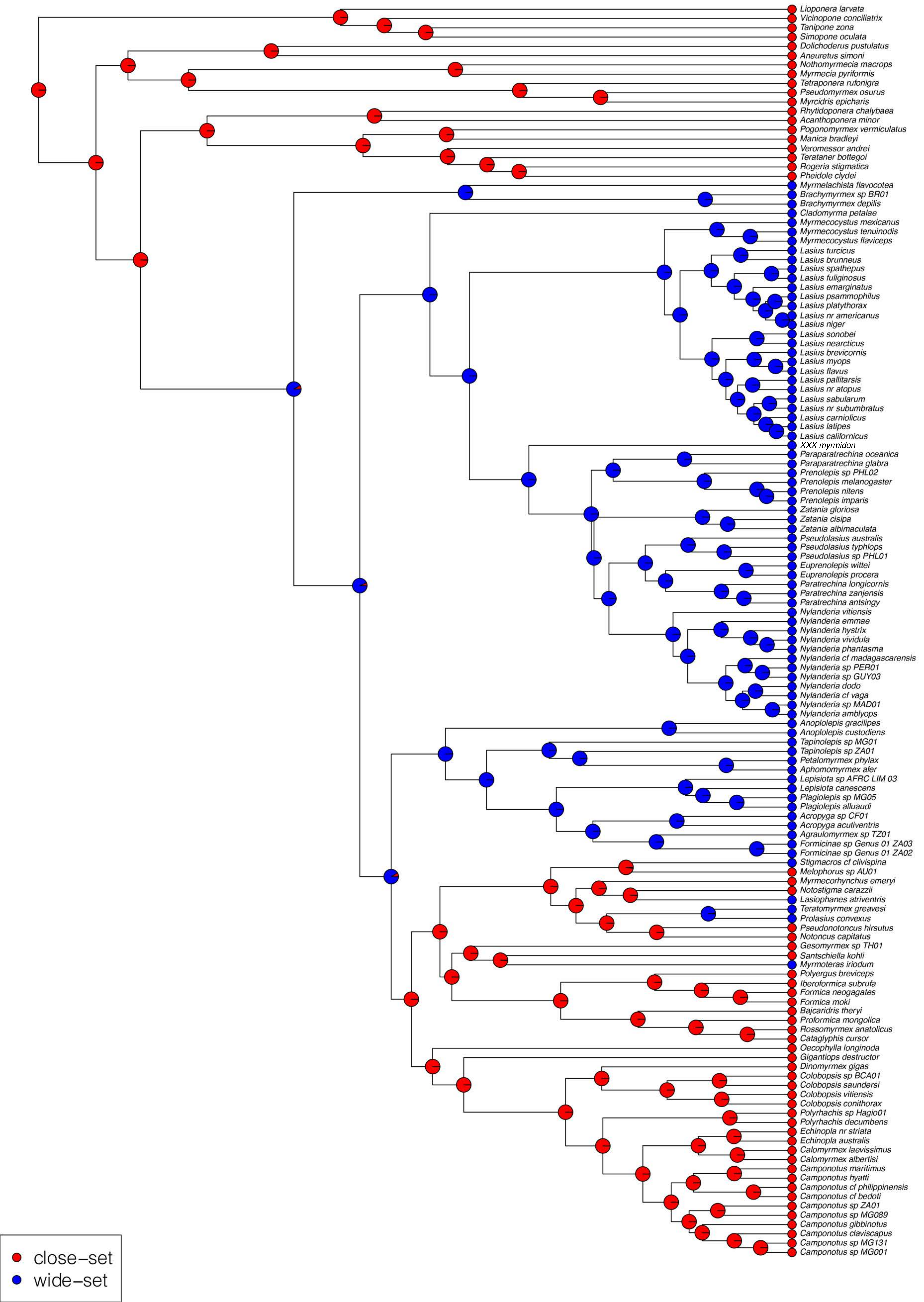

### Fig. S12

#### Lasiini & outgroups 135t Sulci ER ASR

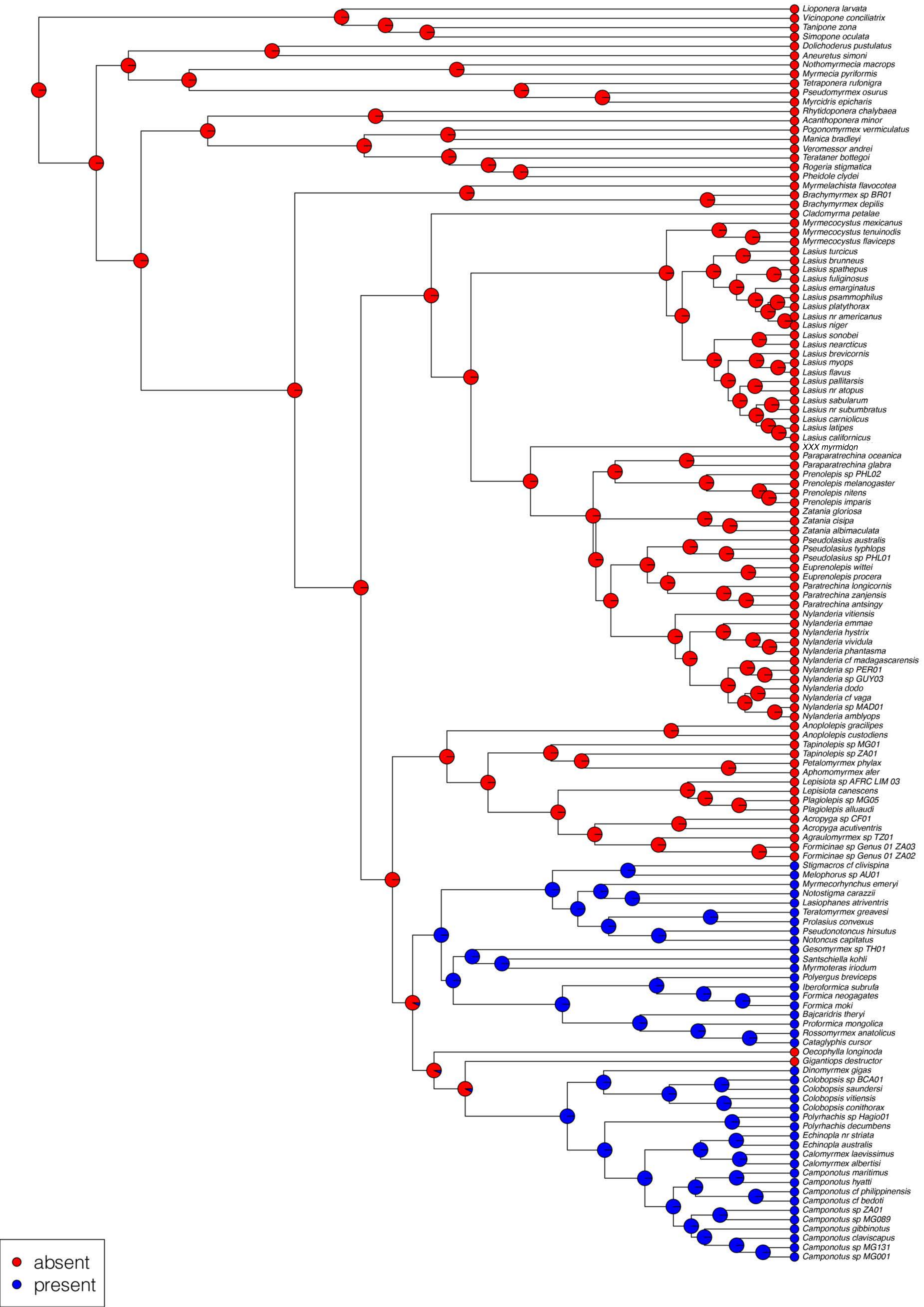

### Fig. S13

#### Lasiini & outgroups 135t Spiracles ER ASR

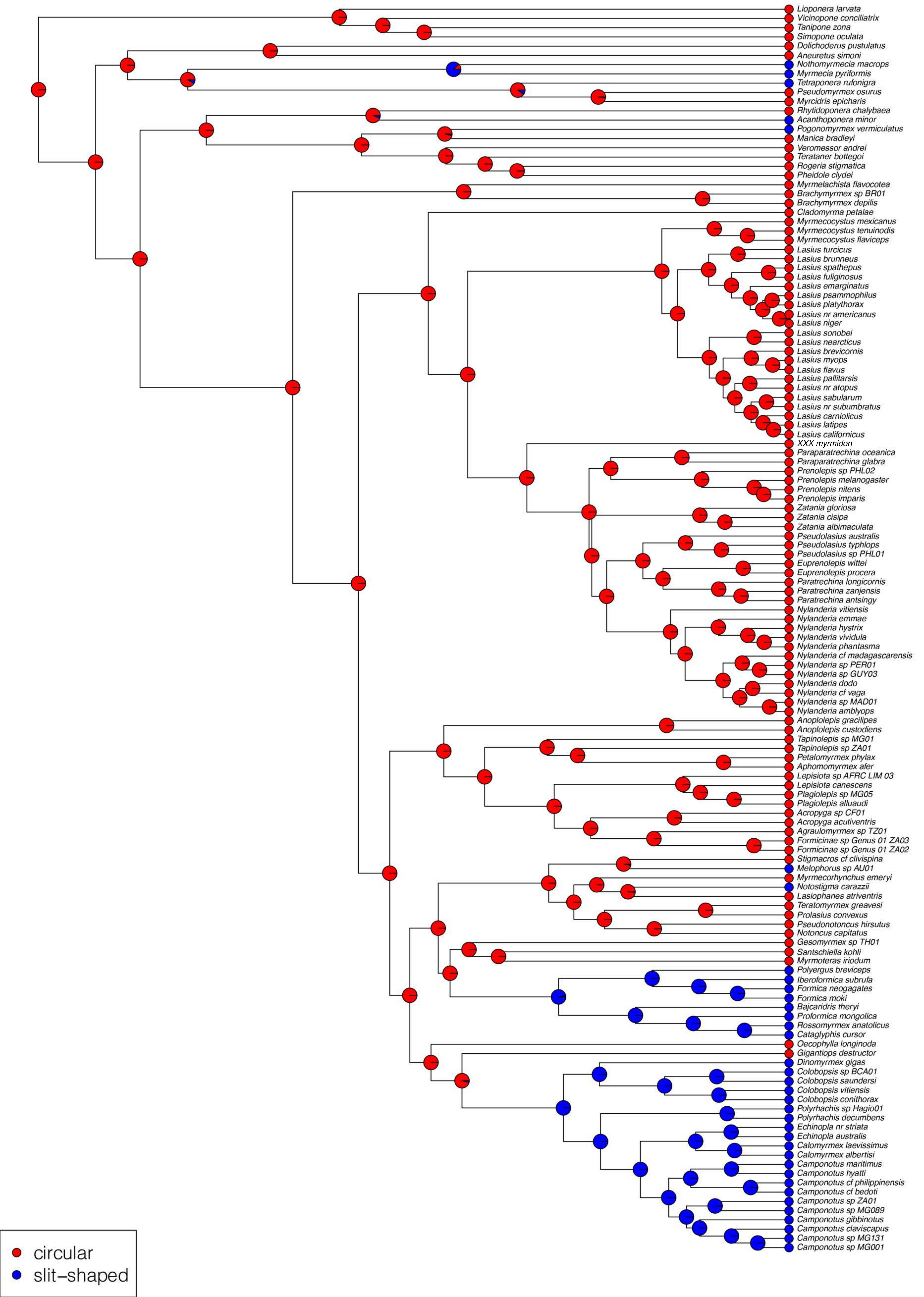

### Fig. S14

#### Lasiini & outgroups 135t Proventriculi ER ASR

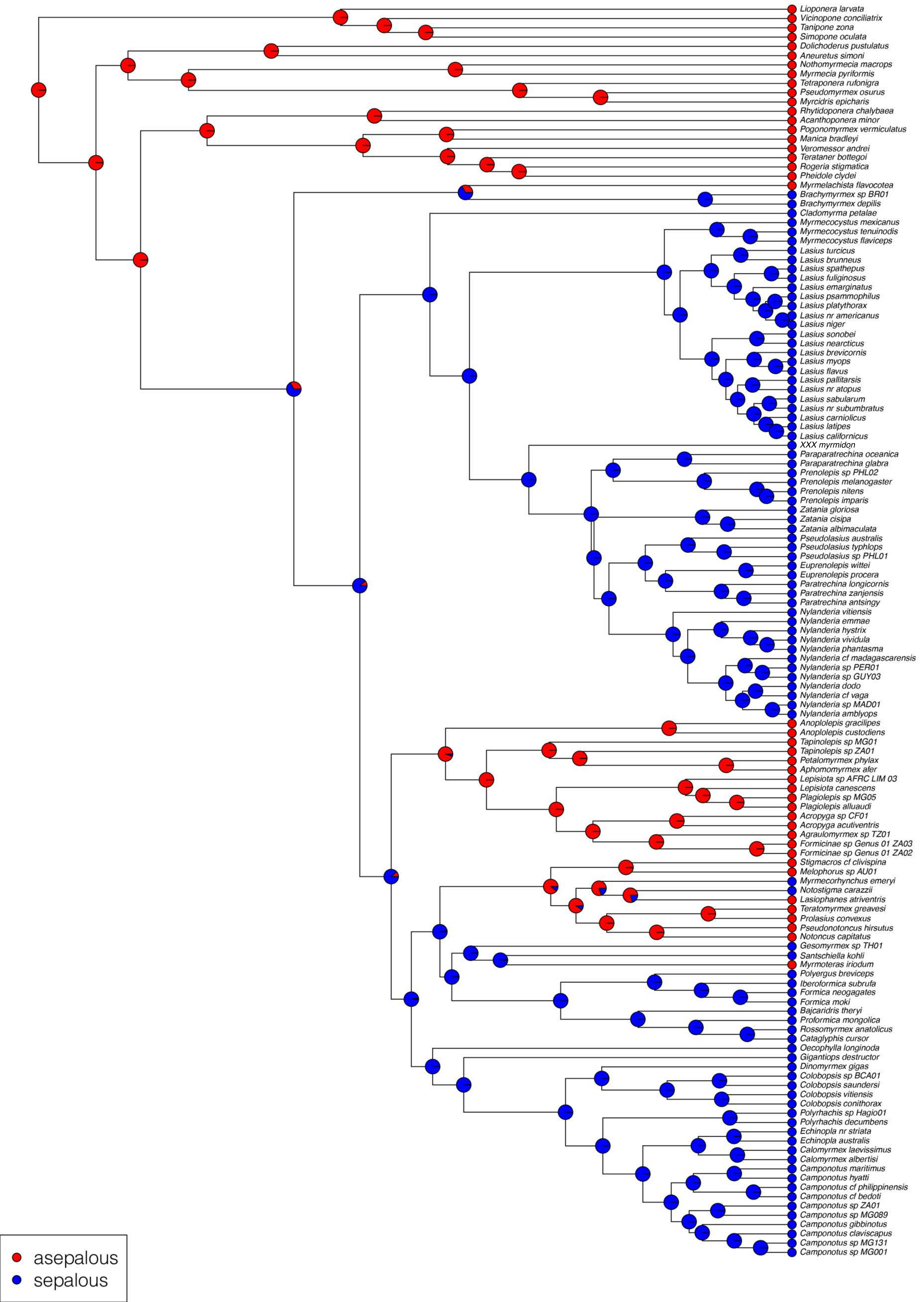

### Fig. S15

#### Lasiini & outgroups 135t AIII ER ASR

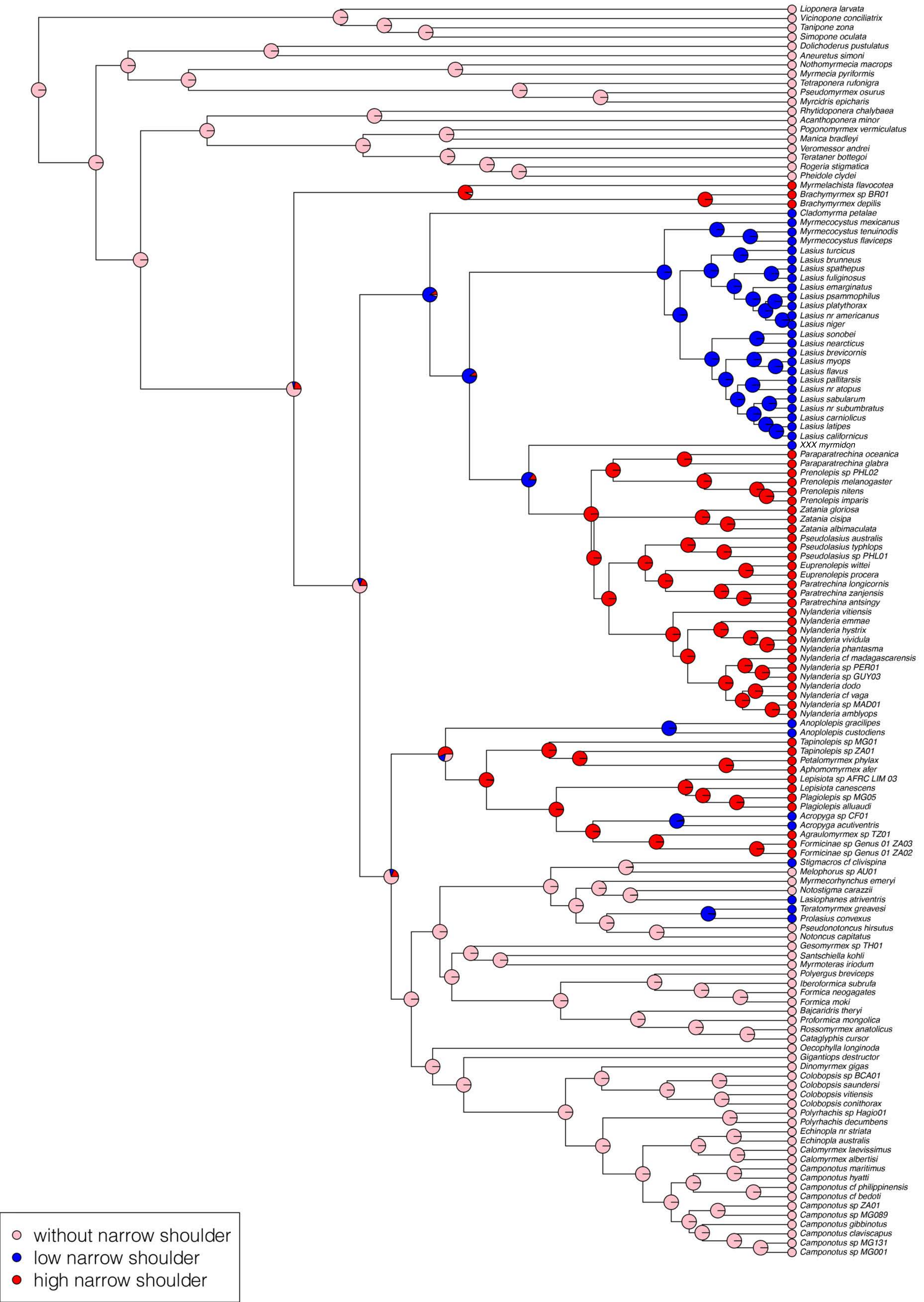

### Fig. S16

#### Lasiini 21t Parasitism ER ASR

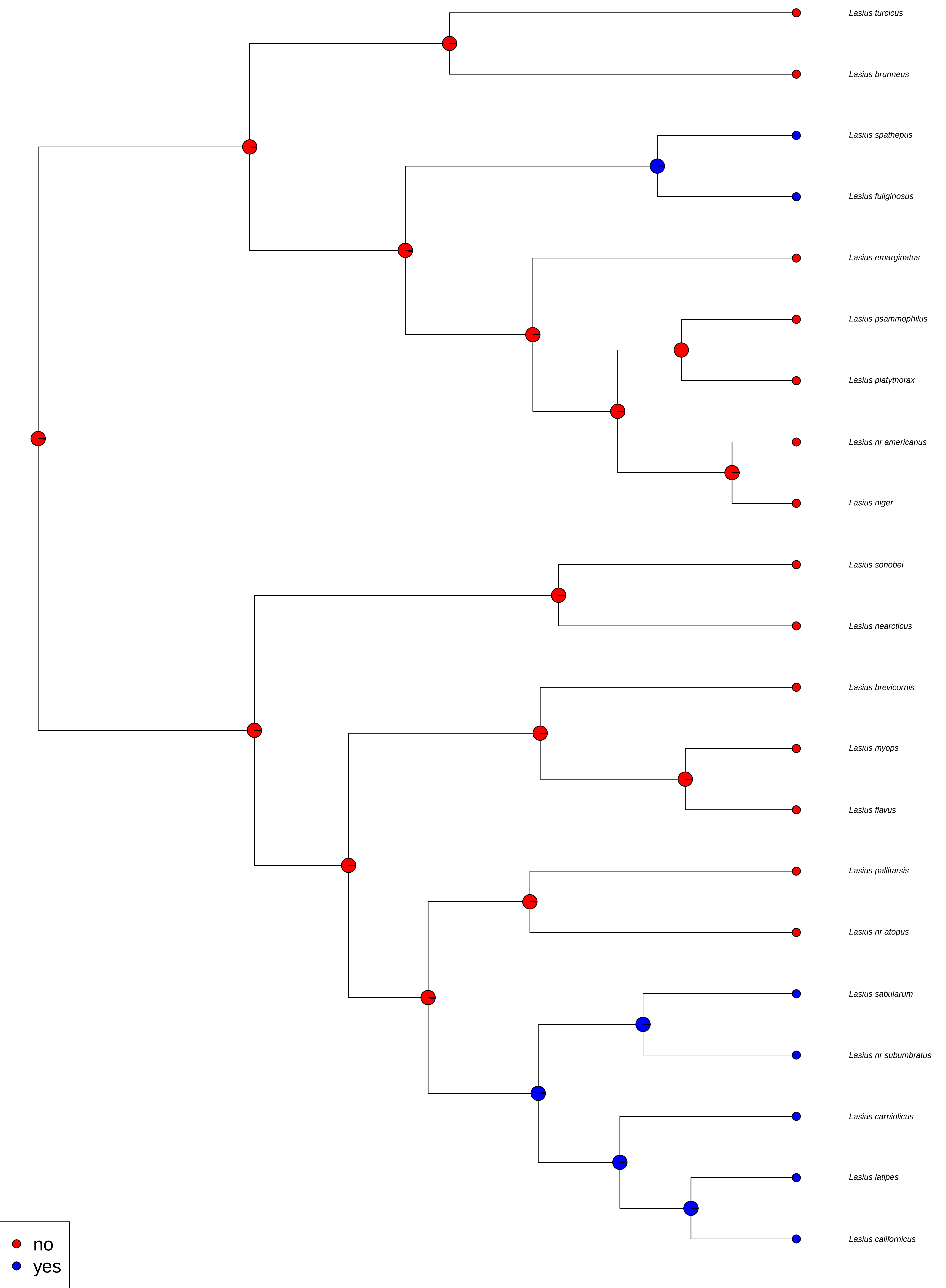

### Fig. S17

#### Lasiini 21t Fungiculture ER ASR

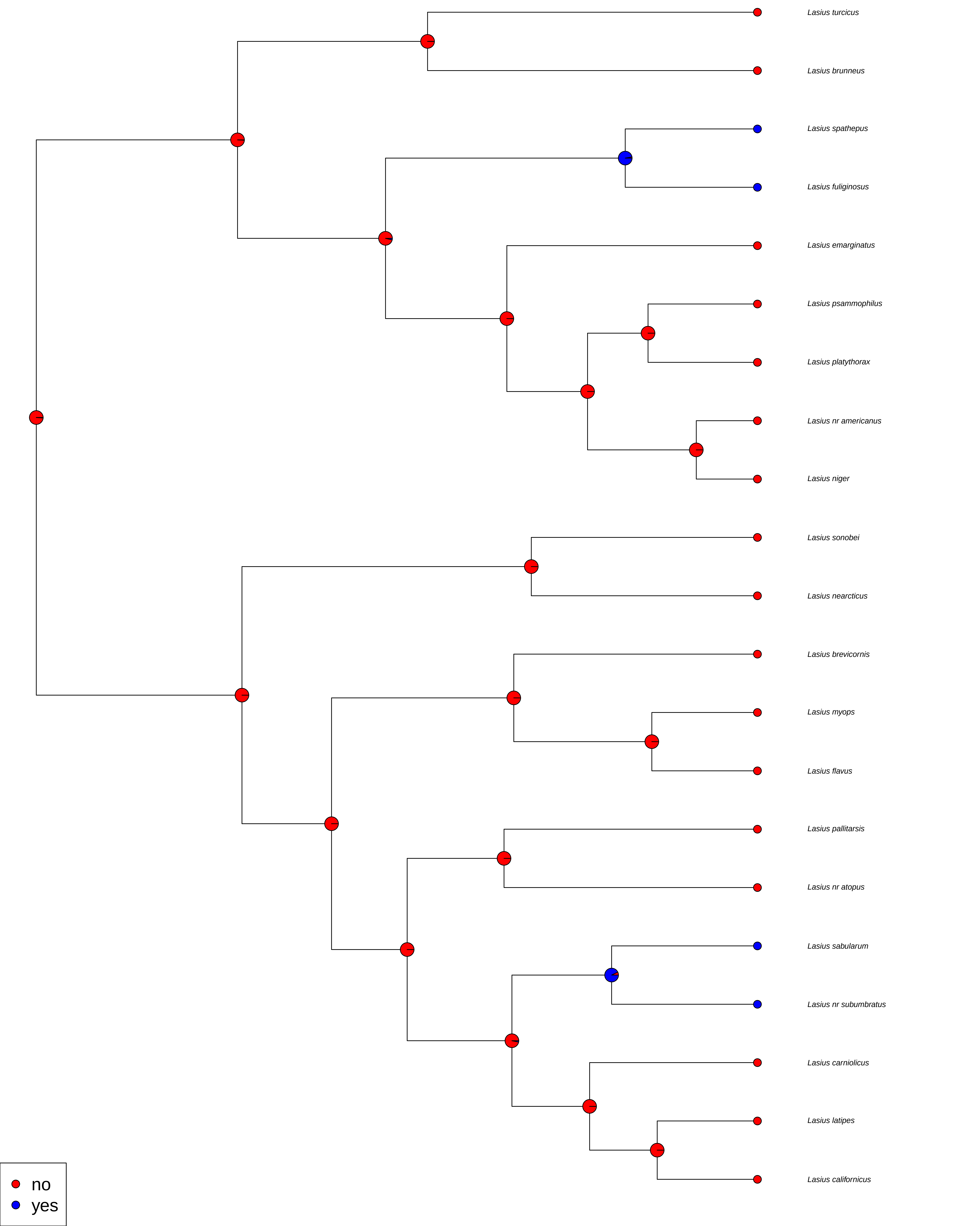

### Fig. S18

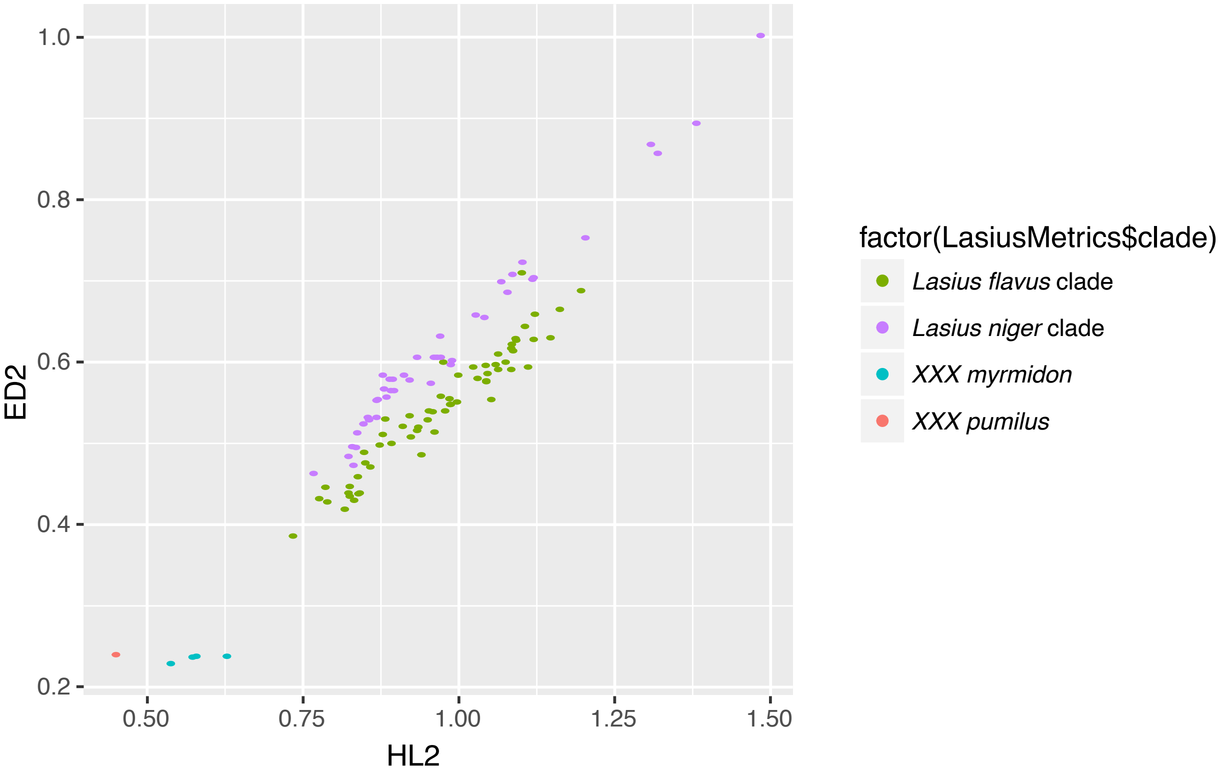

### Fig. S19

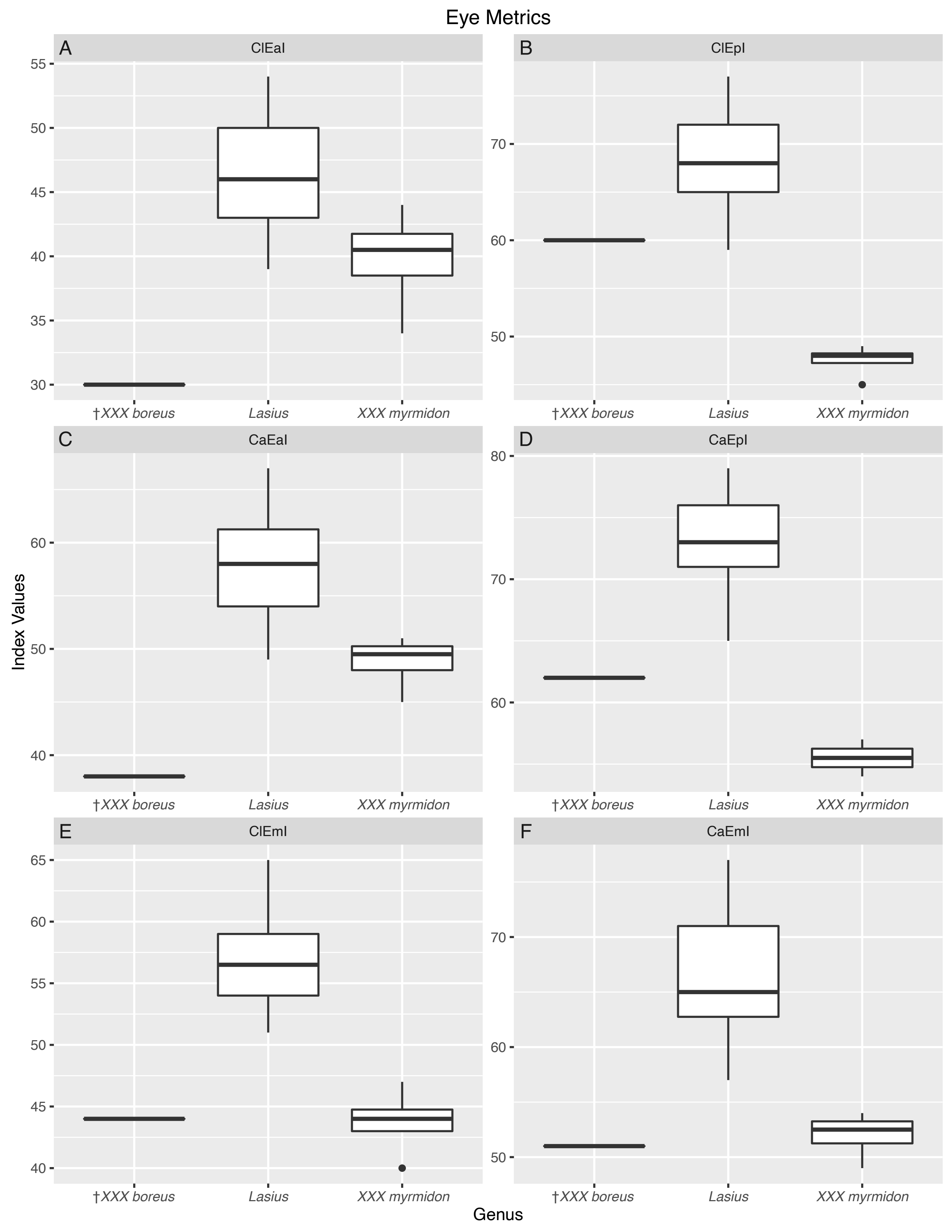

### Fig. S20

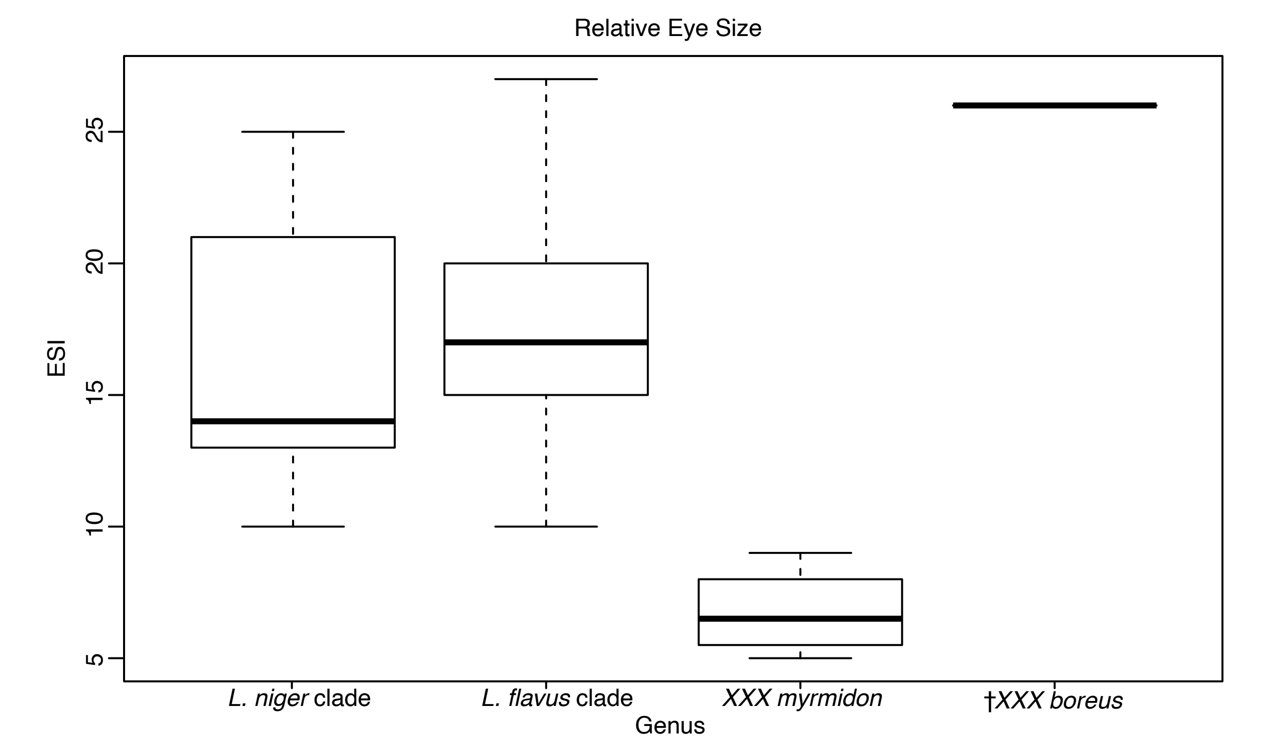
