## Supplementary material for "Phylogeny, evolution, and classification of the ant genus *Lasius*, the tribe Lasiini, and the subfamily Formicinae (Hymenoptera: Formicidae)": Table S1

| Gene | Primer | Sequence (5' to 3') | Reference |
|---|---|---|---|
| *abdA* | AA1182F* | CCGG CGAT ATG AGT ACG AAA TTC | Ward & Downie (2005) |
| | AA1881R* | GG TTG TTG GCA GGA TGT CAA AGG | Ward & Downie (2005) |
| *ArgK* | AK1F7* | ATG GTR GAY GCA GCD GTT YTG GAY AA | Branstetter (2012) |
| | AK244F | GAY CCC ATY ATY GAC GAY TAY CA | Ward et al. (2010) |
| | AK346EF* | AG GGT GAR TAC ATC GTR TCH ACT CG | Ward et al. (2010) |
| | AK345ER | ACTYAC VGT VGG RTC RAG RTT | Ward et al. (2010) |
| | AK392R* | TC CAA RGA GCG RCC GCA TC | Ward et al. (2010) |
| | AK461R | GT GCT RGA YAC YTT CTC YTC CAT | Ward et al. (2010) |
| | AK720ER* | AC CTG YCC RAG RTC ACC RCC CAT | Ward et al. (2010) |
| *CAD* | CD892F* | GGY ACC GGR CGT TGY TAY ATG AC | Ward et al. (2010) |
| | CD1028F | CG TAC TTY TCC GTB CAR TTY CAY CCR G | Ward (pers. comm.) |
| | CD1090F | TTC GAY GTG TTY YTG GAR AGY GT | Ward (pers. comm.) |
| | CD1258F4 | CAG GCY GGW GAR TTY GAY TAY TCD GGY TC | Ward (pers. comm.) |
| | CD1351F | ACG GTR CAG ACV TCV AAR GGH ATG GC | Ward et al. (2010) |
| | CD1423EF* | AG GTR ATA CRA TCG GAR AGR CCD GA | Ward et al. (2010) |
| | CD1540F | CTR GGW ACR CCR ATY GAR TCY ATH AT | Ward et al. (2010) |
| | CD1657EF | ACAG GCR TTR GAA GCY GCN GA | Ward et al. (2010) |
| | CD1106R | TC CAR RAA YAC RTC RAA RAG RCA YTC | Ward (pers. comm.) |
| | CD 1288R | G YGA RCC YGA RTA RTC RAA YTC KCC | Ward (pers. comm.) |
| | CD 1388R | TA YAC YTT RTC RGC CAT DCC YTT BGA | Ward et al. (2010) |
| | CD 1422ER2 | ARCTYAC CTG TTC KAC RTA YTC YG | Ward (pers. comm.) |
| | CD 1491R* | GCC GCA RTT NAG RGC RGT YTG YCC | Ward et al. (2010) |
| | CD 1656ER3 | TTAC CTC KTC YAC RGA RTA YAC RGC | Ward (pers. comm.) |
| | CD 1721R | CC DCC RAG NGA RAA YGC RGC RCG | Ward et al. (2010) |
| | CD 1910R* | CC GAG RGG RTC RAC RTT YTC CAT RTT RCA YAC | Ward et al. (2010) |
| *EF1αF2* | F2-557F* | GAA CGT GAA CGT GGT ATY ACS AT | Brady et al. (2006) |
| | F2-629F | ATY GAY GCY CCY GGA CAY AGR GA | Ward (pers. comm.) |
| | F2-882R | AC YTC YTT CTT RAT YTC CTC RAA YCG | Ward (pers. comm.) |
| | F2-1118R* | TTAC CTG AAG GGG AAG ACG RAG | Brady et al. (2006) |
| *LWRh* | LR116F | GGC GGA TTY GGY AAY CAR ACV GT | Ward (pers. comm.) |
| | LR125F* | GGY AAY CAR ACV GTR GTB GAC AAR GT | Ward (pers. comm.) |
| | LR143F | GAC AAA GTK CCA CCR GAR ATG CT | Ward & Downie (2005) |
| | LR398F | AAT TGC TAT TAY GAR ACN TGG GT | Ward & Downie (2005) |
| | LR480R | GA GCC ACA TCC RAA CAG RGA ACC | Ward & Downie (2005) |
| | LR 508R | GAA YGC RAT CAT CGT CAT YGT CCA | Ward & Downie (2005) |
| | LR 639ER* | YTTAC CG RTT CCA TCC RAA CA | Ward & Downie (2005) |
| *Top1* | TP1339F* | GAR CAY AAR GGA CCK GTR TTY GCA CC | Ward & Sumnicht (2012) |
| | TP1729F2 | GGY AAC TTY AAR ATY GAG CCD CCV GG | Ward & Sumnicht (2012) |
| | TP1901F2 | CY AAT GTY ACD TGG CTH GCR TCH TGG AC | Ward & Sumnicht (2012) |
| | TP1987F | GGH GAA AAR GAY TGG CAR AAR TAY GA | Ward & Sumnicht (2012) |
| | TP2065F | GAR GAY TGG AAR AGY AAR GAR ATG CG | Ward & Sumnicht (2012) |
| | TP2183F | GY TGY TGY TCG YTG CGV GTG GAR CA | Ward (pers. comm.) |
| | TP 1805R | CG CYT CTT YAR YTT RCC CAT YTT RGG | Ward & Sumnicht (2012) |
| | TP 2167R | G ATC YTC RTC CTT YTC RTT RCC RGC | Ward & Sumnicht (2012) |
| | TP 2192R* | GA RCA RCA RCC YAC DGT RTC HGC YTG | Ward & Sumnicht (2012) |
| | TP 2266ER2 | GTTAC C TAA RAA RTC RAA YAC RAC BAC | Ward & Sumnicht (2012) |
| | TP 2266ER3 | GTYAC C TAA RAA RTC RAA BAC RAC | Ward & Sumnicht (2012) |
| | TP 2354R | GG TGA CTT RTT YTC CAT RAA RAG YTG | Ward (pers. comm.) |
| *Ub* | UB1F* | GGRTA ATG AAC TCG TAY TTY GAR CAG | Ward & Sumnicht (2012) |
| | UB 671R* | AT RGC CAT CCA RGG RTA GAA SGT RTG | Ward & Sumnicht (2012) |
| *wg* | Wg290F* | GCW GTR ACT CAC AGY ATC GC | Branstetter (2012) |
| | Wg578F* | TGC ACN GTG AAR ACY TGC TGG ATG CG | Ward & Downie (2005) |
| | Wg716F | AGC AAY TCG GCS AGC AAY TCB GTG C | Ward (pers. comm.) |
| | Wg636R | AG ATT GTC BCC RAC CAC GCG | Ward (pers. comm.) |
| | Wg645R* | CG RTC CTT BAG RTT RTC GCC | Branstetter (2012) |
| | Wg822R | CC GGG YGG CTT RTG YTC CGG RTT | Ward (pers. comm.) |
| | Wg1032R* | AC YTC GCA GCA CCA RTG GAA | Abouheif & Wray (2002) |
