## Supplementary material for "Phylogeny, evolution, and classification of the ant genus *Lasius*, the tribe Lasiini, and the subfamily Formicinae (Hymenoptera: Formicidae)": Table S2

| count | Sequence information | | | | | | | locus | | | | | | | | | | | Alignment notes |
|---|---|---|---|---|---|---|---|---|---|---|---|---|---|---|---|---|---|---|---|
| | extrCode | specNum | Source | Note | Subfamily | Tribe | Terminal | abd-A | ArgK | CAD | EF-1a-F2 | LW Rh | Top1 | Ubx | Wg | EF-1a-F1 | 18S | 28S | |
| 1 | - | CASENT0039772 | Brady *et al.* 2006, Blaimer *et al.* 2015 | Non-formicine outgroup | Heteroponerinae | Heteroponerini | Acanthoponera_minor | EF013081 | KJ861140 | KJ860909 | EF013371 | EF013533 | KJ861751 | KJ860480 | EF013661 | EF013209 | EF012825 | EF012953 | |
| 2 | - | CASENT0106009 | Brady *et al.* 2006, Blaimer *et al.* 2015 | Formicine outgroup | Formicinae | Plagiolepidini | Acropyga_acutiventris | EF013083 | KT443544 | FJ982607 | EF013374 | EF013535 | KT443624 | KT443704 | EF013663 | EF013212 | EF012827 | EF012955 | |
| 3 | - | CASENT0008664 | Blaimer *et al.* 2015 | Formicine outgroup | Formicinae | Plagiolepidini | Acropyga_sp_CF01 | KT443428 | KT443545 | - | KT443200 | KT443486 | KT443625 | KT443705 | KT443256 | KT443144 | KT443312 | KT443370 | |
| 4 | - | CASENT0217051 | Blaimer *et al.* 2015 | Formicine outgroup | Formicinae | Plagiolepidini | Agraulomyrmex_sp_TZ01 | KT443429 | KT443546 | - | KT443201 | KT443487 | KT443626 | KT443706 | KT443257 | KT443145 | KT443313 | KT443371 | |
| 5 | - | CASENT0007014 | Brady *et al.* 2006, Ward *et al.* 2010, Blaimer *et al.* 2015 | Non-formicine outgroup | Aneuretinae | Aneuretini | Aneuretus_simoni | EF013089 | FJ939840 | FJ939894 | EF013382 | EF013541 | KJ523645 | KJ523562 | EF013669 | EF013220 | EF012833 | EF012961 | |
| 6 | - | CASENT0106259 | Blaimer *et al.* 2015 | Formicine outgroup | Formicinae | Plagiolepidini | Anoplolepis_custodiens | KT443430 | KT443547 | - | KT443202 | KT443488 | KT443627 | KT443707 | KT443258 | KT443146 | KT443314 | KT443372 | |
| 7 | - | CASENT0106057 | Brady *et al.* 2006, Blaimer *et al.* 2015 | Formicine outgroup | Formicinae | Plagiolepidini | Anoplolepis_gracilipes | EF013092 | KT443548 | FJ982606 | EF013385 | EF013544 | KT443628 | KT443708 | EF013672 | EF013223 | EF012836 | EF012964 | |
| 8 | - | CASENT0217049 | Blaimer *et al.* 2015 | Formicine outgroup | Formicinae | Plagiolepidini | Aphomomyrmex_afer | KT443431 | KT443549 | - | KT443203 | KT443489 | KT443629 | KT443709 | KT443259 | KT443147 | KT443315 | KT443373 | |
| 9 | - | CASENT0235255 | Blaimer *et al.* 2015 | Formicine outgroup | Formicinae | Formicini | Bajcaridris_theryi | KT443432 | KT443550 | - | KT443204 | KT443490 | KT443630 | KT443710 | KT443260 | KT443148 | KT443316 | KT443374 | |
| 10 | - | CASENT0106038 | Brady *et al.* 2006, Blaimer *et al.* 2015 | Formicine outgroup | Formicinae | Myrmelachistini | Brachymyrmex_depilis | EF013100 | KT443552 | - | EF013395 | EF013552 | KT443632 | KT443712 | EF013680 | EF013233 | EF012844 | EF012972 | |
| 11 | - | CASENT0217326 | Blaimer *et al.* 2015 | Formicine outgroup | Formicinae | Myrmelachistini | Brachymyrmex_sp_BR01 | KT443433 | KT443551 | - | KT443205 | KT443491 | KT443631 | KT443711 | KT443261 | KT443149 | KT443317 | KT443375 | |
| 12 | - | CASENT0106006 | Brady *et al.* 2006, Blaimer *et al.* 2015 | Formicine outgroup | Formicinae | Camponotini | Calomyrmex_albertisi | EF013101 | KT443553 | - | EF013396 | EF013553 | KT443633 | KT443713 | EF013681 | EF013234 | EF012845 | EF012973 | |
| 13 | - | CASENT0106150 | Blaimer *et al.* 2015 | Formicine outgroup | Formicinae | Camponotini | Calomyrmex_laevissimus | KT443434 | KT443554 | - | KT443206 | KT443492 | KT443634 | KT443714 | KT443262 | KT443150 | KT443318 | KT443376 | |
| 14 | - | CASENT0106261 | Blaimer *et al.* 2015 | Formicine outgroup | Formicinae | Camponotini | Camponotus_cf_bedoti | KT443435 | KT443556 | - | KT443207 | KT443493 | KT443636 | KT443716 | KT443263 | KT443151 | KT443319 | KT443377 | |
| 15 | - | CASENT0106324 | Blaimer *et al.* 2015 | Formicine outgroup | Formicinae | Camponotini | Camponotus_cf_philippinensis | KT443449 | KT443573 | - | KT443221 | KT443507 | KT443653 | KT443733 | KT443277 | KT443165 | KT443333 | KT443391 | |
| 16 | - | CASENT0106332 | Blaimer *et al.* 2015 | Formicine outgroup | Formicinae | Camponotini | Camponotus_claviscapus | KT443436 | KT443557 | - | KT443208 | KT443494 | KT443637 | KT443717 | KT443264 | KT443152 | KT443320 | KT443378 | |
| 17 | - | CASENT0106239 | Blaimer *et al.* 2015 | Formicine outgroup | Formicinae | Camponotini | Camponotus_gibbinotus | KT443437 | KT443559 | - | KT443209 | KT443495 | KT443639 | KT443719 | KT443265 | KT443153 | KT443321 | KT443379 | |
| 18 | - | CASENT0106032 | Brady *et al.* 2006, Blaimer *et al.* 2015 | Formicine outgroup | Formicinae | Camponotini | Camponotus_hyatti | EF013104 | KT443561 | - | EF013399 | EF013556 | KT443641 | KT443721 | EF013684 | EF013237 | EF012848 | EF012976 | |
| 19 | - | CASENT0106083 | Ward 2005, Brady *et al.* 2006, Blaimer *et al.* 2015 | Formicine outgroup | Formicinae | Camponotini | Camponotus_maritimus | AY867480 | KT443562 | - | EF013400 | AY867495 | KT443642 | KT443722 | AY867433 | EF013238 | AY867448 | AY867464 | |
| 20 | - | CASENT0499291 | Blaimer *et al.* 2015 | Formicine outgroup | Formicinae | Camponotini | Camponotus_sp_MG001 | KT443439 | KT443563 | - | KT443211 | KT443497 | KT443643 | KT443723 | KT443267 | KT443155 | KT443323 | KT443381 | |
| 21 | - | CASENT0121990 | Blaimer *et al.* 2015 | Formicine outgroup | Formicinae | Camponotini | Camponotus_sp_MG089 | KT443440 | KT443564 | - | KT443212 | KT443498 | KT443644 | KT443724 | KT443268 | KT443156 | KT443324 | KT443382 | |
| 22 | - | CASENT0159208 | Blaimer *et al.* 2015 | Formicine outgroup | Formicinae | Camponotini | Camponotus_sp_MG131 | KT443441 | KT443565 | - | KT443213 | KT443499 | KT443645 | KT443725 | KT443269 | KT443157 | KT443325 | KT443383 | |
| 23 | - | CASENT0217047 | Blaimer *et al.* 2015 | Formicine outgroup | Formicinae | Camponotini | Camponotus_sp_ZA01 | KT443471 | KT443603 | - | KT443241 | KT443529 | KT443683 | KT443763 | KT443297 | KT443185 | KT443355 | KT443413 | |
| 24 | - | CASENT0106260 | Blaimer *et al.* 2015 | Formicine outgroup | Formicinae | Formicini | Cataglyphis_cursor | KT443444 | KT443568 | - | KT443216 | KT443502 | KT443648 | KT443728 | KT443272 | KT443160 | KT443328 | KT443386 | |
| 25 | - | CASENT0235222 | Blaimer *et al.* 2015 | Ingroup | Formicinae | Lasiini | Cladomyrma_petalae | KT443445 | KT443569 | - | KT443217 | KT443503 | KT443649 | KT443729 | KT443273 | KT443161 | KT443329 | KT443387 | |
| 26 | - | CASENT0106008 | Brady *et al.* 2006, Blaimer *et al.* 2015 | Formicine outgroup | Formicinae | Camponotini | Colobopsis_conithorax | EF013103 | KT443558 | - | EF013398 | EF013555 | KT443638 | KT443718 | EF013683 | EF013236 | EF012847 | EF012975 | |
| 27 | - | CASENT0106127 | Blaimer *et al.* 2015 | Formicine outgroup | Formicinae | Camponotini | Colobopsis_saundersi | KT443442 | KT443566 | - | KT443214 | KT443500 | KT443646 | KT443726 | KT443270 | KT443158 | KT443326 | KT443384 | |
| 28 | - | CASENT0106050 | Brady *et al.* 2006, Blaimer *et al.* 2015 | Formicine outgroup | Formicinae | Camponotini | Colobopsis_sp_BCA01 | EF013102 | KT443555 | - | EF013397 | EF013554 | KT443635 | KT443715 | EF013682 | EF013235 | EF012846 | EF012974 | |
| 29 | - | CASENT0106151 | Blaimer *et al.* 2015 | Formicine outgroup | Formicinae | Camponotini | Colobopsis_vitiensis | KT443443 | KT443567 | - | KT443215 | KT443501 | KT443647 | KT443727 | KT443271 | KT443159 | KT443327 | KT443385 | |
| 30 | - | CASENT0106240 | Blaimer *et al.* 2015 | Formicine outgroup | Formicinae | Camponotini | Dinomyrmex_gigas | KT443438 | KT443560 | - | KT443210 | KT443496 | KT443640 | KT443720 | KT443266 | KT443154 | KT443322 | KT443380 | |
| 31 | - | CASENT0106164 | Ward *et al.* 2010, Blaimer *et al.* 2015 | Non-formicine outgroup | Dolichoderinae | Dolichderini | Dolichoderus_pustulatus | FJ939824 | FJ939859 | FJ939913 | FJ939963 | FJ939995 | KJ523685 | KJ523602 | FJ940028 | FJ940060 | FJ939760 | FJ939792 | |
| 32 | - | CASENT0106149 | Blaimer *et al.* 2015 | Formicine outgroup | Formicinae | Camponotini | Echinopla_australis | KT443446 | KT443570 | - | KT443218 | KT443504 | KT443650 | KT443730 | KT443274 | KT443162 | KT443330 | KT443388 | |
| 33 | - | CASENT0106129 | Blaimer *et al.* 2015 | Formicine outgroup | Formicinae | Camponotini | Echinopla_nr_striata | KT443447 | KT443571 | - | KT443219 | KT443505 | KT443651 | KT443731 | KT443275 | KT443163 | KT443331 | KT443389 | |
| 34 | - | CASENT0106328 | Blaimer *et al.* 2015 | Ingroup | Formicinae | Lasiini | Euprenolepis_procera | KT443448 | KT443572 | - | KT443220 | KT443506 | KT443652 | KT443732 | KT443276 | KT443164 | KT443332 | KT443390 | |
| 35 | ParaM39 | USNM00756212 | LaPolla et al. 2010, LaPolla *et al*. 2012 | Ingroup | Formicinae | Lasiini | Euprenolepis_wittei | - | JN562403 | FJ982609 | FJ982566 | - | - | - | JN562470 | FJ982486 | - | - | |
| 35 | - | CASENT0106084 | Ward & Downie 2005, Brady *et al.* 2006, Blaimer *et al.* 2015 | Formicine outgroup | Formicinae | Formicini | Formica_moki | AY703694 | KT443574 | - | EF013425 | AY703761 | KT443654 | KT443734 | AY703627 | EF013263 | AY703493 | AY703560 | |
| 36 | - | CASENT0106302 | Blaimer *et al.* 2015 | Formicine outgroup | Formicinae | Formicini | Formica_neogagates | KT443450 | KT443575 | - | KT443222 | KT443508 | KT443655 | KT443735 | KT443278 | KT443166 | KT443334 | KT443392 | |
| 37 | - | CASENT0217131 | Blaimer *et al.* 2015 | Formicine outgroup | Formicinae | Plagiolepidini | Formicinae_genus_01_ZA02 | KT443483 | KT443620 | - | KT443253 | KT443541 | KT443700 | KT443780 | KT443309 | KT443197 | KT443367 | KT443425 | |
| 38 | - | CASENT0217132 | Blaimer *et al.* 2015 | Formicine outgroup | Formicinae | Plagiolepidini | Formicinae_genus_01_ZA03 | KT443484 | KT443621 | - | KT443254 | KT443542 | KT443701 | KT443781 | KT443310 | KT443198 | KT443368 | KT443426 | |
| 39 | | CASENT0118772 | Blaimer *et al.* 2015 | Formicine outgroup | Formicinae | Gesomyrmecini | Gesomyrmex_sp_TH01 | KT443452 | KT443577 | - | KT443224 | KT443510 | KT443657 | KT443737 | KT443280 | KT443168 | KT443336 | KT443394 | |
| 40 | | CASENT0106169 | Blaimer *et al.* 2015 | Formicine outgroup | Formicinae | Gigantiopini | Gigantiops_destructor | KT443453 | KT443578 | - | KT443225 | KT443511 | KT443658 | KT443738 | KT443281 | KT443169 | KT443337 | KT443395 | |
| 41 | - | CASENT0270631 | Blaimer *et al.* 2015 | Formicine outgroup | Formicinae | Formicini | Iberoformica_subrufa | KT443454 | KT443579 | - | KT443226 | KT443512 | KT443659 | KT443739 | KT443282 | KT443170 | KT443338 | KT443396 | |
| 42 | - | CASENT0106235 | Blaimer *et al.* 2015 | Formicine outgroup | Formicinae | Melophorini | Lasiophanes_atriventris | KT443455 | KT443580 | - | KT443227 | KT443513 | KT443660 | KT443740 | KT443283 | KT443171 | KT443339 | KT443397 | |
| 42 | M162 | CASENT0234864 | this study | Ingroup | Formicinae | Lasiini | Lasius_brevicornis | MZ556635 | MZ556655 | MZ556675 | MZ556715 | MZ556718 | MZ556738 | MZ556760 | MZ556782 | - | - | - | 15 bp at 5' end of AA not included |
| 43 | M41 | CASENT0234862 | this study | Ingroup | Formicinae | Lasiini | Lasius_brunneus | MZ556636 | MZ556656 | MZ556676 | MZ556698 | MZ556719 | MZ556739 | MZ556761 | MZ556783 | - | - | - | 15 bp at 5' end of AA not included |
| 49 | - | CASENT0106045 | Brady *et al.* 2006, LaPolla *et al.* 2010, Brady *et al.* 2014 | Ingroup | Formicinae | Lasiini | Lasius_californicus | EF013126 | KJ523346 | FJ982605 | EF013430 | EF013578 | KJ523692 | KJ523609 | EF013706 | EF013268 | EF012870 | EF012998 | |
| 44 | M49 | CASENT0731085 | this study | Ingroup | Formicinae | Lasiini | Lasius_carniolicus | MZ556637 | MZ556657 | MZ556677 | MZ556699 | MZ556720 | MZ556740 | MZ556762 | MZ556784 | - | - | - | 15 bp at 5' end of AA not included |
| 43 | M44 | CASENT0234865 | this study | Ingroup | Formicinae | Lasiini | Lasius_emarginatus | MZ556638 | MZ556658 | MZ556678 | MZ556700 | MZ556721 | MZ556741 | MZ556763 | MZ556785 | - | - | - | 15 bp at 5' end of AA not included |
| 46 | M39 | CASENT0234860 | this study | Ingroup | Formicinae | Lasiini | Lasius_flavus | MZ556639 | MZ556659 | MZ556679 | MZ556701 | MZ556722 | MZ556742 | MZ556764 | MZ556786 | - | - | - | 15 bp at 5' end of AA not included |
| 47 | M40 | CASENT0234861 | this study | Ingroup | Formicinae | Lasiini | Lasius_fuliginosus | MZ556640 | MZ556660 | MZ556680 | MZ556702 | MZ556723 | MZ556743 | MZ556765 | MZ556787 | - | - | - | 15 bp at 5' end of AA not included |
| 48 | M45 | CASENT0234866 | this study | Ingroup | Formicinae | Lasiini | Lasius_latipes | MZ556641 | MZ556661 | MZ556681 | MZ556703 | MZ556724 | MZ556744 | MZ556766 | MZ556788 | - | - | - | 15 bp at 5' end of AA not included |
| 50 | M164 | CASENT0732040 | this study | Ingroup | Formicinae | Lasiini | Lasius_myops | MZ556642 | MZ556662 | MZ556682 | MZ556716 | MZ556725 | MZ556745 | MZ556767 | MZ556789 | - | - | - | 15 bp at 5' end of AA not included |
| 51 | M119 | CASENT0732003 | this study | Ingroup | Formicinae | Lasiini | Lasius_nearcticus | MZ556643 | MZ556663 | MZ556683 | MZ556704 | MZ556726 | MZ556746 | MZ556768 | MZ556790 | - | - | - | 15 bp at 5' end of AA not included |
| 52 | - | CASENT0106128 | Blaimer *et al.* 2015 | Ingroup | Formicinae | Lasiini | Lasius_niger | KT443456 | KT443581 | MZ556684 | KT443228 | KT443514 | KT443661 | KT443741 | KT443284 | KT443172 | KT443340 | KT443398 | |
| 53 | M117 | CASENT0732004 | this study | Ingroup | Formicinae | Lasiini | Lasius_nr_americanus | MZ556644 | MZ556664 | MZ556685 | MZ556705 | MZ556727 | MZ556747 | MZ556769 | MZ556791 | - | - | - | 15 bp at 5' end of AA not included |
| 44 | M37 | CASENT0234858 | this study | Ingroup | Formicinae | Lasiini | Lasius_nr_atopus | MZ556645 | MZ556665 | MZ556686 | MZ556706 | MZ556728 | MZ556748 | MZ556770 | MZ556792 | - | - | - | 15 bp at 5' end of AA not included |
| 55 | M48 | CASENT0731084 | this study | Ingroup | Formicinae | Lasiini | Lasius_nr_subumbratus | MZ556646 | MZ556666 | MZ556687 | MZ556707 | MZ556729 | MZ556749 | MZ556771 | MZ556793 | - | - | - | 15 bp at 5' end of AA not included |
| 56 | M38 | CASENT0234859 | this study | Ingroup | Formicinae | Lasiini | Lasius_pallitarsis | MZ556647 | MZ556667 | MZ556688 | MZ556708 | MZ556730 | MZ556750 | MZ556772 | MZ556794 | - | - | - | 15 bp at 5' end of AA not included |
| 57 | M120 | CASENT0732002 | this study | Ingroup | Formicinae | Lasiini | Lasius_platythorax | MZ556648 | MZ556668 | MZ556689 | MZ556709 | MZ556731 | MZ556751 | MZ556773 | MZ556795 | - | - | - | 15 bp at 5' end of AA not included |
| 58 | M121 | CASENT0732000 | this study | Ingroup | Formicinae | Lasiini | Lasius_psammophilus | MZ556649 | MZ556669 | MZ556690 | MZ556710 | MZ556732 | MZ556752 | MZ556774 | MZ556796 | - | - | - | 15 bp at 5' end of AA not included |
| 59 | M42 | CASENT0234863 | this study | Ingroup | Formicinae | Lasiini | Lasius_sabularum | MZ556650 | MZ556670 | MZ556691 | MZ556711 | MZ556733 | MZ556753 | MZ556775 | MZ556797 | - | - | - | 15 bp at 5' end of AA not included |
| 60 | M124 | CASENT0730575 | this study | Ingroup | Formicinae | Lasiini | Lasius_sonobei | MZ556651 | MZ556671 | MZ556692 | MZ556712 | MZ556734 | MZ556754 | MZ556776 | MZ556798 | - | - | - | 15 bp at 5' end of AA not included |
| 61 | M122 | CASENT0732006 | this study | Ingroup | Formicinae | Lasiini | Lasius_spathepus | MZ556652 | MZ556672 | MZ556693 | MZ556713 | MZ556735 | MZ556755 | MZ556777 | MZ556799 | - | - | - | 15 bp at 5' end of AA not included |
| 62 | M118 | CASENT0732001 | this study | Ingroup | Formicinae | Lasiini | Lasius_turcicus | MZ556653 | MZ556673 | MZ556694 | MZ556714 | MZ556736 | MZ556756 | MZ556778 | MZ556800 | - | - | - | 15 bp at 5' end of AA not included |
| 45 | - | CASENT0138305 | Blaimer *et al.* 2015 | Formicine outgroup | Formicinae | Plagiolepidini | Lepisiota_canescens | KT443458 | KT443583 | - | KT443230 | KT443516 | KT443663 | KT443743 | KT443286 | KT443174 | KT443342 | KT443399 | |
| 46 | - | CASENT0280596 | Blaimer *et al.* 2015 | Formicine outgroup | Formicinae | Plagiolepidini | Lepisiota_sp_AFRC_LIM_03 | KT443457 | KT443582 | - | KT443229 | KT443515 | KT443662 | KT443742 | KT443285 | KT443173 | KT443341 | KT443400 | |
| 120 | | CASENT0106073 | Ward & Downie 2005, Brady et al. 2006, Brady et al. 2014 | Non-formicine outgroup | Dorylinae | | Lioponera_larvata | AY703692 | KJ523316 | KJ523395 | EF013405 | AY703759 | KJ523661 | KJ523578 | AY233633 | EF013243 | AY703491 | AY703558 | |
| 47 | - | CASENT0106022 | Brady *et al.* 2006, Ward *et al.* 2010, Brady *et al.* 2014 | Non-formicine outgroup | Myrmicinae | Myrmicini | Manica_bradleyi | EF013134 | FJ939877 | FJ939931 | EF013443 | EF013586 | KJ523698 | KJ523615 | EF013714 | EF013281 | EF012878 | EF013006 | |
| 48 | - | CASENT0106148 | Blaimer *et al.* 2015 | Formicine outgroup | Formicinae | Melophorini | Melophorus_sp_AU01 | KT443459 | KT443584 | - | KT443231 | KT443517 | KT443664 | KT443744 | KT443287 | KT443175 | KT443343 | KT443401 | |
| 67 | M163 | CASENT0811544 | this study | Ingroup | Formicinae | Lasiini | XXX_myrmidon | MZ556654 | MZ556674 | MZ556695 | MZ556717 | MZ556737 | MZ556757 | MZ556779 | MZ556801 | - | - | - | 15 bp at 5' end of AA not included |
| 124 | | CASENT0106096 | Ward & Downie 2005, Brady *et al.* 2006, Chomicki *et al*. 2015 | Non-formicine outgroup | Pseudomyrmecinae | | Myrcidris_epicharis | AY703718 | KR829767 | KR830182 | EF013453 | AY703785 | KR830598 | KR830389 | AY703651 | EF013291 | AY703517 | AY703584 | |
| 49 | - | CASENT0106088 | Ward & Downie 2005, Brady *et al.* 2006, Ward *et al.* 2010, Brady *et al.* 2014 | Non-formicine outgroup | Myrmeciinae | Myrmeciini | Myrmecia_pyriformis | AY703701 | FJ939878 | FJ939932 | EF013454 | AY703768 | KJ523699 | KJ523616 | AY703634 | EF013292 | AY703500 | AY703567 | |
| 50 | - | CASENT0106055 | Brady *et al.* 2006, Blaimer *et al.* 2015 | Ingroup | Formicinae | Lasiini | Myrmecocystus_flaviceps | EF013144 | KT443585 | FJ982608 | EF013456 | EF013596 | KT443665 | KT443745 | EF013724 | EF013294 | EF012888 | EF013016 | |
| 70 | M46 | "BCO172" | Moreau *et al.* 2006, this study, O'Meara (unpubl.) | Ingroup | Formicinae | Lasiini | Myrmecocystus_mexicanus | EU142976 | EU143007 | MZ556696 | EU143096 | EU143151 | MZ556758 | MZ556780 | EU143233 | EU143096 | - | - | 15 bp at 5' end of AA not included |
| 71 | M47 | "PSW15178" | this study, O'Meara (unpubl.) | Ingroup | Formicinae | Lasiini | Myrmecocystus_tenuinodis | EU142956 | EU142986 | MZ556697 | EU143076 | EU143131 | MZ556759 | MZ556781 | EU143216 | EU143076 | - | - | 15 bp at 5' end of AA not included |
| 51 | - | CASENT0227590 | Blaimer *et al.* 2015 | Formicine outgroup | Formicinae | Melophorini | Myrmecorhynchus_emeryi | KT443460 | KT443586 | - | KT443232 | KT443518 | KT443666 | KT443746 | KT443288 | KT443176 | KT443344 | KT443402 | |
| 52 | - | CASENT0106049 | Brady *et al.* 2006, Ward *et al.* 2015 | Formicine outgroup | Formicinae | Myrmelachistini | Myrmelachista_flavocotea | EF013145 | KJ861144 | KJ860913 | EF013457 | EF013597 | KJ861755 | KJ860484 | EF013725 | EF013295 | EF012889 | EF013017 | |
| 53 | | CASENT0006837 | Brady *et al.* 2006, Blaimer *et al.* 2015 | Formicine outgroup | Formicinae | Myrmoteratini | Myrmoteras_iriodum | EF013149 | KT443587 | - | EF013462 | EF013601 | KT443667 | KT443747 | EF013729 | EF013300 | EF012893 | EF013021 | |
| 54 | - | CASENT0106089 | Ward & Downie 2005, Brady *et al.* 2006, Ward *et al.* 2010, Brady *et al.* 2014 | Non-formicine outgroup | Myrmeciinae | Prionomyrmecini | Nothomyrmecia_macrops | AY703702 | FJ939880 | FJ939934 | EF013466 | AY703769 | KJ523703 | KJ523620 | AY703635 | EF013304 | AY703501 | AY703568 | |
| 55 | - | CASENT0106015 | Brady *et al.* 2006, Blaimer *et al.* 2015 | Formicine outgroup | Formicinae | Melophorini | Notoncus_capitatus | EF013152 | KT443588 | - | EF013467 | EF013604 | KT443668 | KT443748 | EF013732 | EF013305 | EF012896 | EF013024 | |
| 56 | - | CASENT0106112 | Brady *et al.* 2006, Blaimer *et al.* 2015 | Formicine outgroup | Formicinae | Melophorini | Notostigma_carazzii | EF013153 | KT443589 | - | EF013468 | EF013605 | KT443669 | KT443749 | EF013733 | EF013306 | EF012897 | EF013025 | |
| 77 | BLF8002 | USNM00756213 | LaPolla *et al.* 2010, LaPolla *et al*. 2012 | Ingroup | Formicinae | Lasiini | Nylanderia_amblyops | - | JN562404 | FJ982610 | FJ982567 | - | - | - | FJ982525 | FJ982487 | - | - | |
| 78 | ParaM19 | USNM00756215 | LaPolla *et al.* 2010, LaPolla *et al*. 2012 | Ingroup | Formicinae | Lasiini | Nylanderia_cf_madagascarensis | - | JN562406 | FJ982625 | FJ982581 | - | - | - | FJ982539 | FJ982501 | - | - | |
| 79 | ParaM32 | USNM00756218 | LaPolla *et al.* 2010, LaPolla *et al*. 2012 | Ingroup | Formicinae | Lasiini | Nylanderia_cf_vaga | - | JN562409 | FJ982633 | FJ982589 | - | - | - | FJ982547 | FJ982509 | - | - | |
| 57 | - | CASENT0059506 | LaPolla et al. 2010, Blaimer *et al.* 2015 | Ingroup | Formicinae | Lasiini | Nylanderia_dodo | KT443461 | KT443590 | FJ982617 | FJ982574 | KT443519 | KT443670 | KT443750 | FJ982532 | FJ982494 | KT443345 | KT443403 | |
| 72 | - | CASENT0106329 | Blaimer *et al.* 2015 | Ingroup | Formicinae | Lasiini | Nylanderia_emmae | KT443474 | KT443609 | - | KT443244 | KT443532 | KT443689 | KT443769 | KT443300 | KT443188 | KT443358 | KT443416 | |
| 58 | - | CASENT0106037 | Brady *et al.* 2006, LaPolla *et al.* 2010, Blaimer *et al.* 2015 | Ingroup | Formicinae | Lasiini | Nylanderia_hystrix | EF013162 | KT443591 | FJ982624 | EF013478 | EF013614 | KT443671 | KT443751 | EF013742 | EF013316 | EF012906 | EF013034 | |
| 81 | ParaM24 | USNM00756217 | LaPolla *et al.* 2010, LaPolla *et al*. 2012 | Ingroup | Formicinae | Lasiini | Nylanderia_phantasma | - | JN562408 | FJ982630 | FJ982586 | - | - | - | FJ982544 | FJ982506 | - | - | |
| 59 | ParaM4 | USNM00756214 | LaPolla *et al.* 2010, LaPolla *et al*. 2012 | Ingroup | Formicinae | Lasiini | Nylanderia_sp_GUY03 | - | JN562405 | FJ982623 | FJ982580 | - | - | - | FJ982538 | FJ982500 | - | - | |
| 60 | - | CASENT0049168 | LaPolla *et al.* 2010, Blaimer *et al.* 2015 | Ingroup | Formicinae | Lasiini | Nylanderia_sp_MG01 | KT443462 | KT443592 | FJ982626 | FJ982582 | KT443520 | KT443672 | KT443752 | FJ982540 | FJ982502 | KT443346 | KT443404 | |
| 84 | ParaM8 | USNM00756216 | LaPolla *et al.* 2010, LaPolla *et al*. 2012 | Ingroup | Formicinae | Lasiini | Nylanderia_sp_PER01 | - | JN562407 | FJ982628 | FJ982584 | - | - | - | FJ982542 | FJ982504 | - | - | |
| 85 | ParaM34 | USNM00756219 | LaPolla *et al.* 2010, LaPolla *et al*. 2012 | Ingroup | Formicinae | Lasiini | Nylanderia_vitiensis | - | JN562410 | FJ982634 | FJ982590 | - | - | - | FJ982548 | FJ982510 | - | - | |
| 86 | ParaM25 | USNM00756220 | LaPolla *et al.* 2010, LaPolla *et al*. 2012 | Ingroup | Formicinae | Lasiini | Nylanderia_vividula | - | JN562411 | FJ982635 | FJ982591 | - | - | - | FJ982549 | FJ982511 | - | - | |
| 87 | | CASENT0106331 | Blaimer *et al.* 2015 | Formicine outgroup | Formicinae | Oecophyllini | Oecophylla_longinoda | KT443463 | KT443593 | - | KT443233 | KT443521 | KT443673 | KT443753 | KT443289 | KT443177 | KT443347 | KT443405 | |
| 61 | - | CASENT0067057 | Blaimer *et al.* 2015 | Ingroup | Formicinae | Lasiini | Paraparatrechina_glabra | KT443465 | KT443597 | FJ982638 | KT443235 | KT443523 | KT443677 | KT443757 | KT443291 | KT443179 | KT443349 | KT443407 | |
| 62 | - | CASENT0106330 | Blaimer *et al.* 2015 | Ingroup | Formicinae | Lasiini | Paraparatrechina_oceanica | KT443466 | KT443598 | FJ982639 | KT443236 | KT443524 | KT443678 | KT443758 | KT443292 | KT443180 | KT443350 | KT443408 | |
| 63 | - | CASENT0914212 | Blaimer *et al.* 2015 | Ingroup | Formicinae | Lasiini | Paratrechina_antsingy | KT443467 | KT443599 | - | KT443237 | KT443525 | KT443679 | KT443759 | KT443293 | KT443181 | KT443351 | KT443409 | |
| 64 | - | CASENT0106255 | Blaimer *et al.* 2015 | Ingroup | Formicinae | Lasiini | Paratrechina_longicornis | KT443468 | KT443600 | FJ982644 | KT443238 | KT443526 | KT443680 | KT443760 | KT443294 | KT443182 | KT443352 | KT443410 | |
| 65 | - | CASENT0280583 | Blaimer *et al.* 2015 | Ingroup | Formicinae | Lasiini | Paratrechina_zanjensis | KT443469 | KT443601 | - | KT443239 | KT443527 | KT443681 | KT443761 | KT443295 | KT443183 | KT443353 | KT443411 | |
| 66 | - | CASENT0217048 | Blaimer *et al.* 2015 | Formicine outgroup | Formicinae | Plagiolepidini | Petalomyrmex_phylax | KT443470 | KT443602 | - | KT443240 | KT443528 | KT443682 | KT443762 | KT443296 | KT443184 | KT443354 | KT443412 | |
| 67 | | CASENT0106117 | Brady *et al.* 2006, Ward *et al.* 2015 | Non-formicine outgroup | Myrmicinae | Attini | Pheidole_clydei | EF013163 | KJ861055 | KJ860820 | EF013479 | EF013615 | KJ861662 | KJ860391 | EF013743 | EF013317 | EF012907 | EF013035 | |
| 130 | - | CASENT0077847 | Blaimer *et al.* 2015 | Formicine outgroup | Formicinae | Plagiolepidini | Plagiolepis_alluaudi | KT443472 | KT443604 | - | KT443242 | KT443530 | KT443684 | KT443764 | KT443298 | KT443186 | KT443356 | KT443414 | |
| 68 | - | CASENT0107071 | Blaimer *et al.* 2015 | Formicine outgroup | Formicinae | Plagiolepidini | Plagiolepis_sp_MG05 | KT443473 | KT443605 | - | KT443243 | KT443531 | KT443685 | KT443765 | KT443299 | KT443187 | KT443357 | KT443415 | |
| 69 | | CASENT0106213 | Ward *et al.* 2015 | Non-formicine outgroup | Myrmicinae | Pogonomyrmecini | Pogonomyrmex_vermiculatus | KJ861271 | KJ861069 | KJ860834 | KJ859808 | KJ861456 | KJ861676 | KJ860405 | KJ861884 | KJ860613 | KJ860184 | KJ859996 | |
| 126 | - | CASENT0106048 | Brady *et al.* 2006, Blaimer *et al.* 2015 | Formicine outgroup | Formicinae | Formicini | Polyergus_breviceps | EF013171 | KT443606 | - | EF013488 | EF013623 | KT443686 | KT443766 | EF013751 | EF013326 | EF012915 | EF013043 | |
| 70 | - | CASENT0106014 | Brady *et al.* 2006, Blaimer *et al.* 2015 | Formicine outgroup | Formicinae | Camponotini | Polyrhachis_decumbens | EF013172 | KT443607 | - | EF013489 | EF013624 | KT443687 | KT443767 | EF013752 | EF013327 | EF012916 | EF013044 | |
| 71 | - | CASENT0106007 | Brady *et al.* 2006, Blaimer *et al.* 2015 | Formicine outgroup | Formicinae | Camponotini | Polyrhachis_sp_Hagio01 | EF013173 | KT443608 | - | EF013490 | EF013625 | KT443688 | KT443768 | EF013753 | EF013328 | EF012917 | EF013045 | |
| 73 | D299 | CASENT0106035 | Brady *et al.* 2006, LaPolla *et al.* 2010, Blaimer *et al.* 2015 | Ingroup | Formicinae | Lasiini | Prenolepis_imparis | EF013175 | KT443610 | FJ982647 | EF013492 | EF013627 | KT443690 | KT443770 | EF013755 | EF013330 | EF012919 | EF013047 | |
| 101 | - | USNM00756231 | LaPolla *et al*. 2012 | Ingroup | Formicinae | Lasiini | Prenolepis_melanogaster | - | JN562421 | JN562432 | JN562458 | - | - | - | JN562465 | JN562449 | - | - | |
| 102 | ParaM21 | - | LaPolla *et al.* 2010, LaPolla *et al*. 2012 | Ingroup | Formicinae | Lasiini | Prenolepis_nitens | - | JN562422 | FJ982649 | FJ982602 | - | - | - | FJ982560 | FJ982519 | - | - | |
| 103 | - | USNM00756234 | LaPolla *et al*. 2012 | Ingroup | Formicinae | Lasiini | Prenolepis_sp_PHL02 | - | JN562423 | JN562433 | JN562459 | - | - | - | JN562466 | JN562450 | - | - | Most of AK missing. |
| 74 | - | CASENT0106258 | Blaimer *et al.* 2015 | Formicine outgroup | Formicinae | Formicini | Proformica_mongolica | KT443475 | KT443611 | - | KT443245 | KT443533 | KT443691 | KT443771 | KT443301 | KT443189 | KT443359 | KT443417 | |
| 75 | - | CASENT0227589 | Blaimer *et al.* 2015 | Formicine outgroup | Formicinae | Melophorini | Prolasius_convexus | KT443476 | KT443612 | - | KT443246 | KT443534 | KT443692 | KT443772 | KT443302 | KT443190 | KT443360 | KT443418 | |
| 76 | - | CASENT0106005 | Brady *et al.* 2006, LaPolla *et al.* 2010, Blaimer *et al.* 2015 | Ingroup | Formicinae | Lasiini | Pseudolasius_australis | EF013183 | KT443613 | FJ982651 | EF013501 | EF013635 | KT443693 | KT443773 | EF013763 | EF013339 | EF012927 | EF013055 | |
| 107 | ParaM107 | USNM00756237 | LaPolla *et al.* 2010, LaPolla *et al*. 2012 | Ingroup | Formicinae | Lasiini | Pseudolasius_sp_PHL01 | - | JN562398 | FJ982652 | FJ982603 | - | - | - | FJ982562 | JN562453 | - | - | |
| 108 | - | USNM00756238 | LaPolla *et al.* 2010, LaPolla *et al*. 2012 | Ingroup | Formicinae | Lasiini | Pseudolasius_typhlops | - | JN562426 | FJ982653 | FJ982604 | - | - | - | FJ982563 | JN562454 | - | - | |
| 125 | | CASENT0220669 | Chomicki *et al.* 2015 | Non-formicine outgroup | Pseudomyrmecinae | Pseudomyrmecini | Pseudomyrmex_osurus | KR829239 | KR829836 | KR830251 | KR829630 | KR829417 | KR830667 | KR830458 | KR829044 | - | - | KR828866 | |
| 77 | - | CASENT0227592 | Blaimer *et al.* 2015 | Formicine outgroup | Formicinae | Melophorini | Pseudonotoncus_hirsutus | KT443477 | KT443614 | - | KT443247 | KT443535 | KT443694 | KT443774 | KT443303 | KT443191 | KT443361 | KT443419 | |
| 78 | - | CASENT0106000 | Brady et al. 2006, LaPolla et al. 2010, Brady et al. 2014 | Non-formicine outgroup | Ectatomminae | Ectatommini | Rhytidoponera_chalybaea | EF013186 | FJ939885 | FJ939939 | EF013505 | EF013638 | KJ523705 | KJ523622 | EF013766 | EF013343 | EF012930 | EF013058 | |
| 129 | | CASENT0106171 | Lucky & Sarnat 2010, Ward *et al.* 2015 | Non-formicine outgroup | Myrmicinae | Solenopsidini | Rogeria_stigmatica | KJ861281 | GU085792 | KJ860846 | KJ859818 | GU075845 | KJ861688 | KJ860417 | KJ861894 | KJ860623 | KJ860194 | KJ860006 | |
| 79 | - | CASENT0235254 | Blaimer *et al.* 2015 | Formicine outgroup | Formicinae | Formicini | Rossomyrmex_anatolicus | KT443478 | KT443615 | - | KT443248 | KT443536 | KT443695 | KT443775 | KT443304 | KT443192 | KT443362 | KT443420 | |
| 80 | - | CASENT0004701 | Blaimer *et al.* 2015 | Formicine outgroup | Formicinae | Santschiellini | Santschiella_kohli | KT443479 | KT443616 | - | KT443249 | KT443537 | KT443696 | KT443776 | KT443305 | KT443193 | KT443363 | KT443421 | |
| 121 | | CASENT0139634 | Brady *et al.* 2014 | Non-formicine outgroup | Dorylinae | | Simopone_oculata | KJ523878 | KJ523360 | KJ523436 | KJ523823 | KJ523768 | KJ523710 | KJ523627 | KJ523547 | KJ523492 | KJ523286 | KJ523231 | |
| 81 | - | CASENT0106256 | Blaimer *et al.* 2015 | Formicine outgroup | Formicinae | Melophorini | Stigmacros_cf_clivispina | KT443480 | KT443617 | - | KT443250 | KT443538 | KT443697 | KT443777 | KT443306 | KT443194 | KT443364 | KT443422 | |
| 122 | | CASENT0491199 | Brady *et al.* 2014 | Non-formicine outgroup | Dorylinae | | Tanipone_zona | KJ523883 | KJ523367 | KJ523442 | KJ523828 | KJ523773 | KJ523717 | KJ523634 | KJ523552 | KJ523497 | KJ523291 | KJ523236 | |
| 82 | - | CASENT0041468 | Blaimer *et al.* 2015 | Formicine outgroup | Formicinae | Plagiolepidini | Tapinolepis_sp_MG01 | KT443481 | KT443618 | - | KT443251 | KT443539 | KT443698 | KT443778 | KT443307 | KT443195 | KT443365 | KT443423 | |
| 83 | - | CASENT0217038 | Blaimer *et al.* 2015 | Formicine outgroup | Formicinae | Plagiolepidini | Tapinolepis_sp_ZA01 | KT443482 | KT443619 | - | KT443252 | KT443540 | KT443699 | KT443779 | KT443308 | KT443196 | KT443366 | KT443424 | |
| 128 | | CASENT0106298 | Ward *et al.* 2015 | Non-formicine outgroup | Myrmicinae | Crematogastrini | Terataner_bottegoi | KJ861316 | KJ861119 | KJ860887 | KJ859853 | KJ861500 | KJ861729 | KJ860458 | KJ861929 | KJ860658 | KJ860229 | KJ860041 | |
| 84 | - | QMT162820 | Blaimer *et al.* 2015 | Formicine outgroup | Formicinae | Melophorini | Teratomyrmex_greavesi | KT443485 | KT443622 | - | KT443255 | KT443543 | KT443702 | KT443782 | KT443311 | KT443199 | KT443369 | KT443427 | |
| 85 | - | CASENT0106099 | Ward & Downie 2005, Ward *et al.* 2005, Brady *et al.* 2006, Brady *et al.* 2014 | Non-formicine outgroup | Pseudomyrmecinae | Pseudomyrmecini | Tetraponera_rufonigra | AY703716 | FJ939892 | FJ940011 | EF013524 | AY703783 | KJ523718 | KJ523635 | AY703649 | EF013362 | AY703515 | AY703582 | |
| 127 | | CASENT0106051 | Brady *et al.* 2006, Branstetter 2012, Ward *et al.* 2015 | Non-formicine outgroup | Myrmicinae | Stenammini | Veromessor_andrei | EF013137 | KJ861003 | KJ860766 | EF013446 | EF013589 | JQ742780 | JQ742836 | JQ742892 | EF013284 | EF012881 | EF013009 | |
| 123 | | CASENT0172777 | Brady *et al.* 2014 | Non-formicine outgroup | Dorylinae | | Vicinopone_conciliatrix | KJ523884 | KJ523368 | KJ523443 | KJ523829 | KJ523774 | KJ523719 | KJ523636 | KJ523553 | KJ523498 | KJ523292 | KJ523237 | |
| 86 | - | CASENT0106052 | Brady *et al.* 2006, LaPolla *et al.* 2010, Blaimer *et al.* 2015 | Ingroup | Formicinae | Lasiini | Zatania_albimaculata | EF013174 | KT443623 | FJ982646 | EF013491 | EF013626 | KT443703 | KT443783 | EF013754 | EF013329 | EF012918 | EF013046 | |
| 118 | - | USNM00756239 | LaPolla *et al*. 2012 | Ingroup | Formicinae | Lasiini | Zatania_cisipa | - | JN562428 | JN562435 | JN562460 | - | - | - | JN562468 | JX426004 | - | - | |
| 119 | - | USNM00756241 | LaPolla *et al*. 2012 | Ingroup | Formicinae | Lasiini | Zatania_gloriosa | - | JN562430 | JN562436 | JN562462 | - | - | - | JN562470 | JX426002 | - | - | |
