## Supplementary material for "Phylogeny, evolution, and classification of the ant genus *Lasius*, the tribe Lasiini, and the subfamily Formicinae (Hymenoptera: Formicidae)": Table S3

| | subset | best model | # sites | partition names |
|---|---|---|---|---|
| **Lasiini_w_outgroups_135t** | 1 | K80+I | 1965 | F1_pos2,18S_96t |
| | 2 | GTR+G | 811 | 28S_97t |
| | 3 | GTR+I | 319 | AA_pos1,F1_pos1 |
| | 4 | HKY+I | 408 | AA_pos2,UB_pos2 |
| | 5 | SYM+G | 200 | AA_pos3 |
| | 6 | K80+G | 819 | CD_exon3_pos1,CD_exon1_pos1,CD_exon2_pos1,AK_exon1_pos1,TP_pos1,AK_exon2_pos1 |
| | 7 | SYM+G | 387 | LR_exon2_pos2,AK_exon1_pos2,LR_exon1_pos2,WG_pos2 |
| | 8 | SYM+G | 839 | LR_exon1_pos3,AK_exon2_pos3,AK_exon1_pos3,CD_exon2_pos3,F2_pos3,TP_pos3 |
| | 9 | SYM+I | 286 | F2_pos2,AK_exon2_pos2 |
| | 10 | HKY+G | 185 | AK_intron,CD_intron2,LR_intron |
| | 11 | HKY+G | 597 | CD_exon3_pos2,CD_exon1_pos2,TP_pos2,CD_exon2_pos2 |
| | 12 | K80+G | 474 | CD_exon1_pos3,CD_exon3_pos3,WG_pos3,F1_pos3 |
| | 13 | K80+G | 229 | LR_exon2_pos1,WG_pos1,CD_intron1 |
| | 14 | GTR+G | 171 | F2_pos1 |
| | 15 | SYM+G | 76 | LR_exon1_pos1 |
| | 16 | K80+G | 284 | LR_exon2_pos3,UB_pos3 |
| | 17 | K80+I | 208 | UB_pos1 |
| **Lasiini_55t** | 1 | JC+I | 2364 | UB_pos1,F2_pos2,18S_17t,F1_exon2_pos2 |
| | 2 | GTR+I | 2159 | CD_exon2_pos1,TP_pos1,LR_exon1_pos2,AK_exon1_pos1,AK_exon2_pos1,F1_exon1_pos1,28S_17t,AA_pos1,F2_pos1,F1_exon2_pos1 |
| | 3 | F81 | 424 | AA_pos2,UB_pos2 |
| | 4 | GTR+I | 675 | WG_pos3,F1_exon2_pos3,LR_exon1_pos3,F1_exon1_pos3,AA_pos3 |
| | 5 | HKY+I | 683 | CD_exon3_pos2,CD_exon2_pos2,TP_pos2,CD_exon1_pos2,AK_exon1_pos2 |
| | 6 | HKY+G | 222 | AK_exon2_pos3,AK_exon1_pos3 |
| | 7 | JC | 232 | F1_exon1_pos2,AK_exon2_pos2 |
| | 8 | HKY+I | 96 | AK_55t_intron |
| | 9 | K80+G | 638 | UB_pos3,LR_exon2_pos3,CD_50t_intron2,F1_35t_intron1,LR_exon1_pos1,CD_exon3_pos1,CD_exon1_pos1 |
| | 10 | HKY+G | 347 | CD_50t_intron1,CD_exon2_pos3,CD_exon1_pos3,LR_39t_intron |
| | 11 | K80+G | 526 | CD_exon3_pos3,TP_pos3,F2_pos3 |
| | 12 | K80+I | 418 | LR_exon2_pos2,LR_exon2_pos1,WG_pos1,WG_pos2 |
